# Fluoxetine treatment during postnatal and juvenile temporal epochs evokes diametrically opposing changes in anxio-depressive behaviors, gene expression, mitochondrial function, and neuronal architecture in the medial prefrontal cortex

**DOI:** 10.1101/2023.12.29.573529

**Authors:** Utkarsha Ghai, Parul Chachra, Sashaina E. Fanibunda, Suchith Mendon, Amogh Bhaskaran, Ambalika Sarkar, Kowshik Kukkemane, Vivek Singh, Vidita A. Vaidya

**Affiliations:** Department of Biological Sciences, Tata Institute of Fundamental Research, Mumbai 400005, India; Kasturba Health Society - Medical Research Centre, Mumbai 400056, India

**Author notes:** **Address correspondence to:** Vidita A. Vaidya, Ph.D. Department of Biological Sciences Tata Institute of Fundamental Research 1, Homi Bhabha Road Mumbai, 400005 India.

**Keywords:** selective serotonin reuptake inhibitor (SSRI), Prozac, antidepressant, adolescence, mitochondria, microarray, mTOR, dendrite, anxiety, despair

## Abstract

**Background:** The selective serotonin reuptake inhibitor, fluoxetine is reported to evoke distinct effects on anxio-depressive behaviors based on the temporal window of administration. Here, we systematically addressed the influence of postnatal or juvenile fluoxetine treatment on anxio-depressive behavior, gene expression, mitochondrial biogenesis, and neuronal cytoarchitecture in adulthood.

**Methods:** Rat pups received postnatal fluoxetine (PNFlx) or juvenile fluoxetine (JFlx) treatment from postnatal day 2 (P2)-P21 or P28-48 respectively, and were assessed for changes in anxio-depressive behaviors, global gene expression, mitochondrial biogenesis/function, and dendritic cytoarchitecture in the medial prefrontal cortex (mPFC) in adulthood.

**Results:** PNFlx evoked long-lasting increases in anxio-depressive behaviors, whereas JFlx elicited persistent decreases in anxio-depressive behavior, accompanied by differential and minimally overlapping transcriptional changes in the mPFC in adulthood. We noted opposing changes in mitochondrial function and dendritic cytoarchitecture in the mPFC of PNFlx and JFlx animals, with a decline observed following PNFlx and an increase in response to JFlx treatment. Furthermore, the enhanced despair-like behavior in the PNFlx cohort was reversed by adult-onset treatment with nicotinamide, a precursor for NAD^+^ which enhances mitochondrial bioenergetics.

**Conclusions:** Fluoxetine treatment in early postnatal versus juvenile windows evokes opposing and persistent effects on anxio-depressive behavior in adult male rats, along with differential effects on gene expression, mitochondrial function, and dendritic morphology in the mPFC. Collectively, our findings highlight two distinct temporal windows in which fluoxetine exposure programs starkly differing outcomes in mood-related behavior, and posits a role for altered bioenergetics within the mPFC in contributing to these distinctive changes in emotionality.

## Introduction

Selective serotonin reuptake inhibitors (SSRIs) are the first line of treatment for major depressive disorders (1,2). The SSRI, fluoxetine, with its relatively favourable risk-benefit profile is the most commonly prescribed antidepressant for gestational and postpartum depression, as well as childhood and adolescent depression (3,4). SSRIs can readily cross the placental barrier (5) and are secreted in breast milk (6) raising the possibility that they could impact neurodevelopmental outcomes and exert long-term effects on offspring. Preclinical studies from rodent models indicate that gestational, postnatal or juvenile exposure to fluoxetine can exert persistent consequences on the programming of anxio-depressive behaviors (7–10), in addition to impacting the maturation of limbic brain regions implicated in emotional processing (11–15).

Serotonin plays a vital role in the fine-tuning of emotional neurocircuitry during distinct developmental epochs (16,17). Preclinical research examining the impact of SSRI exposure during early life collectively suggests differential effects on the programming of adult anxio-depressive behaviors, with reports of enhanced, decreased or no change in anxio-depressive behaviors, based on both the temporal window of administration and methodological differences (7–10,18–24). Several reports indicate that both pharmacological (7,12) and genetic (25) approaches that enhance serotonin levels in the brain during critical postnatal windows result in the programming of enhanced anxio-depressive behavior in adulthood in rodent models, accompanied by molecular and cytoarchitectural changes in limbic brain regions such as the medial prefrontal cortex (mPFC) (7,8,19,26–30). In comparison, relatively fewer reports have explored the behavioral consequences of enhanced serotonergic neurotransmission in the juvenile temporal window, and studies indicate either a decline, no change or a differential impact on anxio-depressive behaviors (9,21–24). Given that fluoxetine is often the treatment of choice for perinatal, childhood and adolescent depression (3,31), it is important that preclinical studies carefully assess the long-term impact of early exposure to fluoxetine on the programming of adult emotionality, as well as the influence on limbic neurocircuits such as the mPFC, that regulate anxio-depressive behavior.

Here, we have addressed the persistent behavioral consequences of exposure to the SSRI, fluoxetine, in two distinct, non-overlapping time windows, namely, the postnatal window (P2-P21) (PNFlx) and the juvenile (P28-P48) (JFlx) window. We have also examined the impact on global gene expression in the mPFC, and more extensively examined the regulation of pathways associated with the modulation of bioenergetics and neuronal cytoarchitecture. Our studies indicate opposing changes in anxio-depressive behaviors in adult animals with a history of PNFlx or JFlx, accompanied by unique global gene expression changes in the mPFC, as well as a differing impact on bioenergetics and neuronal architecture within this brain region. These preclinical studies highlight the importance of the temporal window in determining the outcomes of treatment with the SSRI, fluoxetine, and reveal that fluoxetine administration in postnatal and juvenile temporal epochs can evoke starkly opposing molecular, cellular, and behavioral outcomes.

## Materials and Methods

### Animals

Male Sprague-Dawley rats bred in the Tata Institute of Fundamental Research (TIFR) animal facility and maintained on a 12-hour light-dark cycle with *ad libitum* access to food and water were used for all experiments. Experimental procedures followed the guidelines of the Committee for the Purpose of Control and Supervision of Experimental Animals (CPCSEA) and were approved by the TIFR Animal Ethics Committee (56/1999/CPCSEA).

### Drug treatments

Litters born to primiparous dams were randomly assigned to vehicle (PNVeh or JVeh) or fluoxetine treatment (PNFlx or JFlx) groups. For PNFlx treatment, pups received oral administration via a feeding needle, of fluoxetine (10 mg/kg, IPCA laboratories, India) or vehicle (5% sucrose solution) once daily from postnatal day 2 (P2)-P21. Pups were separated at P28, and once weaned, all pups were housed in same-sex sibling groups of 3-4 rats. For JFlx treatment, weaned juvenile males received oral administration of fluoxetine via a feeding needle (10 mg/kg) or vehicle (5% sucrose solution) once daily from postnatal P28 to P48. Animals were left undisturbed until adulthood other than routine animal house maintenance. For the nicotinamide (NAM) experiment, animals with a history of PNFlx were treated with NAM (100 mg/kg, Sigma-Aldrich, USA) in drinking water at three months of age for thirty days. Drinking water was replaced every alternate day, and water consumption was measured and recorded, and did not differ across treatment groups (data not shown).

### Behavioral tests: Open Field, Elevated Plus Maze, and Forced Swim test

Animals were subjected to behavioral tasks to assess anxiety-like and despair-like behaviors in adulthood at three months of age. The open field test (OFT) (PNVeh: n = 9, PNFlx: n = 8, JVeh: n = 10 and JFlx: n = 10) to assess anxiety-like behavior consisted of an arena (100 cm x 100 cm x 70 cm), which was placed in a dimly illuminated room. Animals were released in one corner of the arena, and exploratory behavior was recorded with an automated tracking system (Ethovision, Noldus, Wageningen, Netherlands) for a duration of five minutes. The total distance traversed within the arena, path length, and time spent in the center of the arena were assessed. Animals were examined for anxiety-like behavior using the elevated plus maze (EPM) (PNVeh: n = 6, PNFlx: n = 6, JVeh: n = 11 and JFlx: n = 13) test for assessing anxiety-like behavior consisted of a platform elevated 50 cm from the ground with two open and two closed arms (45 x 10 cm). Animals were released into the center facing the open arm, and behavioral measures were assessed for a duration of 5 minutes. Ethovision analysis was performed for the total distance covered within the maze, path length, and time spent in the open and closed arms in the maze. Animals were assessed for despair-like behavior on a modified version of the forced swim test (FST) (PNVeh: n = 5, PNFlx: n = 6, JVeh: n = 8 and JFlx: n = 6) which involved habituation on day one for fifteen minutes to a plexiglass cylinder (50 cm tall, 21 cm in diameter) filled with water, followed by testing on day two wherein animals were placed in the FST tank for a duration of six minutes and behavior was assessed for the penultimate five minutes. Video recordings were assessed for immobility time and latency to immobility by an investigator blind to the treatment groups. Treatment groups were: postnatal vehicle (PNVeh) and postnatal fluoxetine (PNFlx) or juvenile vehicle (JVeh) and juvenile fluoxetine (JFlx). Behavioral testing on the open field test (OFT) (PNVeh: n = 9, PNFlx: n = 8, JVeh: n = 10 and JFlx: n = 8), elevated plus maze (EPM) (PNVeh: n = 6, PNFlx: n = 6, JVeh: n = 14 and JFlx: n = 13), and forced swim test (FST) was performed in adulthood at three months of age (PNVeh: n = 6, PNFlx: n = 6 and JVeh: n = 14, JFlx: n = 10).

To further examine the long-term influence of PNFlx and JFlx treatment, an independent cohort of animals were tested on the OFT, EPM, and FST at six months of age (OFT: PNVeh: n = 9, PNFlx: n = 8, JVeh: n = 10 and JFlx: n = 8; EPM: PNVeh: n = 9, PNFlx: n = 7, JVeh: n = 14 and JFlx: n = 13 and FST: PNVeh: n = 6, PNFlx: n = 6 and JVeh: n = 14, JFlx: n = 10). To assess the impact of NAM treatment, three-month-old animals with a history of PNFlx (n = 12/group) were assessed prior and post the NAM treatment for anxiety-like behavior on OFT. An independent cohort of three-month-old PNVeh (PNVeh + Vehicle: n = 12, PNVeh + NAM: n = 9) and PNFlx (PNFlx + Vehicle: n = 12, PNFlx + NAM: n = 8) animals were administered drinking water with or without NAM for thirty days and were tested for despair-like behavior on the FST.

### Microarray

Long-lasting global transcriptional changes within the mPFC of PNFlx or JFlx animals were assessed using microarray analysis as described previously (10). Three-month-old animals from the PNFlx (PNVeh (n = 4), PNFlx (n = 4)) and JFlx (JVeh (n = 4), JFlx (n = 4)) treated cohorts were rapidly decapitated, and bilateral mPFC tissue was dissected out. The mPFC tissue was snap-frozen in liquid nitrogen and stored at -80°C. RNA was extracted from the mPFC of six-month-old PNVeh and PNFlx or JVeh and JFlx animals using the Qiagen RNeasy mini kit. Total RNA integrity and purity was assessed using RNA 6000 Nano Lab Chip on the 2100 Bioanalyzer (Agilent, Palo Alto, CA, USA) and using the NanoDrop® ND-1000 UV-Vis Spectrophotometer (Nanodrop Technologies, Rockland, DE, USA) respectively. Total RNA samples with OD260/OD230≥1.3 and OD260/OD280 >1.8 were used for the microarray experiments. RNA samples with an RNA integrity number (RIN) ≥ 7.0, rRNA 28S/18S ratio greater than or equal to 1.5 were included in the array studies. To label the samples for gene expression, an Agilent Quick-Amp labelling Kit was used. PNVeh and PNFlx or JVeh and JFlx samples (500 ng each) were incubated with the reverse transcription mix and converted to double-stranded cDNA primed by oligo dT with a T7 polymerase promoter. cRNA was generated using double-stranded cDNA by *in vitro* transcription and the dye Cy3 was incorporated during this step. The cDNA synthesis *in vitro* transcription was carried out at 40°C. The labeled cRNA samples were hybridized to a Custom Rattus Norvegicus 8x15k with 15000 features, designed by Genotypic Technology Pvt. Ltd, India. The Gene Expression Hybridization kit of Agilent (Part Number 5188-5242) was used for fragmentation of labeled cRNA (600ng) and hybridization, which was carried out in Agilent’s Surehyb Chambers at 65° C for 16 hours. The Agilent Microarray Scanner G2505C at 5μm resolution was used to scan the hybridized slides after they were washed using Agilent Gene Expression wash buffers (Part No: 5188-5327). Significantly up-and down-regulated genes, showing 0.6-fold differences between the samples, were identified. Volcano Plots were used to calculate the *t*-test *p* value. Differentially regulated genes were clustered using hierarchical clustering to identify categories of significant gene expression patterns. Cut-offs greater than 1.35 for up-regulation and lesser than 0.74 for down-regulation were used to filter genes displaying differential regulation. Statistical analysis was performed using a *t*-test with a significance level of *p*<0.05 and was corrected for multiple comparisons using the Benjamini and Hochberg method (32). Based on fold change values, hierarchical clustering was performed for up- and down-regulated genes in mPFC using the Gene Spring software. Data was subjected to functional analysis using the DAVID (Database for Annotation, Visualization, and Integrated Discovery: http://david.abcc.ncifcrf.gov/) functional annotation tool.

### Quantitative PCR

To validate candidate genes from the microarray, we performed qPCR on tissue samples from an independent cohort of PNVeh (n = 8) and PNFlx (n = 7), and JVeh (n = 8) and JFlx (n = 8) animals. Tissue samples from mPFC were prepared as described above. RNA was extracted from the tissue using Tri-Reagent (Sigma, St. Louis, USA). Quality control was performed to check RNA integrity using a spectrophotometer (Nanodrop Technologies, Rockland, USA). RNA was reverse transcribed and subjected to qPCR using Sigma oligo-probes (Supplementary Table 1, 2). Quantification was done using the ^ΔΔ^Ct method, and the calibrator was chosen to be the samples of the vehicle group to which the samples of the treatment group were to be compared (33). The ^ΔΔ^Ct or calibrated value for each sample was given by the formula ^ΔΔ^Ct = ^Δ^Ct sample - ^Δ^Ct calibrator. The fold change for each sample was determined relative to the calibrator = 2 (-^ΔΔ^Ct). Data was normalized to the endogenous housekeeping gene hypoxanthine-guanine phosphoribosyltransferase 1 (*Hprt1*). Results are expressed expressed as fold change of vehicle ± SEM.

### Mitochondrial DNA levels

Mitochondrial DNA (mtDNA) levels were compared in the mPFC of PNVeh (n = 8) versus PNFlx (n = 8) or JVeh (n = 9) versus JFlx (n = 8) by quantitative real-time PCR (qPCR). Total DNA was extracted from cells using the commercially available NucleoSpin Triprep kit (Macherey-Nagel, Germany). To quantify levels of mtDNA, levels of cytochrome B - a mitochondrial genome-encoded gene, were normalized to levels of a nuclear-encoded gene, cytochrome C (Supplementary Table 3). Relative mitochondrial DNA levels between groups were quantified by the ^ΔΔ^Ct method described previously.

### Western Blot Analysis

For western blotting analysis, six-month-old animals from the PNFlx (PNVeh (n = 4), PNFlx (n = 4)) and JFlx (JVeh (n = 4), JFlx (n = 4)) experiments were rapidly decapitated and bilateral mPFC tissue was dissected out. The mPFC was lysed in RIPA buffer, and proteins in the samples were then separated on SDS-PAGE and transferred to a polyvinyl difluoride (PVDF) membrane. The PVDF membranes were blocked with 5% BSA and incubated with rabbit mTOR (1:1000, Cell Signaling Technologies, USA), rabbit p-mTOR (1:1000, Cell Signaling Technologies), rabbit ULK-1 (1:1000, Cell Signaling Technologies), rabbit p-ULK1 (1:1000, Cell Signaling Technologies), rabbit p-70 S6K (1:1000, Cell Signaling Technologies), rabbit phosphorylated p-70 S6K (1:250, Cell Signaling Technologies), rabbit actin (1:5000, ABclonal, USA), rabbit 4EBP (1:1000, Cell Signaling Technologies), rabbit p-4EBP1 (1:500, Cell Signaling Technologies), rabbit SIRT1 (1:1000 Millipore, USA), rabbit TFAM (1:1000 Abcam, USA), mouse ATP5A (1:1000, Abcam) and rabbit VDAC1 (1:1000 Abcam) followed by incubation with horseradish peroxidase-conjugated goat anti-rabbit or goat anti-mouse secondary antibody. The blots were developed using Super Signal West Pico Plus (Thermofisher, USA). Densitometric measurements were performed using Image-J software, and the levels for each protein were normalized to their respective actin protein loading controls. After normalization, the ratio of phosphorylated protein/total protein was calculated respectively for the mTOR signaling pathway proteins.

### ATP Assay

The ATP assay was performed on tissue samples from an independent cohort of PNVeh (n = 7) and PNFlx (n = 7) animals, and JVeh (n = 8) and JFlx (n = 8) animals. To examine the effects of NAM treatment, ATP assay was performed on the mPFC tissue samples derived from PNVeh + Vehicle (n = 12), PNFlx + Vehicle (n = 12), PNVeh + NAM (n = 9), PNFlx + NAM (n = 8). The mPFC tissue was lysed in boiling water, and the lysate was centrifuged at 12,000 rpm for 20 min at 4°C. The supernatant was collected in a fresh microcentrifuge tube. Cellular ATP levels were measured using the ATP bioluminescent kit (Sigma-Aldrich), and analyzed further using a luminometer (Berthold Technologies, Germany). The ATP level for each sample was normalized to the protein content of each sample using the BCA protein assay kit (Sigma-Aldrich). Results are expressed as fold change ± S.E.M. ***Golgi Staining***

For Golgi staining, six-month-old male Sprague Dawley rats were transcardially perfused with 0.9% saline, followed by Golgi staining using the FD Rapid Golgi Stain kit (FD NeuroTechnologies, USA). Each brain was cut into smaller chunks containing the entire mPFC region of the brain and was incubated in the impregnation solution in the dark for 21 days. After the impregnation step, the brain chunk was incubated in the staining solution for 3 days. Sections of 150 μm thickness were cut using a vibratome (Leica, Germany). This was followed by incubating sections with a freshly prepared staining solution as per manufacturer’s instructions for 10 min. After the washing step, the sections were mounted on super frost plus glass slides and were dehydrated using xylene and fixed with DPX mountant medium (Sigma-Aldrich). Slides were coded, and neurons were traced by an experimenter blind to the treatment groups. Tracing of the layer ⅔ neurons of the infralimbic (IL) region of the mPFC was conducted at 20X magnification on the BX53 light microscope (Olympus, Japan) using the Neurolucida 10 system (MBF Biosciences, USA). The Neurolucida 10 Explorer was used for the Sholl analysis of the neuronal traces (PNVeh (n = 7), PNFlx (n = 6), JVeh (n = 8), JFlx (n = 8))

### Seahorse Analysis

Mitochondrial isolation was performed from the mPFC tissue of PNFlx (PNVeh (n = 3), PNFlx (n = 3)) and JFlx (JVeh (n = 3), JFlx (n = 3)) animals using a modified differential centrifugation protocol as described previously. The tissue was homogenized in ice-cold mitochondrial isolation buffer (MIB+ BSA, containing 210 mM mannitol, 70 mM sucrose, 5 mM HEPES, 1 mM EGTA, and 0.5% (w/v) fatty acid-free BSA) using a glass dounce homogenizer. This was followed by centrifugation at 800g for 10 minutes at 4°C, and the supernatant was then centrifuged at 8,000g for 10 min at 4°C. The pellet was then resuspended in a small volume of ice-cold mitochondrial assay buffer (MAS+BSA, containing 220 mM mannitol, 70 mM sucrose, 10 mM KH_2_PO_4_, 5mM MgCl_2_, 2mM HEPES, 1mM EGTA and 0.5% (W/V) fatty acid-free BSA). The amount of protein loaded was estimated using BCA estimations. Isolated mitochondria (10μg) were plated (10μg/50μl) in Seahorse XFe24 micro-plates in MAS+BSA buffer, which was supplemented with substrates, pyruvate (10 mM), and malate (2 mM), and centrifuged at 2000g for 20 min at 4°C. After adding mitochondria to the microplates, 450 μl of the substrate containing MAS+BSA was added to each well. Metabolic analysis was performed using the Seahorse XFe24 Analyzer (Agilent Technologies, USA). At first, basal state-2 respiration (primarily Complex-I dependent), under limiting ADP (endogenous) and succinate (endogenous) concentrations, was measured. Then Complex II-dependent state 3 respiration was measured post-injection of 2 μM rotenone + 10 mM Succinate + 4 mM ADP. State-2 (Complex-I/-II dependent) and State-3 (Complex-II dependent) oxygen consumption rate (OCR) are represented as fold change ± SEM. Oligomycin-sensitive reduction in OCR was computed as the ATP production rate and is represented as fold change ± SEM. All substrates and inhibitors were obtained from Sigma-Aldrich.

### Statistical Analysis

Data were first assessed for normal distribution using the Kolmogorov-Smirnov test. Unpaired, two-tailed Student’s t-test was used for statistical analysis of two group experiments. Sholl analysis on the apical dendrites of neurons was statistically analyzed using two-way ANOVA, followed by Tukey post hoc analysis. For four group experiments, for pre-and post-treatment analysis of NAM treatment of PNFlx animals on OFT, two-way repeated measures ANOVA was used for statistical analysis. For four group experiments, with and without NAM treatment of PNFlx animals on FST and ATP analysis, two-way ANOVA analysis followed by a Tukey *post-hoc* comparison test was used for statistical analysis. Data are expressed as mean ± SEM. Statistical analyses were performed using Graph Pad Prism (version 8.0) and a value of *p* < 0.05 was considered statistically significant.

## Results

### Differential effects of postnatal and juvenile fluoxetine treatment on anxiety- and despair-like behavior in adulthood

We sought to examine the effects of a history of PNFlx (Figure 1A and Supplementary Figure 1A) or JFlx (Figure 1B and Supplementary Figure 1B) treatment on anxiety- and despair-like behavior in three-month-old adult animals. Anxiety-like behavior was assessed using the OFT (Figure 1C and Supplementary Figure 1C) and EPM (Figure 1H and Supplementary Figure 1F). PNFlx treatment enhanced anxiety-like behavior, as revealed by significant decreases in the percent distance traversed (Figure 1D) and the percent time spent in the center (Figure 1E) of the OFT arena. The total distance traversed (Supplementary Figure 1D) in the OFT arena did not vary between the PNVeh and PNFlx groups. In contrast, adult animals with a history of JFlx treatment exhibited a decline in anxiety-like behavior, with a significant increase observed in the percent distance traversed (Figure 1F) and the percent time spent in the center (Figure 1G) of the OFT arena. The total distance moved within the OFT arena did not differ between the JVeh and JFlx groups (Supplementary Figure 1E). PNFlx animals in adulthood also exhibited a significant increase in anxiety-like behavior on the EPM task, as determined by a significant decline in the percent distance traversed (Figure 1I) and the percent time spent in the open arms (Figure 1J) of the maze. The total distance traversed in the EPM maze did not differ across the PNVeh and PNFlx groups (Supplementary Figure 1G). In contrast, JFlx-treated animals showed a decrease in anxiety-like behavior on the EPM, with a significant increase noted in the percent distance traversed (Figure 1K) and the percent time spent in the open arms (Figure 1L), with no difference observed in the total distance traversed in the maze (Supplementary Figure 1H).

**Figure 1.**
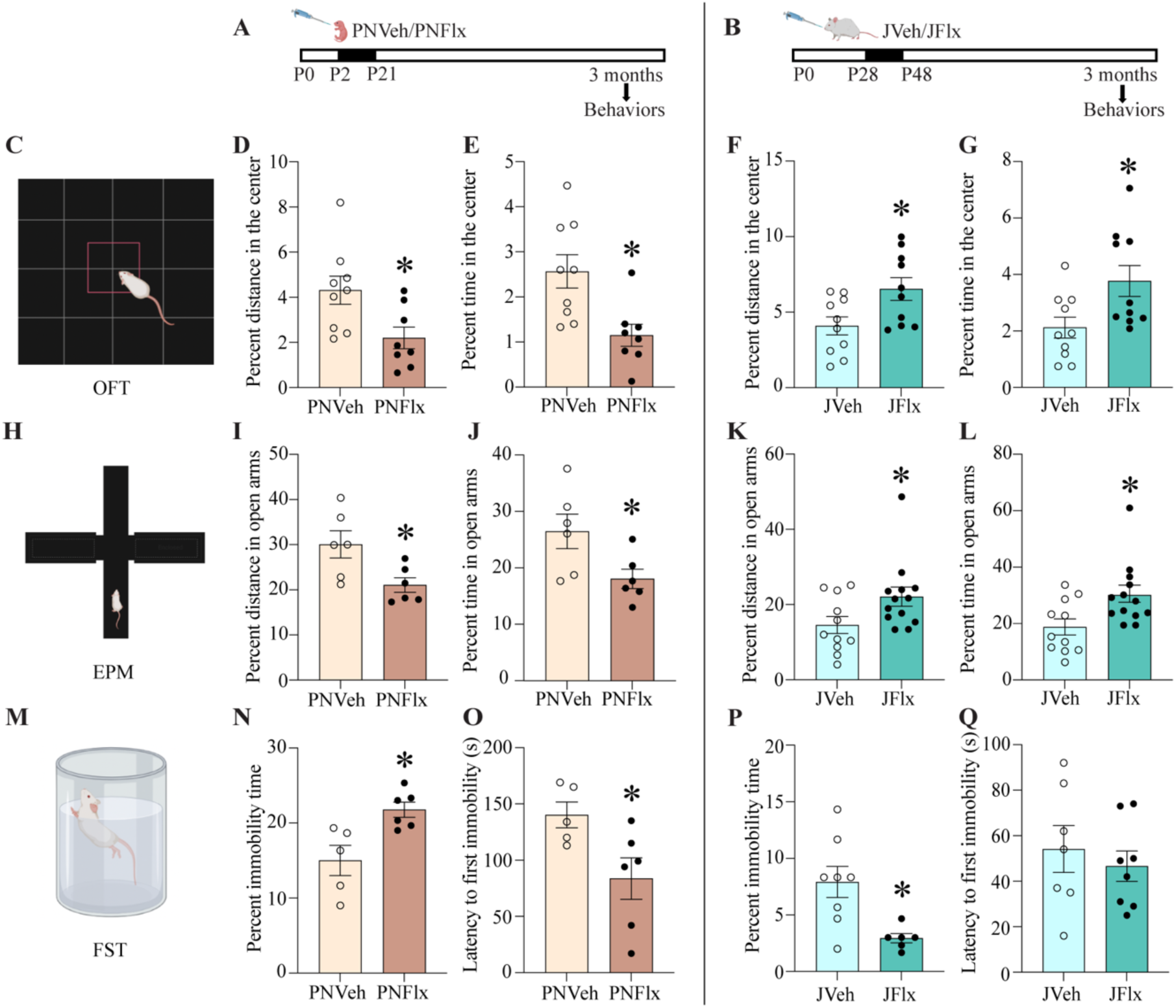
Postnatal and juvenile fluoxetine induce opposing effects on anxiety- and despair-like behavior in adulthood. Shown is a schematic representation of the treatment paradigm for postnatal fluoxetine (PNFlx) (A) from postnatal day 2 (P2) to P21 and juvenile fluoxetine (JFlx) (B) treatment from P28 to P48, with assessment for anxiety-like behavior at three months of age on the Open Field Test (OFT) (C) and Elevated Plus Maze (EPM) (H), and despair-like behavior on the Forced Swim Test (FST) (M). Shown is the percent distance traversed (D: PNFlx experiment; F: JFlx Experiment) and percent time (E: PNFlx experiment; G: JFlx Experiment) spent in the center of the OFT arena (PNVeh: n = 9, PNFlx: n = 8; JVeh: n = 10 and JFlx: n = 10). Shown is the percent distance traversed (I: PNFlx experiment; K: JFlx Experiment) and percent time spent (J: PNFlx experiment; L: JFlx Experiment)) in the open arms of the EPM (PNVeh: n = 6, PNFlx: n = 6; JVeh: n = 11 and JFlx: n = 13). Shown are the percent immobility time (N: PNFlx experiment; P: JFlx Experiment) and latency to first immobility (O: PNFlx experiment; Q: JFlx Experiment) on the FST (PNVeh: n = 5, PNFlx: n = 6; JVeh: n = 8 and JFlx: n = 6). Results are expressed as the mean ± SEM. \**p*<0.05 as compared to vehicle (Student’s *t*-test).

We next assessed whether three-month-old animals with a history of either PNFlx or JFlx treatment exhibited changes in despair-like behavior on the FST (Figure 1M). PNFlx treatment evoked an enhancement of despair-like behavior on the FST, as indicated by both a significant increase in percent immobility (Figure 1N) and decline in latency to immobility (Figure 1O). JFlx treatment evoked an opposing pattern of change with a significant decline noted in despair-like behavior on the FST, with a decrease in the percent immobility time in JFlx animals (Figure 1P) as compared to their age-matched, vehicle treated controls. The latency to first immobility (Figure 1Q) was unaltered across treatment groups in the JFlx experiment.

### Postnatal and juvenile fluoxetine treatment induce long-lasting and differential effects on anxiety- and despair-like behavior

We next addressed whether the opposing nature of behavioral changes noted in three month old animals with a history of PNFlx or JFlx treatment was persistent till six months of age. Six month old Sprague-Dawley male rats with a history of either PNFlx (Figure 2A and Supplementary Figure 2A) or JFlx (Figure 2B and Supplementary Figure 2B) treatment were subjected to behavioral analyses on the OFT (Figure 2C and Supplementary Figure 2C), EPM (Figure 2H and Supplementary Figure 2F), and FST (Figure 2M) to examine whether the effects on anxiety- and despair-like behaviors were long-lasting. Animals with a history of PNFlx treatment continued to exhibit a significant increase in anxiety-like behavior on the OFT, as evidenced by a decrease in the percent distance traversed (Figure 2D) and the percent time spent (Figure 2E) in the center of the OFT arena. Total distance traversed in the OFT arena did not differ across groups in the PNFlx experiment (Supplementary Figure 2D). Six month old animals with a prior history of JFlx treatment exhibited a decline in anxiety-like behavior on the OFT, with a significant increase in both percent distance (Figure 2F) and percent time spent (Figure 2G) in the center of the arena. JFlx treatment did not influence the total distance traversed in the OFT arena (Supplementary Figure 2E). In agreement with the results on the OFT, we also observed a persistent increase in anxiety-like behavior on the EPM in PNFlx animals, with a significant reduction in the percent distance traversed (Figure 2I) and the percent time spent (Figure 2J) in the open arms. JFlx animals at six months of age continued to exhibit a significant decline in anxiety-like behavior on the EPM, with an increase in both the percent distance traversed (Figure 2K) and the percent time spent (Figure 2L) in the open arms of the maze. The total distance traversed in the EPM maze did not differ across the groups in the PNFlx (Supplementary Figure 2G) and JFlx (Supplementary Figure 2H) experiments.

**Figure 2.**
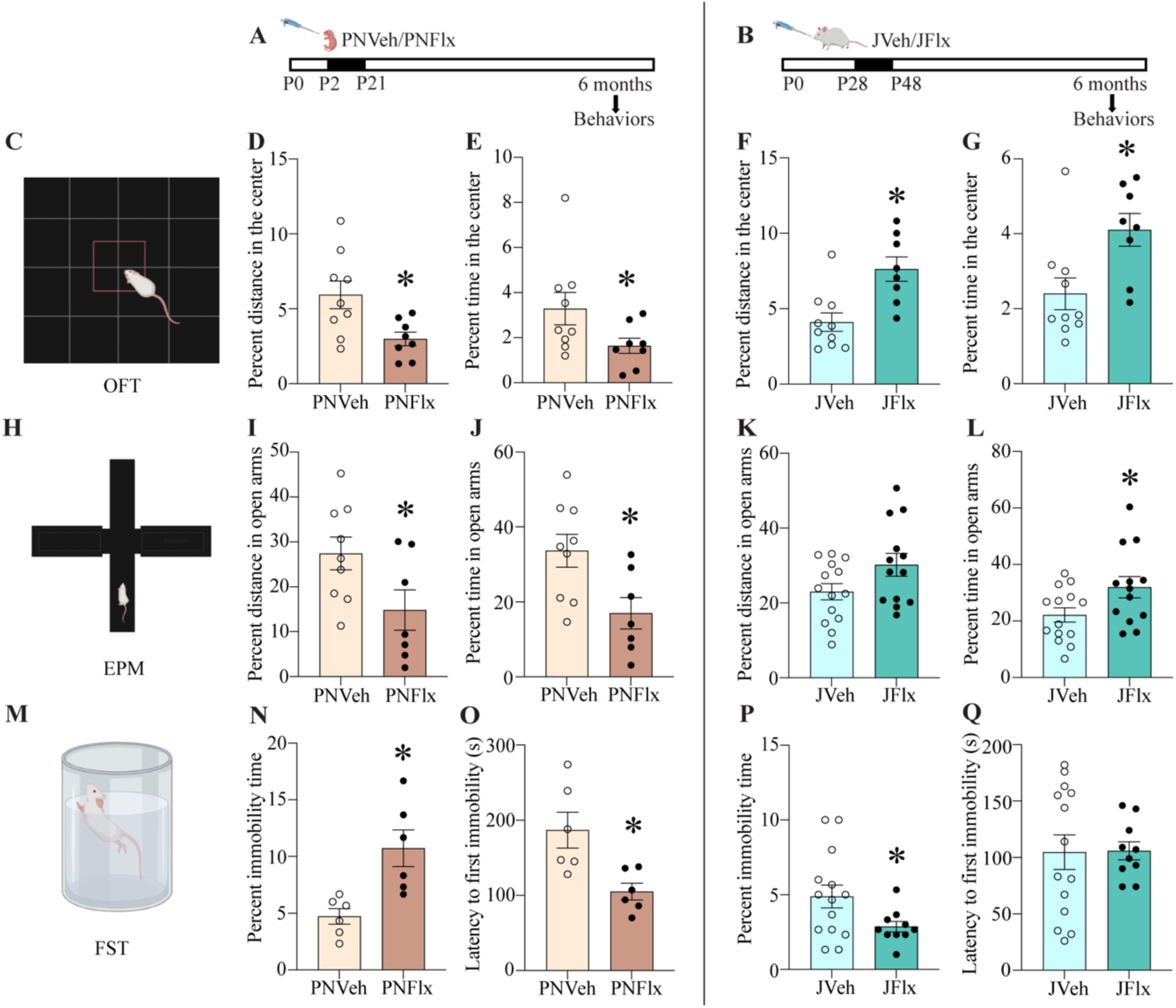
Postnatal and juvenile fluoxetine treatments evoke long-lasting and opposing effects on anxiety- and despair-like behavior. Shown is a schematic representation of the treatment paradigm for postnatal fluoxetine (PNFlx) (A) from postnatal day 2 (P2) to P21 and juvenile fluoxetine (JFlx) (B) treatment from P28 to P48, with assessment for anxiety-like behavior at six months of age on the Open Field Test (OFT) (C) and Elevated Plus Maze (EPM) (H), and despair-like behavior on the Forced Swim Test (FST) (M). Shown is the percent distance traversed (D: PNFLx experiment; F: JFlx Experiment) and percent time (E: PNFlx experiment; G: JFlx Experiment) spent in the center of the OFT arena (PNVeh: n = 9, PNFlx: n = 8; JVeh: n = 10 and JFlx: n = 8). Shown is the percent distance traversed (I: PNFlx experiment; K: JFlx Experiment) and percent time spent (J: PNFlx experiment; L: JFlx Experiment) in the open arms of the EPM (PNVeh: n = 9, PNFlx: n = 7; JVeh: n = 14, JFlx: n = 13). Shown are the percent immobility time (N: PNFlx experiment; P: JFlx Experiment)) and latency to first immobility (O: PNFlx experiment; Q: JFlx Experiment) on the FST (PNVeh: n = 6, PNFlx: n = 6; JVeh: n = 14, JFlx: n = 10). Results are expressed as the mean ± SEM. \**p*<0.05 as compared to vehicle (Student’s *t*-test).

PNFlx treatment evokes long-lasting increases in despair-like behavior on the FST (Figure 2M), with a significant increase in percent immobility (Figure 2N) and a decline in the latency to immobility (Figure 2O) in six-month-old PNFlx animals as compared to their age-matched, vehicle treated controls. The decline in despair-like behavior noted on the FST following JFlx treatment also persisted at six months of age, with significant reductions in percent immobility time (Figure 2P) noted in JFlx animals. The latency to first immobility (Figure 2Q) in the FST was unaltered across treatment groups in the JFlx experiment.

### Postnatal and juvenile fluoxetine treatments evoke a differential regulation of the mPFC transcriptome

To examine the nature of gene expression changes associated with the starkly opposing effects on anxiety- and despair-like behavior observed in animals following PNFlx or JFlx treatment, we performed microarray analysis on mPFC tissue derived from six month old animals with a history of PNFlx (Figure 3A) or JFlx (Figure 3B) treatment. Microarray analysis revealed a highly distinctive pattern of gene regulation within the mPFC of PNFlx (Figure 3C, Supplementary Table 4, 5) and JFlx (Figure 3F, Supplementary Table 6, 7) animals, with minimal overlap observed in gene regulation (Figure 3G). There was a larger number of genes significantly altered in their expression in the JFlx mPFC transcriptome (Figure 3F). Functional analysis with Database for Annotation, Visualization and Integrated Discovery (DAVID) of the PNFlx-regulated mPFC transcriptome (Figure 3D) revealed that amongst the genes downregulated in the PNFlx cohort there was a significant enrichment of functional categories associated with the postsynaptic membrane, dendritic spine, synapse, mitochondrion, AMP binding, ATP-sensitive potassium channel complex, transporter activity, and metabolic process. In contrast, functional analysis with DAVID of the JFlx-regulated mPFC transcriptome (Figure 3E) indicated that the genes upregulated following JFlx treatment showed significant enrichment of functional categories linked to synapse, synaptic transmission, ATP binding, regulation of synaptic transmission, mitochondrial outer membrane, postsynaptic density, and transporter activity. The functional categories significantly regulated in the PNFlx and JFlx mPFC transcriptomes are shown in Supplementary Table 8-11. The starkly differing transcriptomes in the mPFC of PNFlx versus JFlx animals was further indicated by the minimal overlap noted in gene regulation (Figure 3G). The genes that demonstrated a similar nature of regulation were 0.9% and 0.7% of the total transcriptome regulated in PNFlx and JFlx animals respectively (Figure 3G, Supplementary Table 12). Our microarray results reveal that treatment with the SSRI fluoxetine in distinct windows of postnatal life, namely in the early postnatal window and in adolescence, results in a largely non-overlapping, differential pattern of gene regulation in the mPFC.

**Figure 3.**
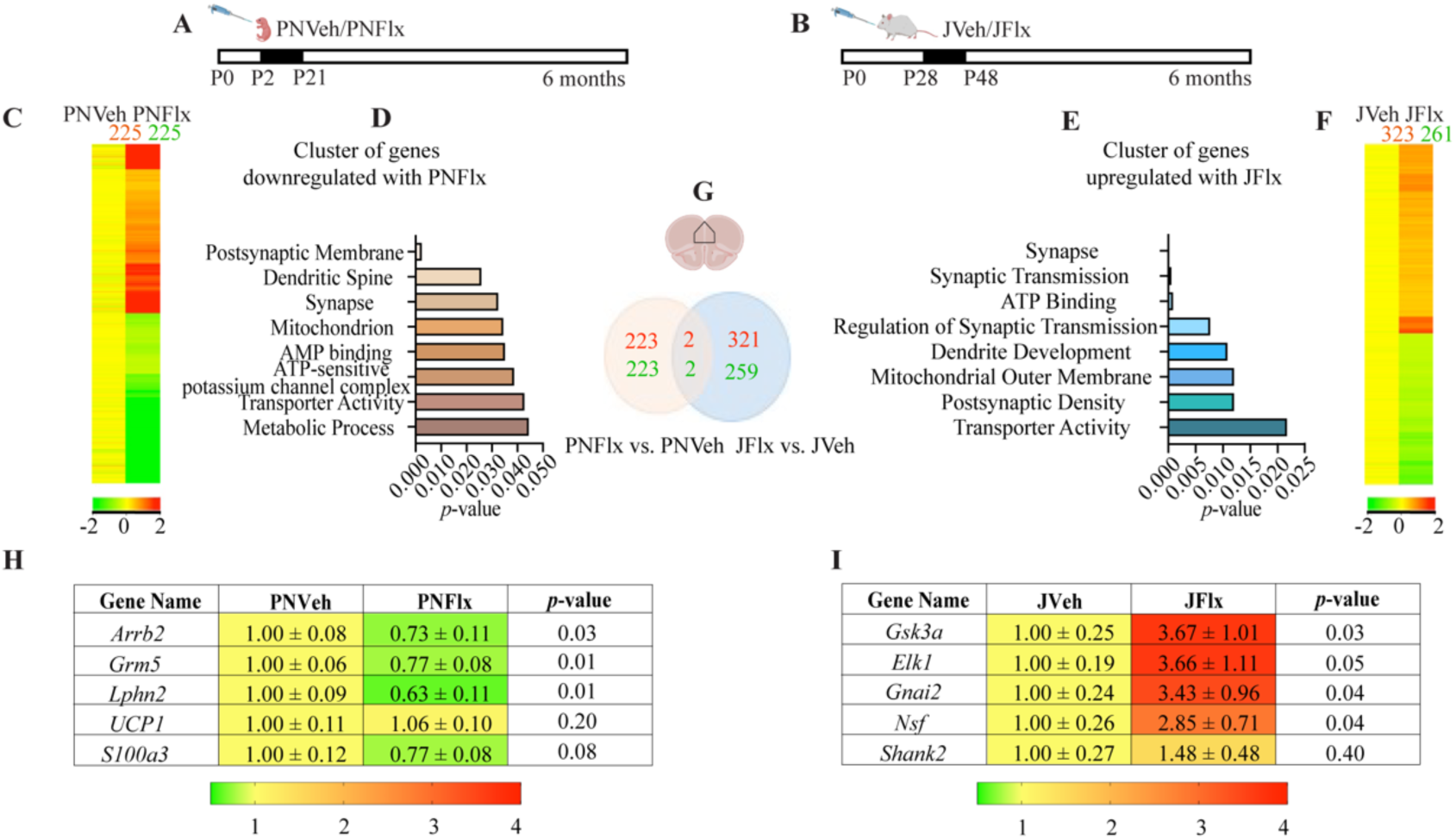
Postnatal fluoxetine and juvenile fluoxetine treatment differentially regulate gene expression in the mPFC. Shown is a schematic representation of the treatment paradigms for postnatal fluoxetine (PNFlx) (A) and juvenile fluoxetine (JFlx) (B) treatment. Microarray analysis was performed to assess the mPFC transcriptome of adult (6 months) (n = 4/group) animals with a history of PNFlx or JFlx as compared to their respective age-matched vehicle control groups. Shown are heat maps (C: PNFlx Experiment; F: JFlx Experiment) displaying the significantly regulated genes (*p* < 0.05, *t*-test corrected for multiple comparisons; upregulated genes are represented in red, and downregulated genes are represented in green). Functional analysis with Database for Annotation, Visualization and Integrated Discovery (DAVID) of the PNFlx mPFC transcriptome revealed a significant enrichment of the following functional categories in the downregulated genes, namely postsynaptic membrane, dendritic spine, synapse, mitochondrion, AMP binding, ATP-sensitive potassium channel complex, transporter activity, and metabolic process (D). DAVID analysis of the JFlx mPFC transcriptome indicated an enrichment of the following functional categories in the upregulated genes, namely synapse, synaptic transmission, ATP binding, regulation of synaptic transmission, dendritic development, mitochondrial outer membrane, postsynaptic density, and transporter activity (F). Analysis of the PNFlx and JFlx mPFC transcriptomes revealed distinct patterns of regulation with minimal overlap in regulation of gene expression (E). The number of upregulated (red) and downregulated (green) genes in PNFlx and JFlx animals compared to their respective age-matched vehicles are represented using Venn diagrams. Shown are changes in gene expression based on qPCR analyses performed for select genes using an independent cohort of animals for PNFlx (H) and JFlx (I) animals, which validated the regulation observed in the microarray. Results are expressed as fold change of vehicle and are the mean ± SEM. \**p*<0.05 as compared to vehicle (Student’s *t*-test).

We next sought to validate the microarray results using qPCR analysis to assess gene expression in the mPFC of an independent cohort of six month old animals with a history of PNFlx or JFlx treatment. We observed a regulation of select genes from our PNFlx transcriptome analysis (Figure 3H), with a significant downregulation observed for beta-arrestin 2 (*Arrb2), metabotropic glutamate receptor 5 (Grm5)*, and Latrophilin 2 *(Lphn2)* mRNA levels in the mPFC of PNFlx animals as compared to their controls. qPCR analysis performed for select genes belonging to distinct functional categories within our JFlx microarray (Figure 3I) confirmed a significant upregulation of glycogen synthase kinase 3 alpha (*Gsk3a),* ETS-like protein 1 (*Elk1), G protein subunit alpha i2 (Gnai2),* and N-Ethylmaleimide Sensitive Factor, Vesicle Fusing ATPase (*Nsf)* mRNA in the mpFC of JFlx animals. We did note that some of the genes regulated in our transcriptome analysis did not exhibit significant regulation in the qPCR analysis run on independent cohorts, namely uncoupling protein 1 (*UCP1)* and S100 calcium binding protein A3 (*S100a3*) in the PNFlx cohort and SH3 And Multiple Ankyrin Repeat Domains 2 (*Shank2*) in the JFlx cohort.

### Postnatal and juvenile fluoxetine treatments evoke opposing effects on mitochondrial gene and protein expression

The microarray data indicated a significant enrichment of functional gene categories such as mitochondrial pathway, metabolic process, nucleotide binding and transport associated genes in the PNFlx and JFlx transcriptomes. Given prior evidence that the neurotransmitter serotonin is a direct regulator of mitochondrial biogenesis and function in the neocortex (34), and that fluoxetine can be bound to mitochondrial fractions and impact mitochondrial function (35–37), we next addressed the influence of PNFlx (Figure 4A and Supplementary Figure 3A) and JFlx (Figure 4B and Supplementary Figure 3B) on mitochondrial biogenesis and function in the mPFC. We observed no significant change in mitochondrial DNA (mtDNA) levels in the mPFC of either the PNFlx (Figure 4C) or JFlx (Figure 4F) cohorts as compared to their respective age-matched controls. We next assessed the gene expression of master modulators of mitochondrial function, NAD-dependent deacetylase sirtuin-1 (*Sirt1*) and peroxisome proliferator-activated receptor gamma coactivator 1-alpha (*Ppargc1a*), mitochondrial transcription factor A (*Tfam*) and Cytochrome C (*Cycs*) in the mPFC of six month old animals with a history of either PNFlx or JFlx. We noted a significant decline in *Sirt1* mRNA levels in the mPFC of PNFlx animals (Figure 4D), but observed no significant change in the gene expression of *Tfam* (Figure 4E), *Ppargc1a* (Supplementary Figure 3C) and *Cycs* (Supplementary Figure 3D). In contrast, we observed a significant increase in the gene expression of *Sirt1* (Figure 4G) and *Tfam* (Figure 4H) while no change was observed in the gene expression of *Ppargc1a* (Supplementary Figure 3E) and *Cycs* (Supplementary Figure 3F) within the mPFC of the JFlx experimental cohort.

**Figure 4.**
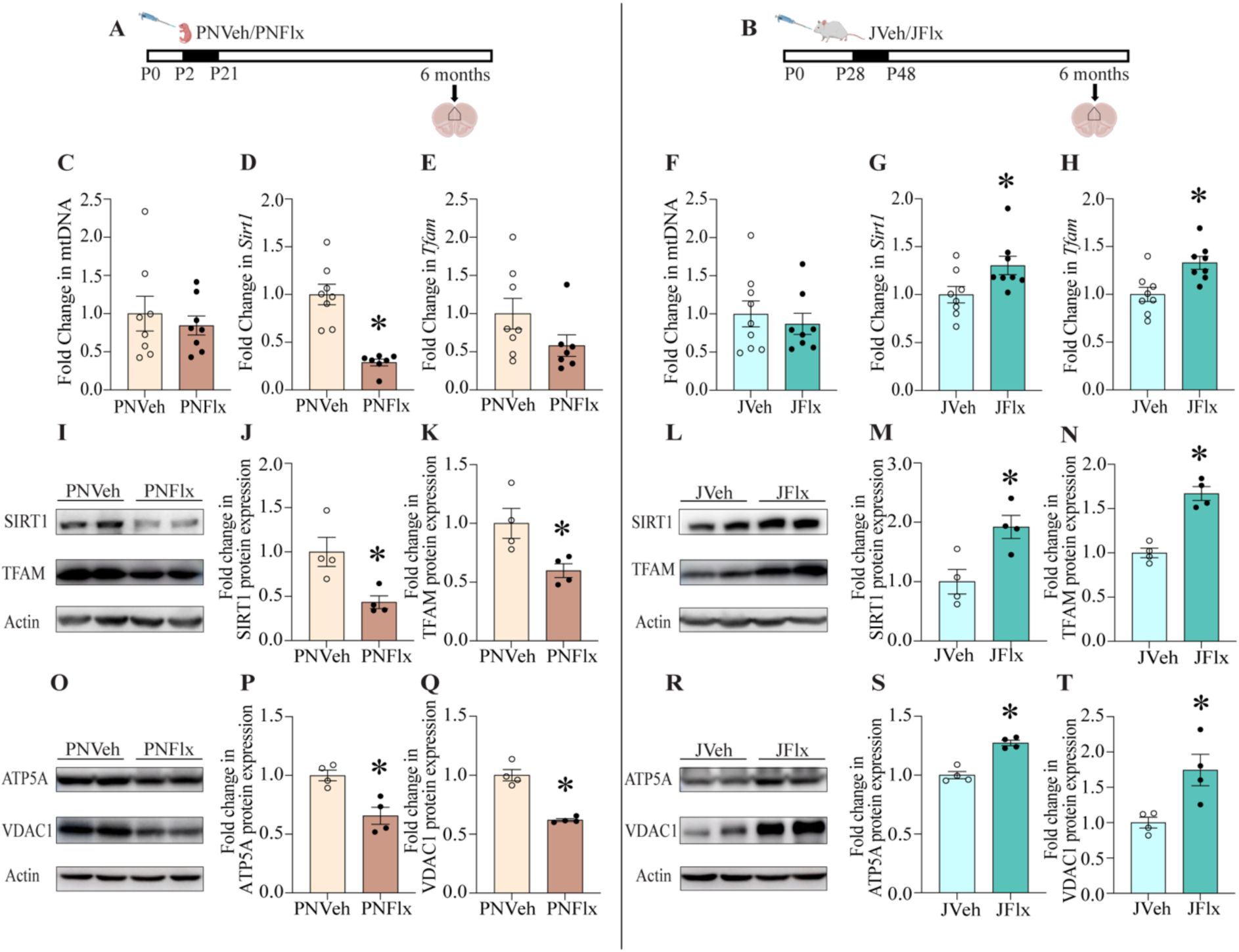
Postnatal and juvenile fluoxetine treatments evoke opposing effects on mitochondrial gene and protein expression. Shown is a schematic representation of the treatment paradigm for postnatal fluoxetine (PNFlx) (A) and juvenile fluoxetine (JFlx) (B) treatment. Shown is the quantification of fold change in mtDNA (C) levels (n = 8/group), gene expression levels of *Sirt1* (D) and *Tfam* (E) in the mPFC with PNFlx treatment (PNVeh: n = 8, PNFlx: n = 7). Shown is the quantification of fold change in mtDNA levels (F) (JVeh: n = 9, JFlx: n = 8) and gene expression levels of *Sirt1* (G) and *Tfam* (H) in the mPFC with JFlx treatment (n = 8/group). Shown are representative immunoblots for (I) SIRT1, TFAM, and actin for PNVeh and PNFlx-treated animals in adulthood. Shown are densitometric quantifications of levels of SIRT1 (J) and TFAM (K) protein expression in the mPFC normalized to actin expression in PNVeh and PNFlx-treated animals (n = 4/group). Shown are representative immunoblots for (L) SIRT1, TFAM, and actin for JVeh and JFlx-treated adult animals (n = 4/group). Shown are densitometric quantifications of levels of SIRT1 (M) and TFAM (N) protein expression in the mPFC normalized to actin levels in JVeh and JFlx-treated animals in adulthood (n = 4/group). Shown are representative immunoblots for (O) ATP5A, VDAC1, and actin in PNVeh and PNFlx-treated animals (n = 4/group). Shown are densitometric quantifications of levels of ATP5A (P) and VDAC1 (Q) protein expression in the mPFC normalized to actin levels for PNVeh and PNFlx-treated animals. Shown are representative immunoblots for (R) ATP5A, VDAC1, and actin in JVeh and JFlx treated animals. Shown are densitometric quantifications of levels of ATP5A (S), VDAC1(T) protein expression in the mPFC normalized to actinin JVeh and JFlx-treated animals in adulthood. Results are expressed as the fold change ± SEM. \**p*<0.05 compared to vehicle-treated (Student’s *t*-test).

We then assessed whether the expression of proteins implicated in the regulation of mitochondrial biogenesis and function, namely SIRT1, TFAM, ATP synthase F1 subunit alpha (ATP5A) and voltage-dependent anion channel 1 (VDAC1) are altered in animals with a history of PNFlx or JFlx treatment. Western blotting analysis revealed a significant decrease in SIRT1, TFAM, ATP5A, and VDAC1 (Figure 4I-K, 4O-Q) protein expression in the mPFC in the PNFlx group. In the JFlx treatment group, we observed a significant increase in SIRT1, TFAM, ATP5A, and VDAC1 (Figure 4L-N, 4R-T) expression in the mPFC. These results indicate that a history of fluoxetine treatment in the postnatal and juvenile window evokes a decrease or an increase in expression of multiple proteins associated with a regulation of mitochondrial biogenesis and function in the mPFC region in adulthood, respectively.

### Postnatal and juvenile fluoxetine treatments evokes opposing effects on mitochondrial bioenergetics

Given we observed a robust, and starkly opposing pattern of regulation of expression of proteins involved in mitochondrial biogenesis and function in the mPFC of PNFlx and JFlx animals, we addressed whether these changes are associated with functional changes in mitochondrial bioenergetics (Figure 5A-B and Supplementary Figure 4A-B). We examined cellular ATP levels which would reflect two primary production pathways, namely glycolysis and oxidative phosphorylation. We noted a significant decrease in cellular ATP levels within the mPFC of PNFlx animals (Figure 5C) and an increase in cellular ATP levels in the mPFC of JFlx animals (Figure 5E) as compared to their respective age-matched controls. We further also assessed for changes in cellular ATP levels in another limbic region involved in regulation of anxiety- and despair-like behavior, the hippocampus (HPC). We observed no significant differences in the cellular ATP levels in the HPC region with a history of PNFlx (Supplementary Figure 4C) or JFlx (Supplementary Figure 4D) treatment.

**Figure 5.**
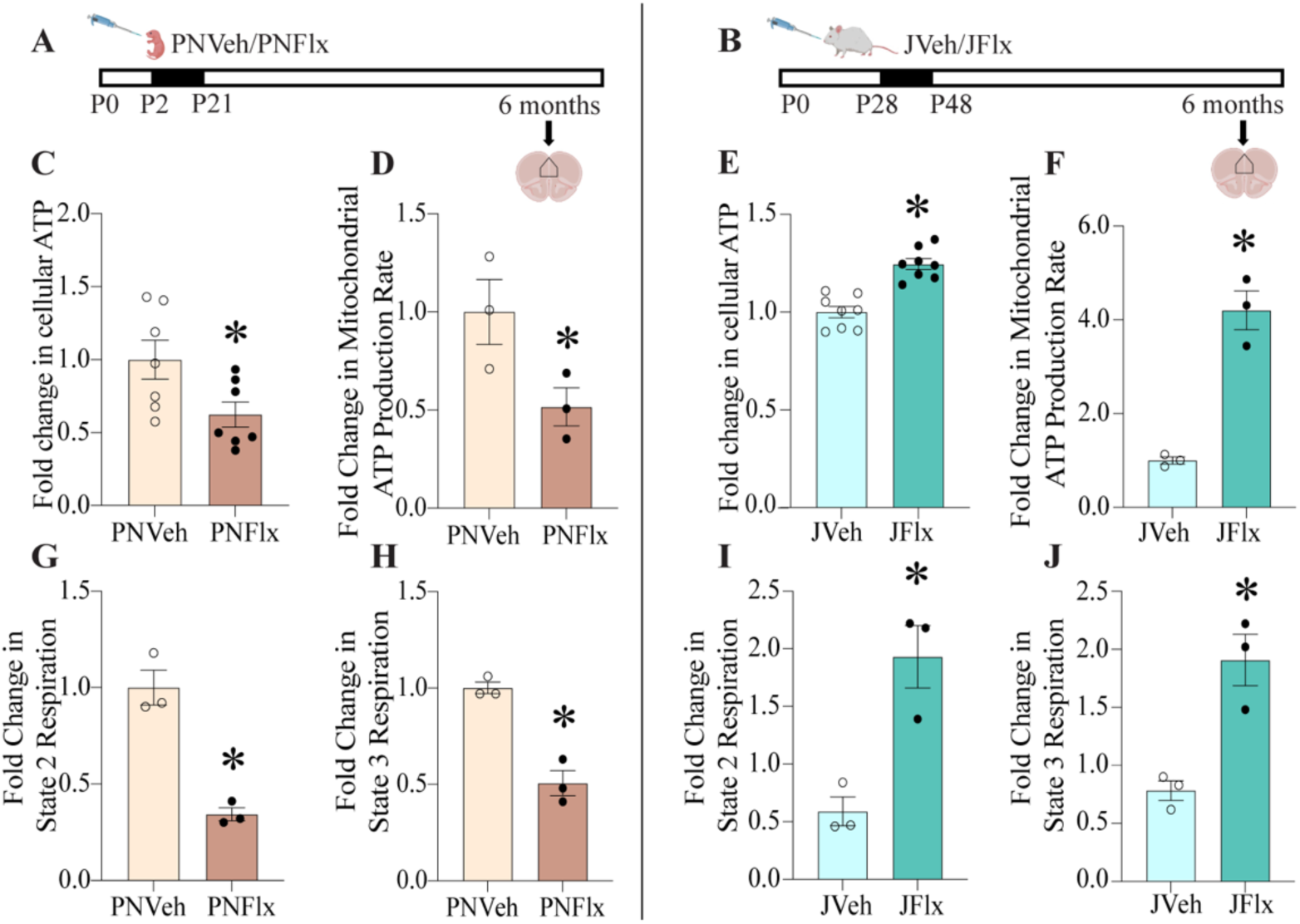
Postnatal and juvenile fluoxetine treatments evoke opposing effects on mitochondrial bioenergetics. Shown is a schematic representation of the treatment paradigms for postnatal (PNFlx) (A) and juvenile fluoxetine (JFlx) (B) treatment. Shown is the quantification of (C) cellular ATP levels in the mPFC of PNVeh and PNFlx-treated animals in adulthood (n = 7/group). Shown is the quantification of the Seahorse assay performed on mitochondria derived from the mPFC of PNVeh and PNFlx treated animals normalized to protein levels (D, G, H). The bar graph depicts the mitochondrial ATP production rate in the mPFC of PNVeh and PNFlx animals (D) (n = 3/group). Shown is the quantification of cellular ATP levels (E) in the mPFC of JVeh and JFlx-treated animals in adulthood (n = 8/group). Shown is the quantification of the Seahorse assay performed on mitochondria derived from the mPFC of JVeh and JFlx treated animals normalized to protein levels (F, I, J). The bar graph depicts the mitochondrial ATP production rate in the mPFC of JVeh and JFlx animals in adulthood (F) (n = 3/group). Shown is the quantitative analysis State 2 (G) and State 3(H) respiration in mitochondria derived from the mPFC of PNVeh and PNFlx animals (n = 3/group). Shown is the quantitative analysis of State 2 (I) and State 3 (J) respiration in mitochondria derived from the mPFC of JVeh and JFlx animals in adulthood (n = 3/group). Results are expressed as fold change ± SEM compared to age-matched vehicle controls. \**p*<0.05 compared to vehicle (Student’s *t*-test).

We performed Seahorse Analysis on isolated mitochondria derived from the mPFC of PNFlx and JFlx animals, and determined the mitochondrial ATP production rate using oligomycin to inhibit the ATP Synthase. We also measured state II respiration, which is primarily complex-I dependent and was calculated under limiting ADP and succinate concentrations, in the absence of inhibitors. State 3 respiration, which is primarily complex II-dependent respiration, calculated post-inhibition of complex I by rotenone, and under non-limiting concentrations of ADP and succinate was also determined. We noted a significant decrease in the mitochondrial ATP production rate (Figure 5D), State 2 (Figure 5G) and State 3 (Figure 5H) respiration in mitochondria derived from the mPFC of six month old PNFlx animals as compared to their age-matched controls. In stark contrast, six month old JFlx animals exhibited a significant increase in mitochondrial ATP production rate (Figure 5F), as well as enhanced State 2 (Figure 5I) and State 3 (Figure 5J) respiration. Given that, the changes in the state 2 and state 3 respiration, as well as the mitochondrial respiration rate, are calculated after normalization with mitochondrial protein content, our results suggests changes in the OxPhos efficiency with both PNFlx and JFlx treatment. These observations indicate that adult animals with a history of postnatal or juvenile treatment with the SSRI fluoxetine, exhibit starkly opposing outcomes on mitochondrial bioenergetics, with a decline following PNFlx and an increase with JFlx administration.

### Differential regulation of the mTOR signalling pathway within the mPFC of PNFlx and JFlx animals

The analysis of functional categories in our microarray indicated the enrichment of genes involved in the mammalian target of rapamycin (mTOR) signaling pathway, as well as genes associated with the regulation of dendritic development and spine architecture. Furthermore, the mTOR pathway besides being implicated in the modulation of dendritic morphology and spine dynamics, in particular in the context of antidepressant-evoked alterations in cytoarchitecture (10, 38–40), is also involved in the regulation of mitochondrial biogenesis and function (41–45). Hence, we next to sought to examine whether key mTOR signaling pathway components were regulated within the mPFC of six month old animals with a history of PNFlx (Figure 6A) or JFlx treatment (Figure 6B). We performed western blotting analysis to assess the status of phosphorylated and total protein levels of distinct components of the mTOR signaling pathway, including mTOR, and downstream effectors such as Unc-51-like autophagy-activating kinases 1 (Ulk1), p70 ribosomal S6 kinase (p70S6K), and eukaryotic translation initiation factor 4E-binding protein 1 (4EBP1) (Figure 6C-N). While PNFlx treatment did not alter p-mTOR/mTOR levels in the mPFC (Figure 6C, D, I), we noted significant decrease in the p-ULK1/ULK1 and p-p70S6K/p70S6K protein levels (Figure 4C, E, I, J) in the mPFC, suggestive of a decline in signaling via downstream effectors of the mTOR cascade. We did not observe any change in p-4EBP1/4EBP1 levels (Figure 6C, I, K) in the mPFC of PNFlx-treated animals. In contrast, JFlx treatment led to a significant increase p-mTOR/mTOR levels and p-4EBP1/4EBP1 levels in the mPFC (Figure 6F, G, L, N), however we observed no change in p-ULK1/ULK1 and p-p70S6K/p70S6K levels (Figure 6F, H, L, M) in the mPFC of animals with a history of JFlx. These findings suggest that PNFlx and JFlx treatments evoke a long-lasting and opposing pattern of regulation of the mTOR signaling cascade within the mPFC.

**Figure 6.**
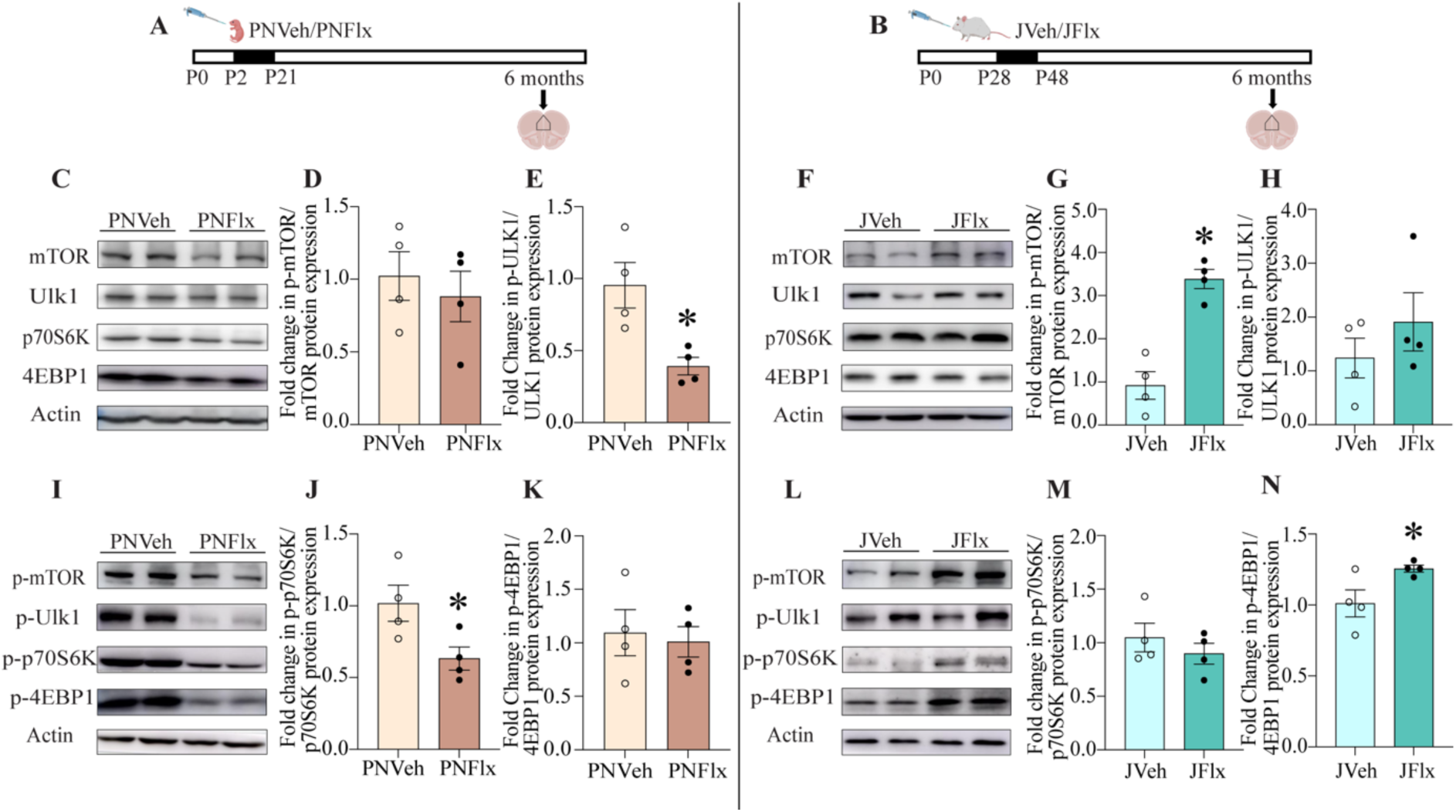
Postnatal fluoxetine and juvenile fluoxetine treatment evokes differential effects on the mTOR signalling pathway within the mPFC. Shown is a schematic representation of the treatment paradigms for postnatal fluoxetine (PNFlx) (A) and juvenile fluoxetine (JFlx) (B) treatment. Shown are representative immunoblots for (C) mTOR, ULK1, p70S6K, 4EBP1, and actin and for (I) phosphorylated-mTOR, phosphorylated-ULK1, phosphorylated-p70 S6K, phosphorylated-4EBP1 and actin from the mPFC of PNVeh and PNFlx-treated animals in adulthood. Shown are densitometric quantifications of phosphorylated-mTOR/ mTOR (D), phosphorylated-ULK1/ ULK1 (E), phosphorylated-p70 S6K/ p70 S6K (J), phosphorylated-4EBP1/ 4EBP1 (K) normalized to their respective beta-actin loading controls in the mPFC of PNVeh and PNFlx-treated animals. Shown are representative immunoblots for (F) mTOR, ULK1, p70S6K, 4EBP1, and actin and for (L) phosphorylated-mTOR, phosphorylated-ULK1, phosphorylated-p70 S6K, phosphorylated-4EBP1 and actin from the mPFC of JVeh and JFlx-treated animals in adulthood. Shown are densitometric quantifications of phosphorylated-mTOR/ mTOR (G), phosphorylated-ULK1/ ULK1 (H), phosphorylated-p70 S6K/ p70 S6K (M), phosphorylated-4EBP1/ 4EBP1 (N) normalized to their respective beta-actin loading controls in the mPFC of JVeh and JFlx-treated animals. Results are expressed as fold change of vehicle-treated controls and are the mean ± SEM. \**p*<0.05 compared to vehicle-treated controls (Student’s *t*-test, n = 4).

### Postnatal Fluoxetine and Juvenile Fluoxetine treatment evoke opposing effects on neuronal arborization within the mPFC

Prior evidence indicates that perinatal fluoxetine treatments can influence neuronal morphology, with reports of dendritic hypotrophy within the infralimbic (IL) cortex pyramidal neurons in the mPFC (12). As we observed a differential pattern of regulation of mTOR signaling pathway components in the mPFC, with a decline noted in the PNFlx animals and an enhancement noted with the JFlx cohort, we sought to address whether this was associated with differential changes in dendritic morphology within this brain region. Using the Golgi-Cox method to label neurons, we examined the impact of a history of PNFlx (Figure 7A and Supplementary Figure 5A) and JFlx (Figure 7B and Supplementary Figure 5B) treatment on dendritic morphology of the neurons that exhibit the architecture of pyramidal neurons in layer II/III in the IL subdivision of the mPFC. We traced individual pyramidal neurons using the Neurolucida software and analysed the morphology of neurons using the Sholl and Dendogram analysis. We observed a significant decrease in the total length of the apical dendrites of layer II/III pyramidal neurons in the IL cortex in six month old animals with a history of PNFlx treatment (Figure 7C). Sholl analysis revealed a significant decrease in the number of intersections with PNFlx treatment for the apical dendrites, indicative of a decline in dendritic complexity (Figure 7D). Dendrogram analysis revealed a significant decrease in the number of higher-order apical dendrites (Figure 7E). The number of primary and secondary apical dendrites of layer II/III pyramidal neurons in the IL cortex remained unchanged in the PNFlx cohort (Figure 7E). While we observed no change in the total length of the basal dendrites (Supplementary Figure 5C), we did note a significant decline in the number of higher-order basal dendrites in layer II/III pyramidal neurons in the IL cortex of six month old animals with a history of PNFlx treatment (Supplementary Figure 5D). The number of primary and secondary basal dendrites remained unchanged following PNFlx treatment (Supplementary Figure 5D).

**Figure 7.**
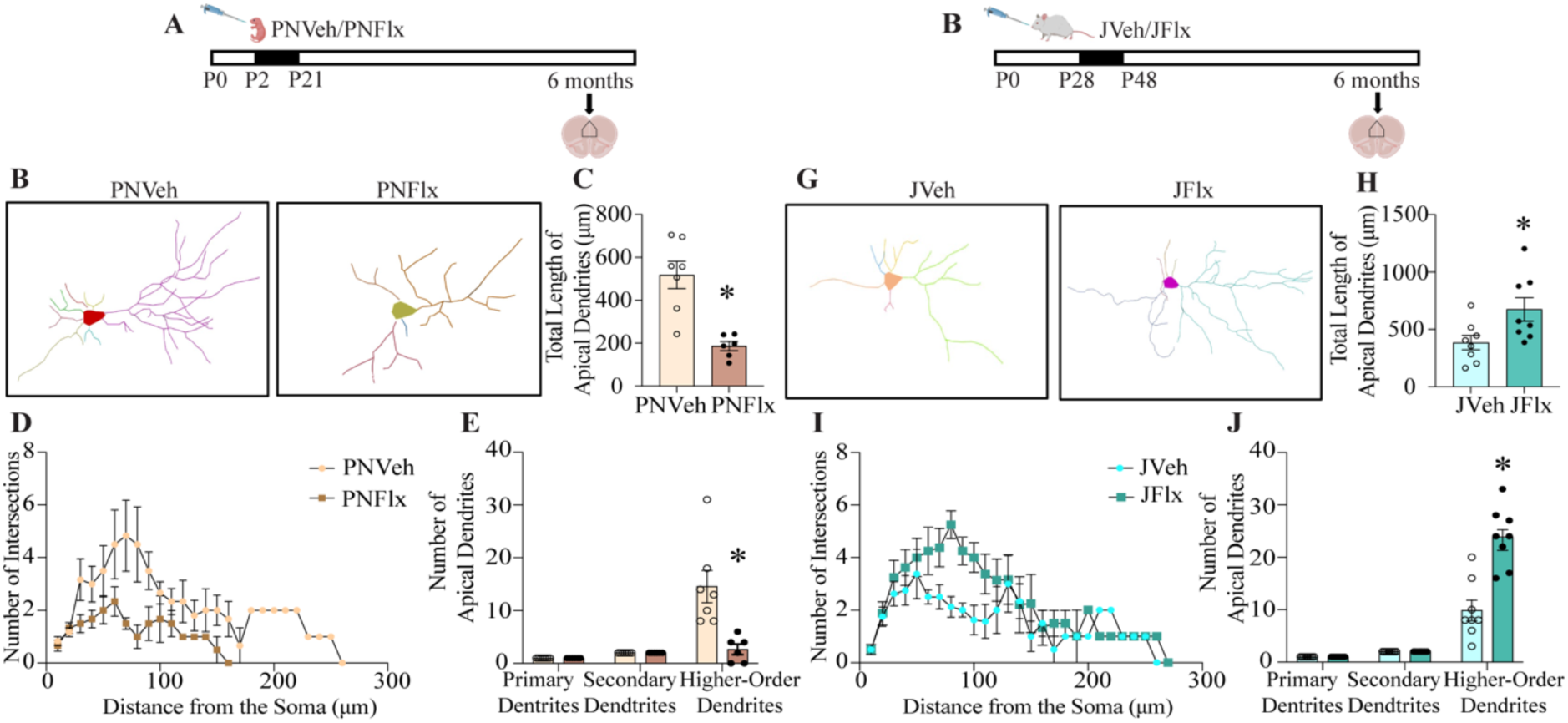
Postnatal Fluoxetine and Juvenile Fluoxetine treatment evoke opposing effects on neuronal arborization within the mPFC. Shown are schematic representations of the treatment paradigms for postnatal fluoxetine (PNFlx) (A) and juvenile fluoxetine (JFlx) (B) treatment. Shown are representative traces of Golgi–Cox-stained layer II/III pyramidal neurons in the infralimbic region (IL) of the mPFC of PNVeh (n = 7) and PNFlx-treated (n = 6) (C) animals, and JVeh and JFlx-treated (E) animals (n = 8/group) in adulthood. Shown are the quantification for the total length of apical dendrites of layer II/III pyramidal neurons in the IL cortex of PNVeh and PNFlx animals (D) and JVeh and JFlx animals (F) in adulthood. Shown are the quantification of the number of intersections per micrometer of distance from the soma across the entire apical dendritic arbor in Golgi–Cox-stained layer ⅔ pyramidal neurons in the IL subdivision of the mPFC of PNVeh and PNFlx-treated animals (G), and JVeh and JFlx animals (I). Shown is the quantification for the number of primary, secondary, and higher-order apical dendrites of layer II/III pyramidal neurons in the IL cortex of PNVeh and PNFlx-treated animals (H), and JVeh and JFlx-treated animals (J) in adulthood. Results are expressed as the number of intersections, dendrites, and total length (µm) compared with their respective age-matched, vehicle-treated controls. \**p*<0.05 compared to vehicle-treated controls (Student’s *t*-test).

In contrast, we observed a significant increase in the total length of the apical dendrites in layer II/III pyramidal neurons in the IL cortex of six month old animals with a history of JFlx treatment (Figure 7G, H). Sholl analysis indicated a significant increase in the number of intersections for apical dendrites with JFlx treatment (Figure 7I), suggesting enhanced dendritic complexity. Dendrogram analysis revealed a significant increase in the number of higher-order apical dendrites (Figure 7J) in layer II/III pyramidal neurons in the IL cortex of JFlx animals. The total length of basal dendrites remained unchanged in the JFlx cohort (Supplementary Figure 5E). The number of primary, secondary and higher-order basal dendrites (Supplementary Figure 5F) in layer II/III pyramidal neurons in the IL cortex of JFlx animals were also unchanged. These results reveal that PNFlx and JFlx treatment evokes persistent changes in the dendritic arborization of layer II/III pyramidal neurons in the IL subdivision of the mPFC.

### Postnatal fluoxetine evoked changes in despair-like, but not anxiety-like, behavior are reversed by adult-onset, nicotinamide treatment

Since, PNFlx treatment evoked enhanced anxio-depressive behaviors in adulthood, accompanied by reduced cellular ATP levels and mitochondrial function in the mPFC, we sought to address whether adult-onset treatment with compounds implicated in boosting mitochondrial function (46) could influence the behavioral consequences of PNFlx. To assess this we subjected animals with a history of PNFlx treatment to adult-onset treatment with nicotinamide (NAM, 100 mg/kg) in drinking water commencing at three months of age for a duration of thirty days (Figure 8A and Supplementary Figure 6A). NAM enhances intracellular levels of nicotinamide adenine dinucleotide (NAD^+^), and thus by boosting a critical cofactor of mitochondrial bioenergetics can serve to drive up mitochondrial biogenesis and function (47).

**Figure 8.**
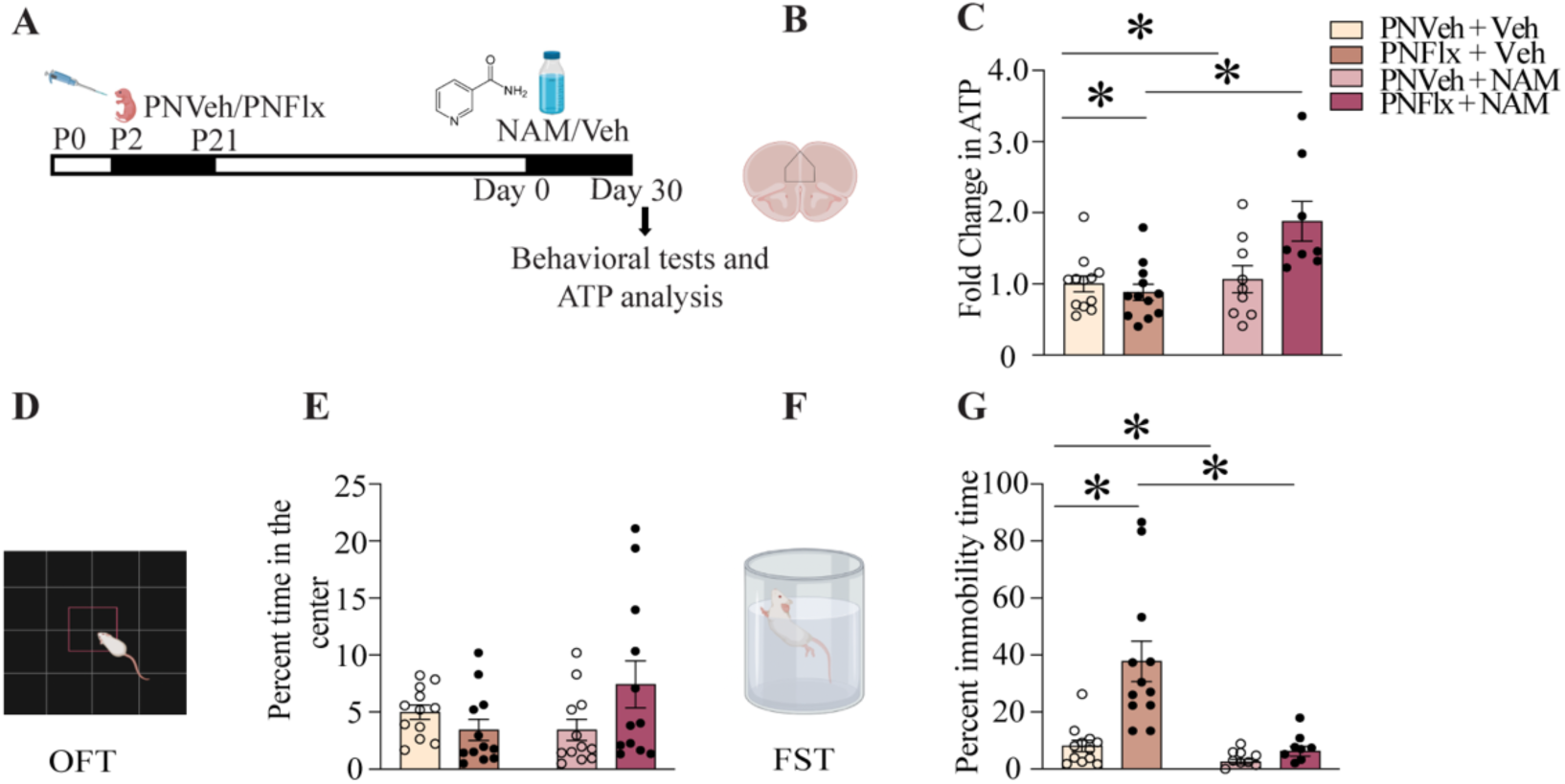
Postnatal fluoxetine evoked changes in despair-like are reversed by adult-onset nicotinamide treatment. Shown is a schematic representation of the treatment paradigm for adult-onset administration of nicotinamide (NAM) or vehicle to PNVeh and PNFlx animals, commencing at three months of age for a duration of thirty days in adulthood (A). The treatment groups were as follows: PNVeh + Vehicle, PnFlx + Vehicle, PNVeh + NAM, PNFlx + NAM. The mPFC region was processed for analysis of cellular ATP levels (B). Shown is the fold change in ATP levels (C) in the mPFC region of PNVeh and PNFlx animals with adult-onset vehicle or NAM treatment (n = 8-12). The experimental groups were assessed for anxiety-like behavior on the open field test (OFT) (D). Shown is the quantification for the percent time spent in the center of the OFT arena (E) in PNVeh and PNFlx animals before and after the NAM treatment on the OFT (n = 12/group). The treatment groups were assessed for despair-like behavior on the forced swim test (FST) (F). Shown is the quantification for the percent immobility on the FST (G) for the PNVeh and PNFlx animals with and without NAM treatment (PNVeh + Vehicle: n = 12, PNVeh + NAM: n = 9, PNFlx + Vehicle: n = 12, PNFlx + NAM: n = 8). Results are expressed as fold change, percent time in the centre of the OFT, or percent immobility on the FST and are the vehicle ± SEM. \**p*<0.05 compared to vehicle-treated controls (Two-way ANOVA analysis, Tukey *post-hoc* test).

We first assessed for the levels of cellular ATP within the mPFC, to examine whether adult-onset NAM treatment influenced the decline in cellular ATP in the mPFC observed in adult animals with a history of PNFlx treatment. Two-way ANOVA analysis indicated a significant PNFlx x NAM interaction effect for cellular ATP levels in mPFC (PNFlx x NAM; F_(1,37)_ = 8.291, *p* = 0.008) (Figure 8B, C). Multiple comparisons done using the Tukey *post-hoc* tests revealed that the significant decline in cellular ATP noted in the mPFC of PNFlx-treated animals was reversed by adult-onset treatment with NAM (Figure 8B, C). Next, we assessed the influence of adult-onset NAM treatment on anxiety-like behavior in the OFT (Figure 8D) and despair-like behavior on the FST (Figure 8F). We observed no significant interaction on the two-way ANOVA test for percent time (PNFlx x NAM: F _(1, 22)_ = 3.804, *p* = 0.064), percent distance in the centre of the OFT arena (PNFlx x NAM: F _(1, 22)_ = 2.193, *p* = 0.152), or for total distance traversed in the arena (PNFlx x NAM: F _(1, 22)_ = 3.821, *p* = 0.063) (Figure 8E and Supplementary Figure 6B-D). Two-way ANOVA analysis for despair-like behavior on the FST revealed a significant PNFlx x NAM interaction for percent immobility time (PNFlx x NAM: F_(1,37)_ = 7.84, *p* = 0.006) (Figure 8G). Tukey *post-hoc* test indicated that the significant increase in percent immobility noted in the PNFlx cohort was rescued by adult-onset NAM treatment (Figure 8G), suggesting that NAM treatment can reverse the enhanced despair-like behavior noted in adult animals with a history of PNFlx treatment.

## Discussion

The major findings of our study indicate that treatment with the SSRI, fluoxetine, in the early postnatal (P2-P21) and adolescent (P28-P48) temporal windows in male Sprague-Dawley rats, results in the programming of long-term, differential changes in anxio-depressive behaviors, accompanied by unique changes in gene expression, an opposing pattern of regulation of the mTOR signaling cascade, and starkly differing effects on mitochondrial function and dendritic architecture in the mPFC. While PNFlx treatment resulted in a sustained increase in anxiety- and despair-like behavior, JFlx treatment elicited an opposing pattern of a persistent decline in anxiety- and despair-like behavior noted at least upto six months of age. This was associated with unique, largely non-overlapping, global gene expression changes in the mPFC of PNFlx and JFlx animals. The functional analysis of the PNFlx and JFlx mPFC transcriptomes indicated the enrichment of categories such as mitochondrion and metabolic process in the downregulated genes noted following PNFlx treatment, whereas the categories of mitochondrial outer membrane and ATP binding were enriched in the upregulated genes observed with JFlx. Similarly, we noted the enrichment of functional categories such as postsynaptic membrane and dendritic spine in the downregulated genes following PNFlx, while genes upregulated following JFlx showed functional enrichment of categories such as synapse, synaptic transmission and postsynaptic density. The opposing nature of regulation was also observed with a significant decline in mitochondrial function in the mPFC of PNFlx animals in adulthood, whereas JFlx treatment evoked enhanced mitochondrial bioenergetics in the mPFC. Furthermore, PNFlx treatment resulted in a decline in expression of key components of the mTOR signaling cascade, associated with a decline in dendritic complexity of layer II/III pyramidal neurons within the IL subdivision of the mPFC. Adult animals with a history of JFlx exhibited enhanced expression of specific mTOR signaling pathway components in the mPFC, and increased dendritic arborization of layer II/III pyramidal neurons in the IL cortex. Collectively, these findings provide evidence of two distinct temporal epochs during postnatal development wherein exposure to the SSRI fluoxetine can program diametrically opposing effects on emotionality, associated with starkly differing changes in gene expression, neurometabolism and neuronal architecture within the mPFC.

Several prior reports indicate that fluoxetine administration during postnatal life, in particular within the critical period of P2-P11, evokes enhanced anxiety and despair-like behavior in adulthood (7–10,12–14,28,48). PNFlx has also been suggested to exert a differential impact on anxiety- and despair-like behavior, with an increase in stress-naïve pups and a decline in prenatally stressed pups (49–51). This underscores the importance of keeping the environmental context in consideration while interpreting the impact of postnatal fluoxetine administration on the programming of emotionality. Nevertheless, the majority of the literature including our results indicate the programming of enhanced anxio-depressive behaviors in adulthood following PNFlx treatment. In comparison, there is a lack of consensus on the impact of fluoxetine treatment during adolescence on the programming of anxio-depressive behaviors in rodent models, with differing reports indicating increased anxiety-like behavior accompanied by reduced despair-like behavior, a decline noted in both anxiety- and despair-like behavior, or no difference in either (9,21–24). These differences may arise due to a number of factors, including but not restricted to the choice of rodent species and strains, precise temporal windows targeted, dose and mode of administration of fluoxetine, variations in the treatment paradigms, as well as the behavioral tasks used to assess anxio-depressive behaviors. However, most studies have not carefully addressed the long-term persistence of the effects of postnatal and juvenile fluoxetine treatment on anxio-depressive behaviors in adulthood. Here, we provide evidence that PNFlx and JFlx treatments evoke markedly differing outcomes in their impact on anxio-depressive behaviors in adulthood, with a significant increase and decrease in emotionality respectively, that persists at least until six months of age in Sprague-Dawley male rats.

The differing outcomes on anxio-depressive behaviors evoked by PNFlx and JFlx are associated with highly distinctive, largely non-overlapping, patterns of gene regulation noted in the mPFC. This was further supported by the analysis for functional enrichment of specific gene categories in the PNFlx and JFlx mPFC transcriptomes, with enrichment of mitochondrial and synapse-associated pathways in the downregulated genes noted with PNFlx, and these similar gene categories enriched in the upregulated genes observed with JFlx. Furthermore, qPCR and western blotting analysis to examine transcript and protein levels of molecular players implicated in regulation of mitochondrial biogenesis and function (SIRT1, TFAM, ATP5A, VDAC1), revealed a decline in expression in the mPFC of PNFlx animals, and an opposing pattern of an increase in the mPFC of JFlx animals in adulthood. We also noted this pattern of a decline in expression with PNFlx and an increase following JFlx with regards to the expression of key components of the mTOR signaling pathway in the mPFC. Several studies implicate mTOR signaling in the regulation of mitochondrial biogenesis and function (41–45). We observed that adult animals with a history of PNFlx exhibited a decline in cellular ATP and mitochondrial bioenergetics in the mPFC, with an increase on these same measures noted in the mPFC of the JFlx cohorts. Collectively, these findings provide novel evidence that fluoxetine administration in the early postnatal versus juvenile window, evokes opposing changes in both the expression of key regulators of mitochondrial biogenesis and function, as well as an opposing impact on OxPhos and cellular ATP in the mPFC, a limbic brain region known to exert critical top-down control of anxio-depressive behaviors.

The mechanisms that underlie the differential impact of fluoxetine on mitochondrial function in these two temporal windows of postnatal neurodevelopment are at present unclear. Fluoxetine is known to enhance serotonin levels by blocking the serotonin transporter (52), and serotonin has recently been shown to be a direct regulator of mitochondrial biogenesis and function via a 5-HT_2A_ receptor dependent recruitment of Sirt1 signaling, albeit in adulthood (34). Interestingly, previous studies indicate the direct (53) and indirect effects (54,55) of the administration of fluoxetine in adulthood on the regulation of mitochondrial function. These effects of adult fluoxetine administration on the regulation of mitochondrial function vary depending on the duration of treatment and the region of interest (37). The influence of serotonin on regulation of mitochondrial biogenesis and function during developmental epochs has been relatively unexplored, with a few reports suggesting that PNFlx enhances mitochondrial bioenergetics in the periphery, namely the liver and heart (56,57) and that JFlx can reverse the hypothalamic mitochondrial dysfunction evoked by early nutritional imbalance (56). Interestingly, reports also indicate enrichment of fluoxetine in mitochondrial fractions in the brain (36), and suggest that fluoxetine may directly interact with VDAC1 to inhibit the mitochondrial permeability transition pore (58). However, it is still unclear why the consequences of serotonin elevation on mitochondria in the mPFC in these postnatal temporal windows are diametrically opposing. One possibility is that the receptor milieu through which serotonin exerts its effects are dynamically shifting during these temporal epochs (59–61), which may impact the nature of signaling pathways recruited in the mPFC. For example, the mTORC1 complex is regulated by diverse 5-HT receptors in distinct brain regions, with 5-HT_1A_ and 5-HT_7_ driving the Ras-PI3K-Akt-mTOR pathway in hippocampal neurons leading to positive regulation of dendritogenesis (62). A history of PNFlx is reported to reduce mTOR signaling in the hippocampus in adulthood (10), and here we find a decline in phosphorylation of several components of the mTOR cascade in the mPFC of PNFlx animals. Our findings motivate detailed future studies to uncover the mechanisms via which fluoxetine exerts differing effects on the expression of mitochondrial biogenesis and function in the mPFC in postnatal developmental epochs.

Several studies suggest a vital role for mitochondria that extend beyond their well-appreciated role in bioenergetics (63), with functions in cellular signaling (64–66), stress adaptation (67,68), synaptic transmission (69,70), growth (71), maintenance and plasticity of dendritic cytoarchitecture (72,73), and more recent, tantalizing reports of a role in the regulation of emotional states (46,74–76) and cognitive behavior (77,78). In this regard, it is of interest that we observed a decline in dendritic complexity of layer II/III pyramidal neurons in the IL subdivision of the mPFC that accompanied the mitochondrial hypofunction noted in the mPFC in PNFlx animals in adulthood. Our results are in agreement with a prior report that indicated dendritic hypotrophy and reduced excitability of layer II/III IL pyramidal neurons following administration of fluoxetine from P2-P11 (12), which was hypothesized to contribute to the anxiogenic effects of PNFlx treatment. Indeed, IL lesions can phenocopy specific aspects of the behavioral phenotypes associated with PNFlx, namely the enhanced anxiety and deficits in fear extinction (12). It is tempting to speculate that our observations of mitochondrial hypofunction within the mPFC with PNFlx treatment, may contribute to the reduced dendritic complexity and the decline in excitability reported to arise with PNFlx treatment (12). In contrast, we find that JFlx animals exhibit enhanced dendritic complexity of layer II/III pyramidal neurons in the IL cortex, as well as increases in cellular ATP and OxPhos, and the expression of several proteins implicated in the regulation of mitochondrial biogenesis and function. Our results along with the current literature hints towards a complex interplay between mitochondrial dysfunction, decreased dendritic complexity and neuronal hypoactivity of IL pyramidal neurons (12) which could contribute to the programming of altered emotionality in PNFlx animals. It would be of interest to delineate whether the mitochondrial dysfunction is the primary driver in contributing to the hypofunctioning of mPFC or whether the reduced excitability noted in the mPFC IL pyramidal neurons leads to lower energy demands thus impacting mitochondrial function in PNFlx animals. While at present our results do not extend beyond the correlative, they indicate that PNFlx and JFlx produce starkly opposing effects on mitochondrial function and neuronal cytoarchitecture in the mPFC, which accompany highly divergent outcomes on anxio-depressive behaviors, raising the intriguing possibility that the changes in mitochondrial function may contribute to the shaping of long-term behavioral consequences.

To explore the relationship between mitochondrial hypofunction in the mPFC evoked by PNFlx and the enhanced anxio-depressive behaviors that are observed with PNFlx, we assessed whether adult-onset treatment with NAM, a precursor of NAD^+^ that enhances mitochondrial biogenesis and function can restore the enhanced anxio-depressive behavioral phenotype evoked by PNFlx. We observed that treatment with NAM in adulthood restores the decline in cellular ATP in the mPFC evoked by PNFlx, and also rescues the enhanced despair-like, but not anxiety-like, behavior noted with PNFlx. These results point to the possibility of a role for mitochondrial hypofunction in the mPFC in programming the depressive behavioral outcomes of PNFlx treatment. In conclusion, our results indicate that SSRI treatment in postnatal versus juvenile windows evokes opposing effects on anxiety- and despair-like behavior, which are accompanied by starkly differing regulation of gene expression, mitochondrial bioenergetics, mTOR signaling and dendritic complexity within the mPFC. These findings motivate future experiments to explore the relationship between altered mitochondrial function in the mPFC and the changes in dendritic cytoarchitecture, neuronal function and emotionality evoked by elevated serotonergic neurotransmission in key neurodevelopmental epochs.

## CRediT authorship contribution statement

**Utkarsha Ghai**- Visualization, investigation, writing original draft. **Parul Chachra**- Visualization, investigation, **Sashaina E. Fanibunda**- Investigation, **Suchith Mendon**- Investigation, **Amogh Bhaskaran**- Investigation, **Ambalika Sarkar**- Investigation, **Kowshik Kukkemane**- Investigation, **Vivek Singh**- Investigation, **Vidita A. Vaidya**- Conceptualization, writing and editing original draft, funding acquisition.

## Author Contribution

UG, PC, SF, SM, AB, AS, KK, VV contributed to experiments and data analysis. UG and VV wrote the manuscript.

## Acknowlegdements

This work was supported by intramural funding from the Tata Institute of Fundamental Research and the Department of Atomic Energy, Mumbai (RTI4003 to VV). We thank Dr. Shital Suryavanshi, K.V. Boby, Jaya Moorjani, and the animal house staff at the Tata Institute of Fundamental Research for technical assistance.

