## Supplementary Information for "Fluoxetine treatment during postnatal and juvenile temporal epochs evokes diametrically opposing changes in anxio-depressive behaviors, gene expression, mitochondrial function, and neuronal architecture in the medial prefrontal cortex"

### Supplementary Figure and Figure Legends

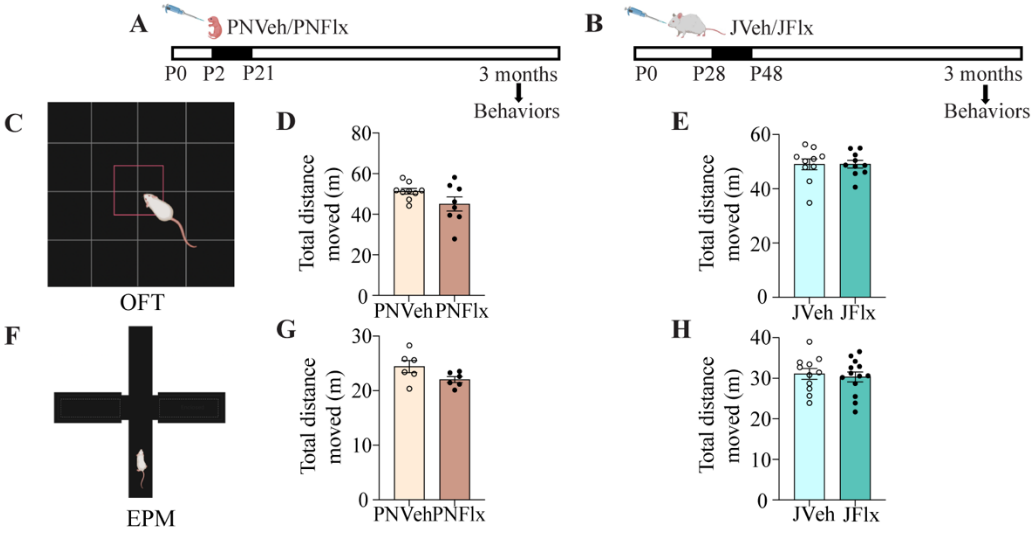


*Supplementary Figure 1. Postnatal and juvenile fluoxetine treatment does not alter the total distance moved on the OFT and EPM at three months of age.*

Shown is a schematic representation of the treatment paradigm for postnatal fluoxetine (PNFlx) (A) from postnatal day 2 (P2) to P21 and juvenile fluoxetine (JFlx) (B) treatment from P28 to P48, with assessment for anxiety-like behavior at three months of age on the Open Field Test (OFT) (C) and Elevated Plus Maze (EPM) (F). Shown is the total distance traversed (D: PNFLx experiment; E: JFlx Experiment) in the OFT arena (PNVeh: n = 9, PNFlx: n = 8; JVeh: n =10 and JFlx: n = 10). Shown is the total distance traversed (G: PNFlx experiment; H: JFlx Experiment) in the EPM arena (PNVeh: n = 6, PNFlx: n = 6; JVeh: n = 11 and JFlx: n = 13). Results are expressed as the mean ± SEM. *$p<0.05$ as compared to vehicle (Student's *t*-test).

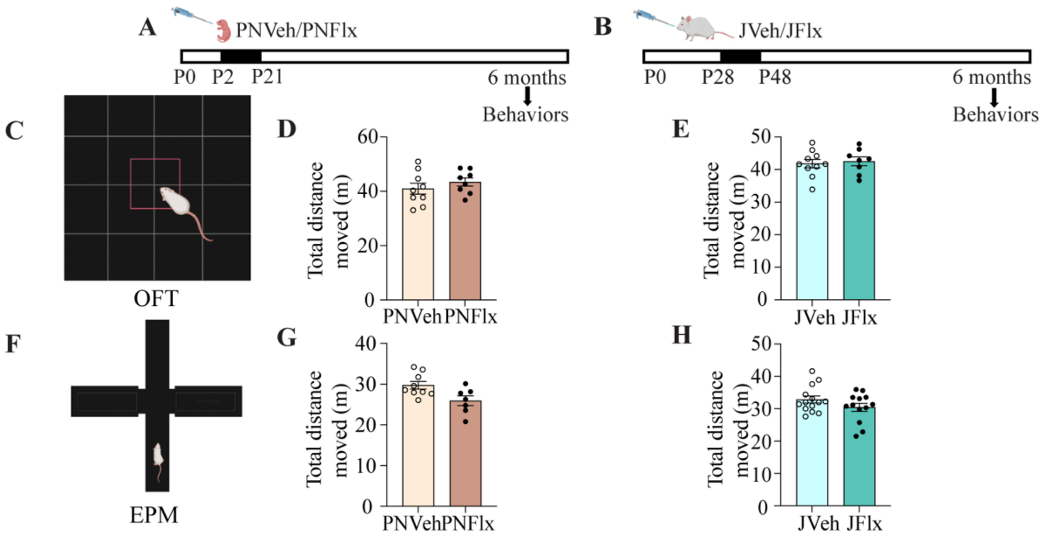


*Supplementary Figure 2. Postnatal and juvenile fluoxetine treatment does not alter the total distance moved on the OFT and EPM at six months of age.*

Shown is a schematic representation of the treatment paradigm for postnatal fluoxetine (PNFlx) (A) from postnatal day 2 (P2) to P21 and juvenile fluoxetine (JFlx) (B) treatment from P28 to P48, with assessment for anxiety-like behavior at six months of age on the Open Field Test (OFT) (C) and Elevated Plus Maze (EPM) (F). Shown is the total distance traversed (D: PNFlx experiment; E: JFlx Experiment) in the OFT arena (PNVeh: n = 9, PNFlx: n = 8; JVeh: n = 10 and JFlx: n = 8). Shown is the total distance traversed (G: PNFlx experiment; H: JFlx Experiment) in the EPM arena (PNVeh: n = 9, PNFlx: n = 7; JVeh: n = 14, JFlx: n = 13). Results are expressed as the mean ± SEM. *$p$<0.05 as compared to vehicle (Student's *t*-test).

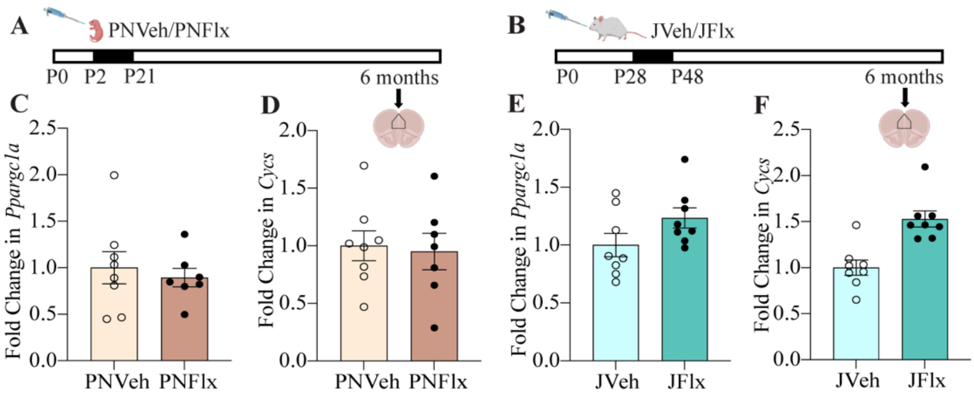


*Supplementary Figure 3. Postnatal and juvenile fluoxetine does not alter gene expression of Ppargc1a and Cycs at six months of age.*

Shown is a schematic representation of the treatment paradigm for postnatal fluoxetine (PNFlx) (A) and juvenile fluoxetine (JFlx) (B) treatment. Shown is the quantification of fold change in gene expression of *Ppargc1a* (C) and *Cycs* (D) in the mPFC following PNFlx treatment (PNVeh: n = 8, PNFlx: n = 7). Shown is the quantification of fold change in gene expression of *Ppargc1a* (E) and *Cycs* (F) in the mPFC following JFlx treatment (n = 8/group). Results are expressed as the fold change ± SEM. *$p<0.05$ compared to vehicle-treated (Student's *t*-test).

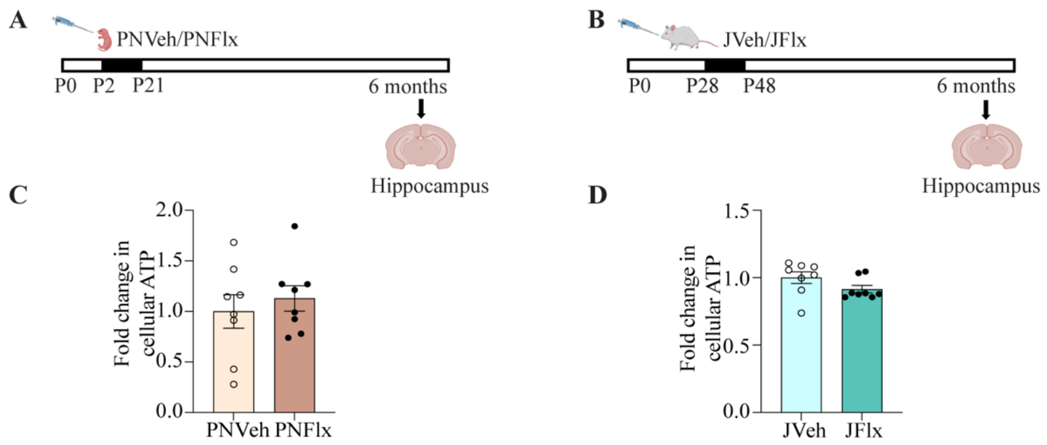


*Supplementary Figure 4. Postnatal and juvenile fluoxetine treatments evoke no change in cellular ATP levels in the hippocampus.*

Shown is a schematic representation of the treatment paradigms for postnatal (PNFlx) (A) and juvenile fluoxetine (JFlx) (B) treatment. Shown is the quantification of cellular ATP levels in the mPFC of PNVeh and PNFlx-treated animals (C) in adulthood (n = 8/group). Shown is the quantification of cellular ATP levels (D) in the mPFC of JVeh and JFlx-treated animals in adulthood (n = 8/group). Results are expressed as fold change ± SEM compared to age-matched vehicle controls. *$p$<0.05 compared to vehicle (Student's *t*-test).

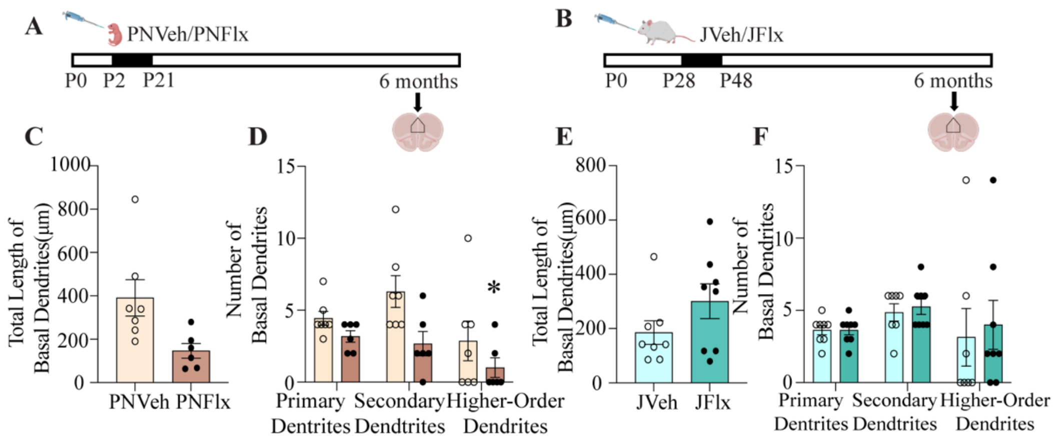


*Supplementary Figure 5. Postnatal Fluoxetine and Juvenile Fluoxetine treatment evokes no change in the total length of basal dendrites in layer II/III pyramidal neurons in the infralimbic subdivision of the mPFC.*

Shown are schematic representations of the treatment paradigms for postnatal fluoxetine (PNFlx) (A) and juvenile fluoxetine (JFlx) (B) treatment. Shown are the quantification for the total length of basal dendrites of layer II/III pyramidal neurons in the IL cortex of PNVeh (n =7) and PNFlx-treated (n = 6) (C) animals. Shown is the quantification for the number of primary, secondary, and higher-order basal dendrites of layer II/III pyramidal neurons in the IL cortex of PNVeh and PNFlx-treated animals (D). Shown are the quantification for the total length of basal dendrites of layer II/III pyramidal neurons in the IL cortex of JVeh and JFlx-treated (E) animals (n = 8/group) in adulthood. Shown is the quantification for the number of primary, secondary, and higher-order basal dendrites of layer II/III pyramidal neurons in the IL cortex of JVeh and JFlx-treated animals (F) in adulthood. Results are expressed as the number of dendrites, and total length (µm) compared with their respective age-matched, vehicle-treated controls. *$p$<0.05 compared to vehicle-treated controls (Student’s $t$-test).

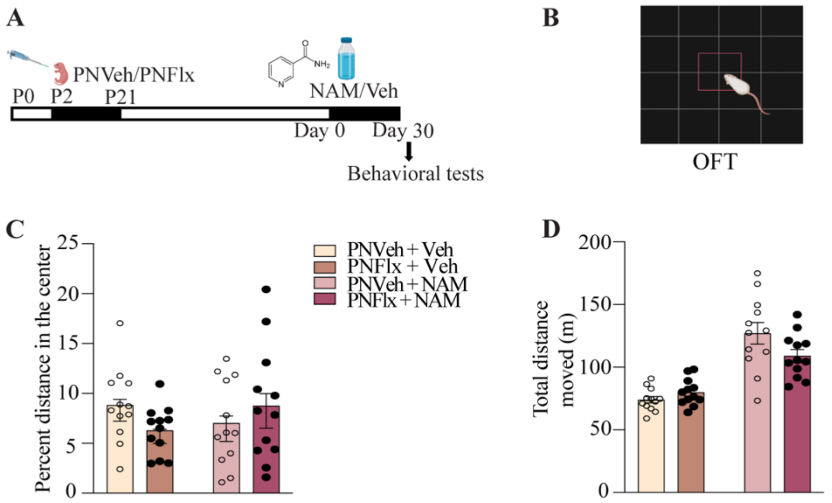


*Supplementary Figure 6. Influence of adult-onset NAM treatment of animals with a history of PNFlx on behavior in the OFT.*

Shown is a schematic representation of the treatment paradigm for adult-onset administration of nicotinamide (NAM) or vehicle to PNVeh and PNFlx animals, commencing at three months of age for a duration of thirty days in adulthood (A). The treatment groups were as follows: PNVeh + Vehicle, PNFlx + Vehicle, PNVeh + NAM, PNFlx + NAM. The experimental groups were assessed for anxiety-like behavior on the open field test (OFT) (B). Shown is the quantification for the percent distance in the center (C) and the total distance traversed in the OFT arena (D) by the PNVeh and PNFlx animals before and after the NAM treatment on the OFT (n = 12/group). Results are expressed as percent distance in the centre of the OFT and the total distance traversed, and are the vehicle ± SEM (Two-way ANOVA analysis, repeated measures).

**Supplementary Table 1: Primer sequences used for validation of microarray target genes.**

| Gene name | Gene symbol | Primer sequence (5'-3') |
|---|---|---|
| Adhesion G Protein-Coupled Receptor L2 | *Lphn2* | F - TGTCCTGGAACCTTGAAAGC<br>R - AATCCAGTACCGTCCACTCG |
| Similar to ETS domain-containing protein Elk-1 | *Elk1* | F - ACCACTGAGATCACCCAACC<br>R -AGGTATGTGTGGGGAGCAAG |
| Glycogen Synthase Kinase 3 Alpha | *Gsk3a* | F - AGGTGGCTTACACCGACATC<br>R – AGCCTCACGATATTGCAGTG |
| Guanine nucleotide binding protein α inhibiting 2 | *Gnai2* | F - ACGGACACATCCATCATCCT<br>R - TCCGTTACAGCATCGAACAC |
| Hypoxanthine-guanine-phosphoribosyl transferase 1 | *Hprt1* | F - GCAGACTTTGCTTTCCTTGG<br>R - GTCTGGCCTGTATCCAACACT |
| SH3/ankyrin domain gene 2 isoform a | *Shank2* | F - AGACATGAGCGCACAGAATG<br>R - AGGGCACAATGTCTGTTTCC |
| Uncoupling protein 1 | *UCP1* | F - AGAAACGCCTGCCTCTTTG<br>R - AAGCATTGTAGGTCCCAGTGT |
| Arrestin, beta 2 | *Arrb2* | F - GAGATTCGAGCCTTCTGTGC<br>R - CCGGTCAGACATGAGGAAGT |
| Glutamate receptor, metabotropic 5 | *Grm5* | F - TGAGAGGAAGTGTGGTGCAG<br>R - CTTTGTTCTAGGGCCACAGC |
| S100 calcium binding protein alpha | *S100α* | F - AGCTGAGCAAGAAGGAGCTG<br>R - TTGTTACAAGCCACCGTGAG |
| N-ethylmaleimide sensitive fusion protein | *Nsf* | F - TGGGCTGGACTTTCTATTGG<br>R - TGAACTGCTGGATGAACTCG |

**Supplementary Table 2: Primer sequences used for quantitative PCR analysis.**

| Gene name | Gene symbol | Primer sequence (5'-3') |
|---|---|---|
| *Ppargc1a* | peroxisome proliferator-activated receptor gamma, coactivator 1 alpha | F - TGAACTACGGGATGGCAACT<br>R - GAAGAGCAAGAAGGCGACAC |
| *Tfam* | transcription factor A, mitochondrial | F - GCTAAACACCCAGATGCAAA<br>R - GCTTCCTTCTCTAAGCCCATC |
| *Sirt1* | sirtuin 1 | F -AGAACCACCAAAGCGGAAA<br>R-ACAGCAAGGCGAGCATAAA |
| *Cycs* | cytochrome c, somatic | F-ACCAGCCCGGACCGAATTTA<br>R-GTGTAAGAGAATCCAGCAGCCT |

**Supplementary Table 3: Primer sequences used for analysis of relative mitochondrial DNA (mtDNA) content.**

| Gene name | Gene symbol | Primer sequence (5'-3') |
|---|---|---|
| mt-Cytb | cytochrome b, mitochondrial | F-ACGCTCCATTCCCAACAAAC<br><br>R-GTTGGCCTCCGATTCATGTT |
| Cycs | cytochrome c, somatic | F-<br>AGGCTGCTGGATTCTCTTACA<br><br>R-GTCTGCCCTTTCTCCCTTCT |

**Supplementary Table 4: List of genes differentially upregulated in the mPFC transcriptome of adult PNFlx animals**

Six month old PNFlx animals and their age-matched controls were subjected to microarray analysis. The list of all the significantly upregulated genes is shown in the table ($p < 0.05$ ; Fold change > 1.35).

| Systematic Name | GeneName | *P* Value | Fold change | Product |
|---|---|---|---|---|
| AA875107 | AA875107 | 0.013 | 6.07 | CI-B9 mRNA for ubiquinone oxidoreductase complex |
| XM_341669 | Abce1 | 0.047 | 1.76 | similar to ATP-binding cassette sub-family E member 1 |
| XM_231137 | Abl1_mapped | 0.034 | 2.57 | similar to Proto-oncogene tyrosine-protein kinase ABL1 |
| XM_574921 | Acad9 | 0.001 | 16.40 | similar to very-long-chain acyl-CoA dehydrogenase VLCAD homolog isoform 2 |
| NM_019144 | Acp5 | 0.013 | 4.93 | acid phosphatase 5 |
| NM_022600 | Adcy5 | 0.003 | 1.91 | adenylate cyclase 5 |
| NM_022681 | Adnp | 0.005 | 2.61 | activity-dependent neuroprotective protein |
| NM_017161 | Adora2b | 0.032 | 1.57 | adenosine A2B receptor |
| NM_022618 | Akap6 | 0.049 | 1.73 | A-kinase anchor protein 6 |
| XM_001059377 | Ankrd32_predicted | 0.034 | 2.10 | similar to ankyrin repeat domain 32 |
| NM_001011918 | Anxa11 | 0.046 | 3.24 | annexin A11 |
| XM_230761 | Apba2bp_predicted | 0.000 | 7.06 | similar to amyloid beta precursor protein-binding, family A, member 1 binding protein |
| NM_181083 | Arfgef2 | 0.004 | 1.78 | ADP-ribosylation factor guanine nucleotide-exchange factor 2 |
| XM_224637 | Arhgap22_predicted | 0.002 | 5.96 | similar to Rho GTPase activating protein 22 |
| NM_001013246 | Arhgef12 | 0.003 | 1.91 | Rho guanine nucleotide exchange factor (GEF) 12 |
| NM_019186 | Arl4a | 0.002 | 6.48 | ADP-ribosylation factor-like 4 |
| NM_053311 | Atp2b1 | 0.040 | 14.44 | plasma membrane calcium ATPase 1 |
| NM_134364 | Atp5b | 0.016 | 2.68 | ATP synthase, H+ transporting, mitochondrial F1 complex, beta subunit |
| NM_053756 | Atp5g3 | 0.034 | 4.91 | ATP synthase, H+ transporting, mitochondrial F0 complex, subunit c, isoform 3 |
| NM_001013158 | B3galt3 | 0.011 | 2.72 | UDP-Gal:betaGlcNAc beta 1,3-galactosyltransferase, polypeptide 3 |
| NM_022698 | Bad | 0.005 | 4.89 | bcl2-associated death promoter |
| XM_215664 | Bcas2_predicted | 0.016 | 2.42 | similar to Breast carcinoma amplified sequence 2 homolog |
| BM391736 | BM391736 | 0.002 | 3.03 | UI-R-DZ0-cks-e-06-0-UI.s1 NCI_CGAP_DZ |
| NM_001009604 | Bri3 | 0.000 | 79.28 | brain protein I3 |
| XM_236307 | Brunol6_predicted | 0.008 | 1.63 | similar to bruno-like 6, RNA binding protein |
| NM_012517 | Cacna1c | 0.050 | 1.53 | calcium channel, voltage-dependent, L type, alpha 1C subunit |
| NM_012727 | Camk4 | 0.022 | 1.66 | calcium/calmodulin-dependent protein kinase IV |

| | | | | |
|---|---|---|---|---|
| XM_341784 | Cdc42ep5_predicted | 0.009 | 2.71 | similar to CDC42 effector protein 5 |
| XM_341076 | Cdk2ap1_predicted | 0.032 | 12.65 | similar to CDK2-associated protein 1 |
| NM_031762 | Cdkn1b | 0.034 | 1.84 | cyclin-dependent kinase inhibitor 1B |
| NM_173103 | Clcnkb | 0.025 | 3.54 | chloride channel Kb |
| NM_019299 | Cltc | 0.039 | 1.57 | clathrin, heavy polypeptide |
| NM_012784 | Cnr1 | 0.044 | 2.09 | cannabinoid receptor 1 |
| XM_214400 | Col4a1 | 0.032 | 1.80 | similar to Collagen alpha-1(IV) chain precursor |
| NM_053472 | Cox4i2 | 0.033 | 1.52 | cytochrome c oxidase subunit IV isoform 2 precursor |
| NM_012836 | Cpd | 0.032 | 1.37 | carboxypeptidase D |
| NM_133381 | Crebp | 0.046 | 1.46 | CREB binding protein |
| XM_220093 | Crsp3 | 0.026 | 1.81 | similar to cofactor required for Sp1 transcriptional activation, subunit 3 |
| XM_217086 | Crsp6 | 0.032 | 1.39 | similar to cofactor required for Sp1 transcriptional activation, subunit 6 |
| NM_022177 | Cxcl12 | 0.023 | 2.50 | chemokine ligand 12 isoform alpha |
| NM_080891 | Daxx | 0.047 | 1.50 | Fas death domain-associated protein |
| NM_053379 | Dcx | 0.008 | 2.88 | doublecortin |
| XM_342565 | Dhx35_predicted | 0.001 | 4.02 | similar to Probable ATP-dependent RNA helicase DHX35 |
| XM_215751 | Dnajc10 | 0.036 | 2.00 | similar to ER-resident protein ERdj5 |
| NM_001012205 | Dpp10 | 0.001 | 7.82 | dipeptidylpeptidase 10 |
| XM_216479 | Echdc2_predicted | 0.013 | 1.68 | similar to enoyl Coenzyme A hydratase domain containing 2 |
| XM_226624 | Ell2 | 0.021 | 2.48 | similar to RNA polymerase II elongation factor ELL2 |
| NM_001004228 | Emcn | 0.027 | 5.18 | endomucin |
| NM_134331 | Epha7 | 0.049 | 1.84 | EphA7 |
| NM_198742 | Etfdh | 0.000 | 3.50 | electron-transferring-flavoprotein dehydrogenase |
| NM_012555 | Ets1 | 0.023 | 2.01 | v-ets erythroblastosis virus E26 oncogene homolog 1 |
| NM_012555 | Ets1 | 0.044 | 1.76 | v-ets erythroblastosis virus E26 oncogene homolog 1 |
| NM_024147 | Evl | 0.025 | 1.76 | Ena-vasodilator stimulated phosphoprotein |
| NM_138850 | Fap | 0.025 | 4.08 | fibroblast activation protein |
| XM_213746 | Fis1 | 0.003 | 3.17 | similar to tetratricopeptide repeat domain 11 |
| NM_001012137 | Fus | 0.008 | 1.64 | fusion (involved in t(12;16) in malignant liposarcoma) |
| XM_216605 | G1p2_predicted | 0.010 | 3.19 | similar to Ubiquitin cross-reactive protein precursor |
| NM_012793 | Gamt | 0.003 | 2.04 | guanidinoacetate methyltransferase |
| NM_024375 | Gdf10 | 0.028 | 1.36 | growth differentiation factor 10 |
| NM_017276 | Gdi2 | 0.032 | 1.81 | GDP dissociation inhibitor 2 |
| XM_343817 | Gla_mapped | 0.046 | 2.58 | similar to Alpha-galactosidase A precursor |
| NM_053765 | Gne | 0.004 | 2.13 | UDP-N-acetylglucosamine-2-epimerase/N-acetylmannosamine kinase |

| | | | | |
|---|---|---|---|---|
| NM_030846 | Grb2 | 0.006 | 2.03 | growth factor receptor bound protein 2 |
| NM_012796 | Gstt2 | 0.048 | 1.62 | glutathione S-transferase, theta 2 |
| XM_230734 | H13_predicted | 0.006 | 4.02 | similar to minor histocompatibility antigen 13 isoform 3 |
| NM_001008357 | Hcfc2 | 0.030 | 1.61 | host cell factor C2 |
| AF321132 | Hdac4 | 0.008 | 1.60 | histone deacetylase 4 |
| NM_145785 | Hdgfrp3 | 0.028 | 2.41 | hepatoma-derived growth factor, related protein 3 |
| NM_198132 | Hnrpa3 | 0.000 | 14.29 | heterogeneous nuclear ribonucleoprotein A3 |
| XM_342131 | Hnrph3_predicted | 0.019 | 7.70 | similar to heterogeneous nuclear ribonucleoprotein H3 isoform a |
| XM_214927 | Htatip2_predicted | 0.014 | 1.69 | similar to HIV-1 tat interactive protein 2, homolog |
| NM_053783 | Ifngr | 0.035 | 3.21 | interferon gamma receptor 1 |
| NM_017020 | Il6ra | 0.021 | 1.66 | interleukin 6 receptor, alpha |
| XM_342650 | Impdh1_predicted | 0.010 | 1.76 | similar to inosine monophosphate dehydrogenase 1 isoform b |
| NM_001012131 | Inpp1 | 0.031 | 2.13 | inositol polyphosphate-1-phosphatase |
| NM_031002 | Inpp4a | 0.030 | 1.82 | inositol polyphosphate-4-phosphatase, type 1 |
| NM_053949 | Kcnh2 | 0.011 | 8.27 | voltage-gated potassium channel, subfamily H, member 2 |
| NM_013192 | Kcnj6 | 0.000 | 1.44 | potassium inwardly-rectifying channel, subfamily J, member 6 |
| NM_198726 | Kpna1 | 0.037 | 2.57 | karyopherin alpha 1 (importin alpha 5) |
| NM_198726 | Kpna1 | 0.045 | 1.75 | karyopherin alpha 1 (importin alpha 5) |
| L04739 | L04739 | 0.019 | 1.54 | plasma membrane calcium ATPase |
| NM_001013188 | Leprotl1 | 0.022 | 5.58 | leptin receptor overlapping transcript-like 1 |
| NM_022226 | Lgmn | 0.004 | 3.28 | legumain |
| NM_139112 | Lmo3 | 0.035 | 2.19 | LIM domain only 3 |
| NM_139101 | LOC245960 | 0.049 | 1.40 | potassium channel regulator 1 |
| XM_233435 | LOC313535 | 0.014 | 5.81 | similar to prion protein interacting protein 1 |
| NM_001014269 | LOC367314 | 0.015 | 1.40 | leucine rich repeat (in FLII) interacting protein 1 |
| NM_001035255 | LOC502603 | 0.000 | 24.61 | splicing factor, arginine/serine-rich 11 |
| XM_001053080 | LOC678709 | 0.039 | 4.90 | similar to Glucocorticoid receptor DNA-binding factor 1 |
| XM_001054517 | LOC679060 | 0.035 | 1.36 | similar to transmembrane emp24 protein transport domain containing 7 |
| NM_130822 | Lphn3 | 0.007 | 3.54 | calcium-independent alpha-latrotoxin receptor homolog 3 |
| XM_228137 | Lrrtm3_predicted | 0.018 | 2.53 | similar to leucine-rich repeat transmembrane neuronal 3 protein |
| NM_031655 | Lxn | 0.000 | 78.93 | latexin |
| XM_215213 | Mcee_predicted | 0.017 | 7.33 | similar to Methylmalonyl-CoA epimerase, mitochondrial precursor |
| NM_131904 | Mgea5 | 0.043 | 2.10 | meningioma expressed antigen 5 |
| NM_138843 | Mpst | 0.001 | 3.50 | 3-mercaptopyruvate sulfurtransferase |

| | | | | |
|---|---|---|---|---|
| NM_001006973 | mrpl11 | 0.005 | 2.70 | mitochondrial ribosomal protein L11 |
| XM_213513 | Mrps7 | 0.014 | 2.76 | similar to mitchondrial ribosomal protein S7 |
| NM_212536 | Msh5 | 0.019 | 5.13 | mutS homolog 5 |
| XM_227485 | Msr2_predicted | 0.010 | 29.36 | similar to macrophage scavenger receptor 2 |
| XM_341374 | Ndfip2_predicted | 0.024 | 1.58 | similar to Nedd4 family interacting protein 2 |
| NM_031967 | Ndrg4 | 0.000 | 6.20 | N-myc downstream regulated 4 |
| NM_053516 | Nol3 | 0.002 | 11.47 | nucleolar protein 3 |
| NM_001012356 | Nono | 0.031 | 1.42 | non-POU domain containing, octamer-binding |
| NM_053750 | Nppc | 0.032 | 1.73 | natriuretic peptide precursor type C |
| NM_001033951 | Npuk68 | 0.030 | 1.54 | nuclear protein UKp68 isoform 1 |
| NM_080766 | Nras | 0.033 | 1.51 | neuroblastoma RAS viral (v-ras) oncogene homolog |
| NM_001013925 | Nub1 | 0.002 | 5.60 | Nedd8 ultimate buster-1 |
| NM_001006991 | Nudt9 | 0.013 | 1.76 | nudix -type motif 9 |
| XM_001056084 | Nup153 | 0.049 | 1.64 | similar to Nuclear pore complex protein Nup153 |
| XM_230286 | Nup160_predicted | 0.016 | 1.40 | similar to nucleoporin 160 |
| NM_001000476 | Olr1330_predicted | 0.049 | 2.13 | olfactory receptor Olr1330 |
| NM_001000016 | Olr1439_predicted | 0.006 | 4.07 | olfactory receptor Olr1439 |
| NM_001000105 | Olr1658_predicted | 0.005 | 1.84 | olfactory receptor Olr1658 |
| NM_201422 | Pcdhac2 | 0.039 | 1.80 | protocadherin alpha subfamily C, 2 |
| NM_138543 | Pde9a | 0.041 | 1.45 | phosphodiesterase 9A |
| NM_133284 | Pgc | 0.003 | 2.67 | progastricsin |
| XM_216565 | Pink1_predicted | 0.004 | 1.75 | similar to PTEN induced putative kinase 1 |
| NM_053772 | Pkia | 0.015 | 1.55 | protein kinase inhibitor, alpha |
| NM_199101 | Plekha4 | 0.019 | 1.38 | pleckstrin homology domain containing, family A member 4 |
| XM_342710 | Pole4_predicted | 0.006 | 2.35 | similar to DNA polymerase epsilon subunit 4 |
| NM_031335 | Polr2f | 0.001 | 18.44 | polymerase II |
| NM_031347 | Ppargc1a | 0.039 | 5.71 | peroxisome proliferative activated receptor, gamma, coactivator 1 alpha |
| XM_579602 | Ppat | 0.005 | 1.76 | similar to Amidophosphoribosyltransferase precursor |
| NM_053890 | Ppp1r12a | 0.000 | 17.56 | protein phosphatase 1, regulatory (inhibitor) subunit 12A |
| NM_022209 | Ppp2r2b | 0.000 | 160.19 | BRbeta B-regulatory subunit of protein phosphatase 2A |
| NM_022209 | Ppp2r2b | 0.037 | 1.59 | BRbeta B-regulatory subunit of protein phosphatase 2A |
| XM_001062510 | Ppp2r5d | 0.009 | 2.17 | similar to delta isoform of regulatory subunit B56, protein phosphatase 2A isoform 1 |
| XM_216225 | Ppp4r2_predicted | 0.016 | 1.51 | similar to protein phosphatase 4, regulatory subunit 2 |

| | | | | |
|---|---|---|---|---|
| BC079072 | Pqlc1 | 0.007 | 4.89 | PQ loop repeat containing 1 |
| BC093606 | Psip1 | 0.037 | 1.72 | Psip1 protein |
| NM_175765 | Psip1 | 0.026 | 1.49 | PC4 and SFRS1 interacting protein 1 |
| NM_001031639 | Psmd2 | 0.000 | 5.38 | proteasome 26S subunit, non-ATPase, 2 |
| NM_031606 | Pten | 0.002 | 2.87 | phosphatase and tensin homolog |
| NM_031975 | Ptms | 0.037 | 2.05 | parathymosin |
| NM_017066 | Ptn | 0.001 | 37.09 | pleiotrophin |
| XM_214440 | Ptpdc1_predicted | 0.000 | 3.73 | similar to protein tyrosine phosphatase domain containing 1 protein |
| NM_031090 | Rab1 | 0.004 | 1.81 | RAB1, member RAS oncogene family |
| NM_031152 | Rab11a | 0.000 | 6.63 | RAB11a, member RAS oncogene family |
| NM_023950 | Rab7 | 0.021 | 1.98 | RAB7, member RAS oncogene family |
| XM_216367 | Rarsl_predicted | 0.001 | 10.48 | similar to arginyl-tRNA synthetase-like |
| XM_221731 | Rbm11_predicted | 0.031 | 1.64 | similar to RNA binding motif protein 11 |
| NM_001007618 | Rchy1 | 0.026 | 2.74 | ring finger and CHY zinc finger domain containing 1 |
| NM_001033884 | Rexo4 | 0.032 | 3.30 | XPMC2 prevents mitotic catastrophe 2 homolog |
| NM_001013090 | Rg9mtd3 | 0.049 | 1.54 | RNA (guanine-9-) methyltransferase domain containing 3 |
| XM_224449 | RGD1304646_predicted | 0.011 | 1.71 | similar to KIAA1008 |
| NM_001034918 | RGD1304762 | 0.048 | 1.91 | IWS1 homolog |
| XM_001057949 | RGD1304890 | 0.003 | 1.55 | similar to Ubiquitin-fold modifier 1 precursor |
| XM_214589 | RGD1305072 | 0.009 | 2.55 | similar to CG8931-PA |
| NM_001025124 | RGD1305160 | 0.000 | 3.62 | N-terminal Asn amidase |
| NM_001030027 | RGD1307397 | 0.047 | 1.60 | similar to RIKEN cDNA 2810037C03 |
| NM_001014088 | RGD1307401 | 0.004 | 7.23 | hypothetical protein LOC315500 |
| XM_341397 | RGD1307583_predicted | 0.005 | 1.52 | similar to CG32736-PA, isoform A |
| XM_232819 | RGD1307658_predicted | 0.041 | 1.35 | similar to swan |
| NM_001013974 | RGD1308813 | 0.038 | 2.09 | hypothetical protein LOC303606 |
| XM_217114 | RGD1311744_predicted | 0.007 | 2.04 | similar to BCSC-1 |
| NM_019339 | Rgs12 | 0.004 | 2.11 | regulator of G-protein signalling 12 |
| NM_013224 | Rps26 | 0.011 | 5.84 | ribosomal protein S26 |
| NM_031113 | Rps27a | 0.003 | 27.79 | ribosomal protein S27a |
| NM_031985 | Rps6kb1 | 0.047 | 1.37 | ribosomal protein S6 kinase, polypeptide 1 |

| | | | | |
|---|---|---|---|---|
| XM_341851 | Rras_predicted | 0.020 | 1.95 | similar to Ras-related protein R-Ras |
| XM_227540 | Rsbn1_predicted | 0.024 | 1.39 | similar to rosbin, round spermatid basic protein 1 |
| NM_147146 | Rwdd1 | 0.015 | 1.87 | small androgen receptor-interacting protein |
| XM_343443 | Rwdd2_predicted | 0.030 | 4.91 | similar to RWD domain containing 2 |
| S68809 | S100a1 | 0.026 | 1.82 | S100 alpha |
| NM_053681 | S100a3 | 0.003 | 1.35 | S100 calcium binding protein A3 |
| XM_342174 | Scamp1 | 0.032 | 1.62 | similar to Secretory carrier-associated membrane protein 1 |
| NM_012647 | Scn2a1 | 0.000 | 10.38 | sodium channel, voltage-gated, type 2, alpha 1 polypeptide |
| NM_013119 | Scn3a | 0.034 | 1.44 | sodium channel, voltage-gated, type III, alpha polypeptide |
| NM_013082 | Sdc2 | 0.010 | 7.83 | syndecan 2 |
| NM_001005534 | Sdhc | 0.010 | 2.12 | succinate dehydrogenase complex, subunit C |
| XM_001060531 | Sgkl | 0.013 | 1.50 | similar to serum/glucocorticoid regulated kinase 3 |
| NM_080905 | Siah1a | 0.036 | 1.93 | seven in absentia 1A |
| NM_057121 | Slc15a1 | 0.018 | 1.79 | solute carrier family 15, member 1 isoform PepT1 |
| NM_031743 | Slc24a2 | 0.021 | 3.42 | solute carrier family 24, member 2 |
| NM_001014027 | Slc25a22 | 0.024 | 1.66 | solute carrier family 25, member 22 |
| NM_139100 | Slc25a3 | 0.029 | 1.33 | solute carrier family 25, member 3 |
| NM_031736 | Slc27a2 | 0.023 | 2.20 | solute carrier family 27, member 32 |
| NM_022252 | Slc33a1 | 0.011 | 1.59 | acetyl-CoA transporter |
| XM_213381 | Slc43a2_predicted | 0.016 | 2.08 | similar to solute carrier family 43, member 2 |
| NM_053521 | Slc5a7 | 0.043 | 1.77 | solute carrier family 5, member 7 |
| XM_237241 | Smarcal1_predicted | 0.003 | 10.24 | similar to SWI/SNF-related matrix-associated actin-dependent regulator of chromatin subfamily A-like protein 1 |
| XM_214621 | Snrpd1_predicted | 0.046 | 1.61 | similar to Small nuclear ribonucleoprotein Sm D1 |
| NM_199087 | Spint2 | 0.013 | 2.12 | serine protease inhibitor, Kunitz type 2 isoform b |
| NM_001034150 | Srpr | 0.024 | 1.43 | signal recognition particle receptor |
| NM_031119 | Ssb | 0.015 | 1.54 | Sjogren syndrome antigen B |
| NM_031122 | St13 | 0.013 | 2.17 | suppression of tumorigenicity 13 |
| NM_031735 | Stk3 | 0.037 | 1.51 | serine/threonine kinase 3 |
| NM_017166 | Stmn1 | 0.018 | 1.68 | stathmin 1 |
| NM_134378 | Sulf1 | 0.022 | 1.67 | sulfatase 1 |
| NM_001014263 | Sypl | 0.012 | 2.15 | synaptophysin-like protein |
| XM_001066806 | Taf6 | 0.038 | 8.72 | TAF6 RNA polymerase II, TATA box binding protein associated factor |
| XM_230631 | Tasp1_predicted | 0.042 | 5.73 | taspase, threonine aspartase 1 |
| NM_001008291 | Tfip11 | 0.017 | 2.31 | tuftelin interacting protein 11 |

| | | | | |
|---|---|---|---|---|
| XM_575591 | Tia1 | 0.005 | 1.95 | cytotoxic granule-associated RNA binding protein 1 (predicted) |
| XM_575591 | Tia1 | 0.031 | 1.35 | cytotoxic granule-associated RNA binding protein 1 (predicted) |
| NM_022592 | Tkt | 0.042 | 6.93 | transketolase |
| NM_001005554 | Tm9sf2 | 0.000 | 118.84 | transmembrane 9 superfamily member 2 |
| NM_021261 | Tmsb10 | 0.019 | 2.28 | thymosin, beta 10 |
| NM_012870 | Tnfrsf11b | 0.043 | 1.79 | tumor necrosis factor receptor superfamily, member 11b |
| BM389034 | Trap1 | 0.015 | 3.63 | UI-R-DZ0-cko-d-13-0-UI.s1 NCI_CGAP_DZ0 |
| NM_001009536 | Trim25_mapped | 0.015 | 3.09 | tripartite motif protein 25 |
| NM_053559 | Trpc6 | 0.041 | 2.19 | transient receptor protein 6 |
| XM_340973 | Ttc3_predicted | 0.027 | 1.34 | similar to tetratricopeptide repeat domain 3 |
| XM_216713 | Ttc6_predicted | 0.050 | 1.78 | similar to Tetratricopeptide repeat protein 6 |
| NM_183052 | Ube2v2 | 0.046 | 1.81 | ubiquitin-conjugating enzyme E2 variant 2 |
| XM_237865 | Usp42_predicted | 0.013 | 1.33 | similar to ubiquitin specific protease 42 |
| NM_198785 | Usp48 | 0.026 | 1.43 | ubiquitin specific protease 48 |
| NM_213563 | Vars2l | 0.009 | 9.59 | valyl-tRNA synthetase 2-like |
| NM_012686 | Vsnl1 | 0.024 | 1.70 | visinin-like 1 |
| NM_133581 | Wfdc1 | 0.021 | 3.06 | WAP four-disulfide core domain 1 |
| NM_199373 | Wrb | 0.010 | 3.59 | tryptophan rich basic protein |
| XM_216518 | XM_216518 | 0.050 | 1.73 | c-myc binding protein |
| XM_216838 | XM_216838 | 0.008 | 1.74 | testicular haploid expressed |
| XM_216940 | XM_216940 | 0.001 | 22.67 | ATPase, H+ transporting, V1 subunit C, isoform 1 |
| XM_228776 | XM_228776 | 0.001 | 1.52 | fusion (involved in t(12;16) in malignant liposarcoma) |
| XM_235185 | Xpot_predicted | 0.034 | 1.61 | similar to tRNA exportin |
| NM_177419 | Xrcc5 | 0.004 | 5.64 | X-ray repair complementing defective repair in Chinese hamster cells 5 |
| XM_233345 | Zcchc11_predicted | 0.001 | 23.20 | similar to zinc finger, CCHC domain containing 11 isoform c |
| XM_344780 | Zcchc14_predicted | 0.008 | 2.02 | similar to BDG-29 protein |
| XM_232972 | Zfp189_predicted | 0.001 | 1.76 | similar to zinc finger protein 189 |
| NM_001039020 | Zfp207 | 0.001 | 7.37 | zinc finger protein 207 |
| XM_230846 | Zswim1_predicted | 0.005 | 3.94 | similar to zinc finger, SWIM domain containing 1 |

**Supplementary Table 5: List of genes differentially downregulated in the mPFC transcriptome of adult PNFlx animals**

Six month old PNFlx animals and their age-matched controls were subjected to microarray analysis. The list of all the significantly downregulated genes is shown in the table ($p < 0.05$ ; Fold change < 0.74).

| Systematic Name | Gene Name | *P* Value | Fold change | Product |
|---|---|---|---|---|
| NM_012488 | A2m | 0.003 | 0.07 | alpha-2-macroglobulin |
| NM_013039 | Abcc8 | 0.032 | 0.64 | ATP-binding cassette, subfamily C member 8 |
| NM_021262 | Acp1 | 0.001 | 0.09 | acid phosphatase 1, soluble |
| NM_022681 | Adnp | 0.017 | 0.67 | activity-dependent neuroprotective protein |
| NM_138506 | Adra2c | 0.010 | 0.56 | adrenergic receptor, alpha 2c |
| NM_001009502 | Aer61 | 0.046 | 0.20 | glycosyltransferase Aer61 |
| NM_001002277 | Ahi1 | 0.025 | 0.71 | Abelson helper integration site 1 |
| NM_053781 | Akr1b7 | 0.017 | 0.17 | aldo-keto reductase family 1, member B7 |
| NM_013059 | Alpl | 0.045 | 0.52 | alkaline phosphatase, tissue-nonspecific |
| XM_001055725 | Ankrd15 | 0.019 | 0.10 | similar to ankyrin repeat domain 15 |
| AF039891 | Anpep | 0.032 | 0.62 | aminopeptidase N |
| NM_080478 | Apbb1 | 0.032 | 0.71 | amyloid beta (A4) precursor protein-binding, family B, member 1 |
| NM_012909 | Aqp2 | 0.033 | 0.63 | aquaporin 2 |
| NM_022960 | Aqp9 | 0.011 | 0.09 | aquaporin 9 |
| XM_236599 | Armc8_predicted | 0.006 | 0.10 | similar to armadillo repeat containing 8 |
| NM_001037767 | Arpc5l | 0.009 | 0.20 | actin related protein 2/3 complex, subunit 5-like |
| NM_012911 | Arrb2 | 0.030 | 0.11 | arrestin, beta 2 |
| XM_222785 | Astn1 | 0.037 | 0.67 | similar to astrotactin 1 |
| NM_053359 | Atox1 | 0.042 | 0.69 | ATX1 (antioxidant protein 1) homolog 1 |
| AW915889 | AW915889 | 0.047 | 0.34 | Bento Soares Rattus norvegicus cDNA clone RGIDC15 5- end |
| NM_022300 | Basp1 | 0.042 | 0.53 | brain abundant, membrane attached signal protein 1 |
| BE119829 | BE119829 | 0.004 | 0.44 | UI-R-CA0-ban-e-05-0-UI.s1 UI-R-CA0 Rattus norvegicus cDNA clone |
| BF389602 | BF389602 | 0.009 | 0.44 | UI-R-CJ0-bfd-f-04-0-UI.s1 UI-R-CJ0 Rattus norvegicus cDNA clone |
| XM_345190 | Bhlhb5_predicted | 0.020 | 0.18 | similar to basic helix-loop-helix domain containing, class B5 |
| NM_134413 | Btbd14b | 0.016 | 0.23 | BTB (POZ) domain containing 14B |
| XM_230616 | Btbd3_predicted | 0.044 | 0.15 | similar to BTB/POZ domain containing protein 3 isoform 1 |
| NM_080695 | Cacng7 | 0.003 | 0.73 | calcium channel, voltage-dependent, gamma subunit 7 |
| NM_144754 | Cant1 | 0.012 | 0.21 | calcium activated nucleotidase 1 |
| NM_001012164 | Cd97 | 0.002 | 0.18 | CD97 antigen |
| XM_238346 | Chchd3_predicted | 0.023 | 0.72 | similar to coiled-coil-helix-coiled-coil-helix domain containing 3 |
| NM_031701 | Cldn5 | 0.043 | 0.66 | claudin 5 |

| | | | | |
|---|---|---|---|---|
| NM_053753 | Clec4f | 0.001 | 0.65 | C-type lectin, superfamily member 13 |
| NM_012812 | Cox6a2 | 0.007 | 0.70 | cytochrome c oxidase, subunit VIa, polypeptide 2 |
| XM_233798 | Crim1_predicted | 0.039 | 0.08 | similar to cysteine rich transmembrane BMP regulator 1 |
| NM_138518 | Crispld2 | 0.027 | 0.08 | late gestation lung protein 1 |
| NM_053955 | Crym | 0.009 | 0.38 | crystallin, mu |
| NM_033233 | Csh1l1 | 0.001 | 0.09 | chorionic somatomammotropin hormone 1 variant |
| NM_053615 | Csnk1a1 | 0.029 | 0.70 | casein kinase 1, alpha 1 |
| XM_215451 | Cspg2 | 0.001 | 0.10 | similar to Versican core protein precursor |
| XM_216386 | Ctnnal1_predicted | 0.016 | 0.04 | similar to Alpha-catulin |
| NM_181087 | Cyp26b1 | 0.046 | 0.12 | cytochrome P450, family 26, subfamily b, polypeptide 1 |
| NM_001011978 | Dhdds | 0.042 | 0.14 | dehydrodolichyl diphosphate synthase |
| XM_213964 | Disp1_predicted | 0.001 | 0.10 | similar to dispatched homolog 1 |
| NM_133419 | Dkc1 | 0.025 | 0.24 | dyskeratosis congenita 1, dyskerin |
| NM_138519 | Dkk3 | 0.002 | 0.01 | dickkopf homolog 3 |
| NM_001015021 | Dnajb11 | 0.046 | 0.17 | DnaJ (Hsp40) homolog, subfamily B, member 11 |
| NM_001013196 | Dnajc4 | 0.017 | 0.73 | DnaJ (Hsp40) homolog, subfamily C, member 4 |
| NM_001025650 | Dusp11 | 0.047 | 0.70 | dual specificity phosphatase 11 |
| NM_133578 | Dusp5 | 0.001 | 0.07 | dual specificity phosphatase 5 |
| XM_216563 | Eif4g3_predicted | 0.010 | 0.54 | similar to Eukaryotic translation initiation factor 4 gamma 3 |
| NM_199394 | Entpd5 | 0.002 | 0.05 | ectonucleoside triphosphate diphosphohydrolase 5 |
| BC071175 | Ero1l | 0.026 | 0.75 | ERO1-like |
| XM_343055 | Etv1_predicted | 0.000 | 0.03 | similar to ets variant gene 1 |
| NM_053511 | Fbxo2 | 0.014 | 0.71 | F-box only protein 2 |
| NM_031066 | Fez1 | 0.015 | 0.65 | fasciculation and elongation protein zeta 1 |
| NM_012954 | Fosl2 | 0.015 | 0.16 | fos-like antigen 2 |
| XM_220287 | Foxi1_predicted | 0.003 | 0.49 | similar to forkhead box I1 |
| NM_053895 | Frag1 | 0.004 | 0.03 | FGF receptor activating protein 1 |
| NM_022706 | Gabarapl2 | 0.021 | 0.52 | GABA(A) receptor-associated protein like 2 |
| NM_031802 | Gabbr2 | 0.030 | 0.43 | G protein-coupled receptor 51 |
| NM_017295 | Gabra5 | 0.036 | 0.15 | gamma-aminobutyric acid A receptor, alpha 5 |
| NM_031733 | Gabrq | 0.004 | 0.42 | gamma-aminobutyric acid A receptor, theta |
| NM_001008321 | Gadd45b | 0.041 | 0.74 | growth arrest and DNA-damage-inducible 45 beta |
| XM_231109 | Gfi1b_predicted | 0.039 | 0.55 | similar to growth factor independent 1B |
| NM_019280 | Gja5 | 0.041 | 0.13 | gap junction membrane channel protein alpha 5 |
| NM_001025736 | Gnl2 | 0.000 | 0.01 | guanine nucleotide binding protein-like 2 |
| NM_053403 | Grb7 | 0.008 | 0.14 | growth factor receptor bound protein 7 |
| NM_053807 | Gripap1 | 0.023 | 0.68 | GRIP1 associated protein 1 |
| NM_017012 | Grm5 | 0.007 | 0.26 | glutamate receptor, metabotropic 5 |
| XM_001060252 | Gs3 | 0.000 | 0.04 | putative regulation protein GS3 |

| | | | | |
|---|---|---|---|---|
| XM_217299 | Gup1_predicted | 0.005 | 0.52 | similar to Gup1, glycerol uptake/transporter homolog |
| XM_217299 | Gup1_predicted | 0.000 | 0.57 | similar to Gup1, glycerol uptake/transporter homolog |
| NM_153468 | Gzma | 0.045 | 0.53 | granzyme A |
| NM_019189 | Hapln1 | 0.029 | 0.12 | hyaluronan and proteoglycan link protein 1 |
| NM_053375 | Hcn1 | 0.003 | 0.04 | hyperpolarization-activated, cyclic nucleotide-gated potassium channel 1 |
| NM_080896 | Hnrph1 | 0.035 | 0.12 | heterogeneous nuclear ribonucleoprotein H1 |
| XM_001061027 | Hsf1 | 0.048 | 0.73 | similar to Heat shock factor protein 1 (HSF 1) |
| XM_344130 | Inhbb | 0.044 | 0.75 | similar to Inhibin beta B chain precursor |
| NM_019311 | Inpp5d | 0.043 | 0.66 | inositol polyphosphate-5-phosphatase D |
| XM_230604 | Itpa_mapped | 0.022 | 0.14 | similar to Inosine triphosphate pyrophosphatase |
| XM_573259 | Itsn1 | 0.000 | 0.08 | similar to Intersectin-1 |
| U70050 | Jag2 | 0.041 | 0.35 | jagged2 precursor |
| NM_031514 | Jak2 | 0.002 | 0.09 | Janus kinase 2 |
| NM_012855 | Jak3 | 0.035 | 0.29 | Janus kinase 3 |
| NM_138875 | Jund | 0.045 | 0.49 | Jun D proto-oncogene |
| NM_017099 | Kcnj8 | 0.000 | 0.75 | potassium inwardly-rectifying channel, subfamily J, member 8 |
| NM_021688 | Kcnk1 | 0.035 | 0.67 | potassium channel, subfamily K, member 1 |
| XM_344237 | Kctd8_predicted | 0.006 | 0.60 | similar to potassium channel tetramerisation domain containing 8 |
| XM_235478 | Kdelr3_predicted | 0.046 | 0.10 | similar to ER lumen protein retaining receptor 3 |
| NM_022249 | Khdrbs3 | 0.001 | 0.08 | etoile, Sam68-like protein SLM-2 |
| NM_213626 | kif13B | 0.035 | 0.61 | kinesin 13B |
| NM_001011910 | Lap3 | 0.003 | 0.13 | leucine aminopeptidase 3 |
| NM_017024 | Lcat | 0.038 | 0.68 | lecithin cholesterol acyltransferase |
| NM_145769 | Lgi1 | 0.000 | 0.03 | leucine-rich, glioma inactivated 1 |
| NM_001002016 | Lmna | 0.012 | 0.07 | lamin A isoform C2 |
| NM_139255 | LOC246120 | 0.005 | 0.19 | hypothetical protein LOC246120 |
| NM_147136 | LOC257642 | 0.002 | 0.45 | rRNA promoter binding protein |
| NM_173317 | LOC286983 | 0.001 | 0.18 | putative pheromone receptor |
| XM_001069678 | LOC501046 | 0.044 | 0.67 | similar to Phakinin |
| XM_001056340 | LOC679437 | 0.029 | 0.12 | similar to solute carrier family 25, member 28 |
| XM_001055358 | LOC680021 | 0.003 | 0.71 | similar to glyoxylate reductase/hydroxypyruvate reductase |
| XM_001073780 | LOC689397 | 0.006 | 0.19 | similar to C184L-22 |
| XM_001075711 | LOC690810 | 0.009 | 0.18 | similar to adenosine deaminase, tRNA-specific 1 |
| NM_134408 | Lphn2 | 0.000 | 0.05 | latrophilin 2 |
| NM_031342 | Lypla2 | 0.032 | 0.52 | lysophospholipase 2 |

| | | | | |
|---|---|---|---|---|
| NM_019318 | Maf | 0.015 | 0.16 | v-maf musculoaponeurotic fibrosarcoma oncogene homolog |
| NM_001031655 | Manba | 0.005 | 0.39 | mannosidase, beta A, lysosomal |
| NM_012757 | Mas1 | 0.031 | 0.72 | MAS1 oncogene |
| XM_342750 | Mfap5_predicted | 0.002 | 0.19 | similar to Microfibrillar-associated protein 5 precursor |
| NM_138976 | Mfn1 | 0.006 | 0.07 | mitofusin 1 |
| NM_134459 | Mic2l1 | 0.009 | 0.57 | MIC2 like 1 |
| BC086585 | Mki67ip | 0.012 | 0.05 | Unknown (protein for MGC:105693) |
| NM_001007637 | mrpl24 | 0.010 | 0.66 | mitochondrial ribosomal protein L24 |
| NM_030995 | Mtap1a | 0.019 | 0.08 | microtubule-associated protein 1 A |
| XM_215769 | Mtch2_predicted | 0.012 | 0.21 | similar to Mitochondrial carrier homolog 2 |
| XM_238555 | Nckipsd_predicted | 0.000 | 0.01 | similar to NCK interacting protein with SH3 domain |
| XM_216859 | Ndufa7_predicted | 0.013 | 0.10 | similar to NADH dehydrogenase 1 alpha subcomplex, 7 |
| NM_054010 | Neu3 | 0.012 | 0.20 | neuraminidase 3 |
| NM_012865 | Nfya | 0.010 | 0.25 | nuclear transcription factor-Y alpha |
| XM_218718 | Nipa2_predicted | 0.003 | 0.09 | similar to non-imprinted in Prader-Willi/Angelman syndrome 2 |
| NM_020087 | Notch3 | 0.008 | 0.24 | Notch gene homolog 3 |
| NM_145775 | Nr1d1 | 0.013 | 0.70 | nuclear receptor subfamily 1, group D, member 1 |
| NM_012993 | Nrd1 | 0.011 | 0.16 | n-arginine dibasic convertase 1 |
| NM_207607 | Ns5atp4 | 0.002 | 0.44 | NS5A (hepatitis C virus) transactivated protein 4 |
| NM_001000519 | Olr1307 | 0.048 | 0.43 | olfactory receptor Olr1307 |
| NM_001000519 | Olr1307 | 0.049 | 0.65 | olfactory receptor Olr1307 |
| NM_001001108 | Olr1548_predicted | 0.035 | 0.19 | olfactory receptor Olr1548 |
| XM_221921 | Olr1572_predicted | 0.016 | 0.53 | similar to olfactory receptor Olr1768 |
| NM_001000694 | Olr16_predicted | 0.031 | 0.42 | olfactory receptor Olr16 |
| NM_001001028 | Olr174_predicted | 0.046 | 0.35 | olfactory receptor Olr174 |
| NM_001000205 | Olr230_predicted | 0.022 | 0.60 | olfactory receptor Olr230 |
| NM_001000672 | Olr535_predicted | 0.043 | 0.23 | olfactory receptor Olr535 |
| NM_001000663 | Olr581_predicted | 0.042 | 0.20 | olfactory receptor Olr581 |
| NM_001000926 | Olr611_predicted | 0.021 | 0.09 | olfactory receptor Olr611 |
| NM_001000608 | Olr776_predicted | 0.010 | 0.09 | olfactory receptor Olr776 |
| NM_001000600 | Olr796_predicted | 0.049 | 0.37 | olfactory receptor Olr796 |
| NM_001000842 | Olr821_predicted | 0.009 | 0.48 | olfactory receptor Olr821 |
| NM_057143 | Park7 | 0.028 | 0.61 | DJ-1 protein |

| | | | | |
|---|---|---|---|---|
| XM_001055332 | Pctk2 | 0.008 | 0.01 | similar to Serine/threonine-protein kinase PCTAIRE-2 |
| NM_022265 | Pdcd4 | 0.009 | 0.03 | programmed cell death 4 |
| NM_001006966 | Peci | 0.022 | 0.28 | peroxisomal delta3, delta2-enoyl-Coenzyme A isomerase |
| NM_057125 | Pex6 | 0.003 | 0.08 | peroxisomal biogenesis factor 6 |
| NM_013190 | Pfkl | 0.001 | 0.25 | phosphofructokinase, liver, B-type |
| XM_217188 | Pias1_predicted | 0.003 | 0.14 | similar to protein inhibitor of activated STAT 1 |
| NM_001012207 | Pigc | 0.012 | 0.75 | phosphatidylinositol glycan, class C |
| NM_001008316 | Plag1 | 0.014 | 0.31 | pleiomorphic adenoma gene 1 |
| NM_172031 | Plunc | 0.004 | 0.19 | palate, lung, and nasal epithelium carcinoma associated protein precursor |
| M57728 | Pmpca | 0.023 | 0.74 | general mitochondrial matrix processing protease 55 kDa subunit |
| NM_001004259 | Pnkp | 0.000 | 0.05 | polynucleotide kinase 3 ' -phosphatase |
| NM_053322 | Pom210 | 0.004 | 0.06 | nuclear pore membrane glycoprotein 210 |
| NM_022951 | Ppp1r10 | 0.007 | 0.68 | protein phosphatase 1, regulatory subunit 10 |
| NM_184051 | Prkag2 | 0.008 | 0.12 | AMP-activated protein kinase gamma2 subunit |
| XM_343046 | Prkar2b | 0.013 | 0.16 | similar to cAMP-dependent protein kinase type II-beta regulatory subunit |
| XM_238534 | Prkcsh_predicted | 0.027 | 0.73 | similar to Glucosidase II beta subunit precursor |
| NM_138882 | Pspla1 | 0.007 | 0.14 | phosphatidylserine-specific phospholipase A1 |
| NM_172223 | Pxmp4 | 0.008 | 0.60 | peroxisomal membrane protein 4 |
| XM_342542 | Pygb | 0.003 | 0.11 | similar to Glycogen phosphorylase, brain form |
| M83679 | Rab15 | 0.029 | 0.61 | RAB15 |
| NM_053821 | Ralb | 0.012 | 0.09 | v-ral simian leukemia viral oncogene homolog B |
| XM_001079347 | Rapgef1 | 0.006 | 0.22 | similar to Rap guanine nucleotide exchange factor 1 isoform 3 |
| XM_226421 | Rbm35b_predicted | 0.022 | 0.62 | similar to fusilli CG8205-PD, isoform D |
| XM_226421 | Rbm35b_predicted | 0.033 | 0.75 | similar to fusilli CG8205-PD, isoform D |
| NM_172077 | Reg3a | 0.007 | 0.65 | regenerating islet-derived 3 alpha |
| XM_217550 | RGD1304704 | 0.040 | 0.73 | hypothetical protein LOC302247 |
| NM_001013976 | RGD1305007 | 0.009 | 0.61 | hypothetical protein LOC303749 |
| XM_234484 | RGD1309059_predicted | 0.002 | 0.16 | similar to falafel CG9351-PA, isoform A |
| XM_215112 | RGD1309350_predicted | 0.043 | 0.18 | similar to Hypothetical transthyretin-like protein R09H10.3 |
| XM_219477 | RGD1310052_predicted | 0.001 | 0.13 | similar to beta1,4-N-acetylgalactosaminyltransferase IV |
| XM_001076059 | RGD1311612_predicted | 0.010 | 0.05 | similar to CG8257-PA |

| | | | | |
|---|---|---|---|---|
| NM_001013982 | RGD1311873 | 0.044 | 0.66 | hypothetical protein LOC304496 |
| NM_199493 | RGD735029 | 0.022 | 0.74 | hypothetical protein LOC307480 |
| NM_019341 | Rgs5 | 0.011 | 0.13 | regulator of G-protein signaling 5 |
| NM_001013222 | Rnd1 | 0.002 | 0.29 | Rho family GTPase 1 |
| NM_021582 | Rpa2 | 0.025 | 0.07 | replication protein A2 |
| NM_133591 | Rph3al | 0.014 | 0.46 | rabphilin 3A-like |
| NM_017151 | Rps15 | 0.030 | 0.53 | ribosomal protein S15 |
| NM_053982 | Rps15a | 0.001 | 0.07 | ribosomal protein S15a |
| XM_234419 | Rps6kl1_predicted | 0.019 | 0.71 | similar to ribosomal protein S6 kinase-like 1 |
| NM_053471 | Sema6b | 0.009 | 0.15 | sema domain, transmembrane domain, and cytoplasmic domain |
| XM_243863 | Sema7a_predicted | 0.007 | 0.47 | similar to Semaphorin-7A precursor |
| NM_145097 | Serpina4 | 0.048 | 0.68 | serine (or cysteine) proteinase inhibitor, clade A, member 4 |
| NM_001011920 | Sf4 | 0.042 | 0.71 | splicing factor 4 |
| XM_238127 | Sh2bp1 | 0.006 | 0.65 | similar to SH2 domain binding protein 1 |
| NM_022866 | Slc13a3 | 0.032 | 0.75 | solute carrier family 13 member 3 |
| NM_198760 | Slc16a6 | 0.016 | 0.19 | solute carrier family 16, member 6 |
| NM_031743 | Slc24a2 | 0.001 | 0.02 | solute carrier family 24, member 2 |
| NM_053965 | Slc25a20 | 0.043 | 0.59 | solute carrier family 25, member 20 |
| XM_221940 | Slc29a4_predicted | 0.040 | 0.47 | similar to solute carrier family 29, member 4 |
| XM_235840 | Slc36a4_predicted | 0.036 | 0.52 | similar to proton/amino acid transporter 4 |
| NM_145776 | Slc38a3 | 0.024 | 0.71 | solute carrier family 38, member 3 |
| NM_177481 | Slco3a1 | 0.036 | 0.69 | solute carrier organic anion transporter family, member 3a1 |
| XM_341132 | Smg7_predicted | 0.000 | 0.05 | similar to SMG-7 homolog |
| XM_001080400 | Sos2 | 0.001 | 0.06 | similar to son of sevenless homolog 2 |
| XM_225864 | Spire1_predicted | 0.030 | 0.63 | similar to spire homolog 1 |
| NM_053464 | Srm | 0.004 | 0.37 | spermidine synthase |
| NM_133522 | Sstr3 | 0.009 | 0.68 | somatostatin receptor 3 |
| NM_019123 | St6galnac3 | 0.045 | 0.71 | sialyltransferase 7c |
| XM_214712 | Sult5a1_predicted | 0.010 | 0.16 | similar to sulfotransferase family 5A, member 1 |
| XM_223981 | Supt16h_predicted | 0.043 | 0.25 | similar to suppressor of Ty 16 homolog |
| NM_021695 | Synpo | 0.050 | 0.52 | synaptopodin |
| NM_001012161 | Tbce | 0.031 | 0.42 | tubulin-specific chaperone e |
| NM_031129 | Tceb2 | 0.020 | 0.03 | transcription elongation factor B (SIII), polypeptide 2 |
| XM_217232 | Tcfdp2_predicted | 0.048 | 0.21 | similar to transcription factor Dp-2 |
| NM_138871 | Tdrd7 | 0.029 | 0.74 | tudor domain containing 7 |

| | | | | |
|---|---|---|---|---|
| NM_199098 | Tmem19 | 0.020 | 0.29 | transmembrane protein 19 |
| XM_215641 | Tmod4_predicted | 0.000 | 0.47 | similar to Tropomodulin-4 |
| XM_345932 | Tmprss4_predicted | 0.017 | 0.43 | similar to transmembrane protease, serine 4 |
| NM_001014039 | Tnfaip8l2 | 0.032 | 0.63 | tumor necrosis factor, alpha-induced protein 8-like 2 |
| NM_013091 | Tnfrsf1a | 0.009 | 0.13 | tumor necrosis factor receptor superfamily, member 1a |
| NM_031357 | Tpp1 | 0.038 | 0.69 | tripeptidyl-peptidase I |
| NM_019322 | Tpsab1 | 0.040 | 0.57 | tryptase alpha/beta 1 |
| NM_053867 | Tpt1 | 0.006 | 0.18 | tumor protein, translationally-controlled 1 |
| XM_225947 | Trim36_predicted | 0.025 | 0.18 | similar to haprin |
| NM_021762 | Tsn | 0.023 | 0.18 | translin |
| NM_031614 | Txnrd1 | 0.040 | 0.15 | thioredoxin reductase 1 |
| NM_012682 | Ucp1 | 0.011 | 0.26 | uncoupling protein 1 |
| XM_344168 | Vamp4_predicted | 0.002 | 0.05 | similar to Vesicle-associated membrane protein 4 |
| NM_012889 | Vcam1 | 0.025 | 0.15 | vascular cell adhesion molecule 1 |
| NM_031836 | Vegfa | 0.047 | 0.72 | vascular endothelial growth factor A |
| XM_214669 | Wwp2_predicted | 0.013 | 0.75 | similar to WW domain-containing protein 2 |
| XM_221138 | XM_221138 | 0.000 | 0.07 | SEC14-like 1 |
| XM_342024 | XM_342024 | 0.038 | 0.60 | damage-specific DNA binding protein 1 |
| XM_230639 | Zfp339_predicted | 0.004 | 0.75 | similar to ovo-like 2 isoform A |

**Supplementary Table 6: List of genes differentially upregulated in the mPFC transcriptome of adult JFlx animals.**

Six month old JFlx animals and their age-matched controls were subjected to microarray analysis. The list of all the significantly upregulated genes is shown in the table ($p < 0.05$ ; Fold change > 1.35) .

| Systematic Name | GeneName | *P* Value | Fold Change (Log 2) | Product |
|---|---|---|---|---|
| XM_241525 | Abca4 | 0.0306 | 2.60 | similar to Retinal-specific ATP-binding cassette transporter |
| NM_013084 | Acadsb | 0.0033 | 1.38 | acyl-Coenzyme A dehydrogenase, short/branched chain |
| XM_340916 | Adam11 | 0.0002 | 1.69 | similar to ADAM 11 precursor |
| NM_031006 | Adar | 0.0004 | 1.45 | adenosine deaminase, RNA-specific |
| XM_223616 | Adcy1 | 0.0113 | 1.48 | similar to adenylate cyclase 1 |
| NM_133511 | Adcyap1r1 | 0.0034 | 1.55 | adenylate cyclase activating polypeptide 1 receptor 1 |
| NM_031552 | Add3 | 0.0169 | 1.36 | adducin 3 (gamma) |
| AY724514 | Adipor2 | 0.0043 | 1.61 | adiponectin receptor 2 |
| NM_019220 | Aes | 0.0008 | 1.69 | amino-terminal enhancer of split |
| AF468681 | AF468681 | 0.0136 | 1.42 | helix-loop-helix protein |
| NM_001002277 | Ahi1 | 0.0173 | 1.34 | Abelson helper integration site 1 |
| NM_001002277 | Ahi1 | 0.0009 | 1.43 | Abelson helper integration site 1 |
| NM_017135 | Ak3l1 | 0.0088 | 1.42 | adenylate kinase 3-like 1 |
| NM_031753 | Alcam | 0.0016 | 1.39 | activated leukocyte cell adhesion molecule |
| XM_343457 | Amotl2 | 0.0399 | 1.78 | similar to Angiomotin-like protein 2 |
| NM_053714 | Ank | 0.0277 | 1.36 | progressive ankylosis homolog |
| NM_017277 | Ap1b1 | 0.0016 | 1.32 | adaptor protein complex AP-1, beta 1 subunit |
| NM_001007005 | Arhgdia | 0.0026 | 1.77 | Rho GDP dissociation inhibitor (GDI) alpha |
| XM_224588 | Arhgef3 | 0.0272 | 1.55 | similar to Rho guanine nucleotide exchange factor 3 |
| NM_053740 | Arhgef7 | 0.0290 | 1.37 | Rho guanine nucleotide exchange factor 7 |
| NM_031018 | Atf2 | 0.0015 | 1.96 | activating transcription factor 2 |
| XM_345102 | Atoh7 | 0.0162 | 1.77 | similar to atonal homolog 7 |
| NM_001005871 | Atp2b4 | 0.0004 | 1.45 | plasma membrane calcium ATPase 4 |
| NM_031535 | Bcl2l1 | 0.0012 | 1.80 | Bcl2-like 1 isoform 1 |
| XM_232252 | Bcl2l13 | 0.0427 | 1.54 | similar to Bcl-2-like 13 protein |
| XM_214967 | Bclaf1 | 0.0036 | 1.43 | BCL2-associated transcription factor 1 |
| NM_017178 | Bmp2 | 0.0398 | 1.37 | bone morphogenetic protein 2 |
| XM_230952 | Cables2 | 0.0021 | 1.33 | hypothetical protein |
| NM_133529 | Cabp1 | 0.0009 | 1.36 | calcium binding protein 1 isoform 1 |
| NM_001007730 | Cabp7 | 0.0115 | 1.65 | calcium binding protein 7 |
| AB056125 | Camk2a | 0.0278 | 1.54 | calcium/calmodulin-dependent protein kinase II alpha |
| NM_172008 | Canx | 0.0319 | 1.51 | calnexin |

| | | | | |
|---|---|---|---|---|
| NM_147144 | Casc3 | 0.0094 | 1.46 | metastatic lymph node 51 |
| NM_053736 | Casp4 | 0.0496 | 1.43 | caspase 11 |
| NM_131914 | Cav2 | 0.0167 | 1.63 | caveolin 2 |
| XM_223270 | Ccng2 | 0.0089 | 1.42 | similar to Cyclin-G2 |
| XM_223083 | Cd34 | 0.0422 | 2.18 | similar to CD34 antigen |
| NM_053983 | Cd52 | 0.0439 | 1.45 | CD52 antigen |
| XM_228203 | Cdc2l6 | 0.0121 | 1.42 | similar to Cell division cycle 2-like protein kinase 6 |
| AB010437 | Cdh8 | 0.0151 | 1.41 | cadherin-8 |
| NM_001025682 | Cdr2 | 0.0371 | 1.46 | cerebellar degeneration-related 2 |
| NM_053643 | Cds2 | 0.0495 | 1.57 | phosphatidate cytidylyltransferase 2 |
| NM_133567 | Centa1 | 0.0032 | 1.74 | centaurin, alpha 1 |
| XM_576238 | Centg1 | 0.0058 | 1.32 | similar to Centaurin-gamma 1 |
| XM_237381 | Centg2 | 0.0199 | 1.51 | similar to centaurin, gamma 2 isoform 2 |
| XM_220602 | Chd3 | 0.0011 | 2.08 | similar to chromodomain helicase DNA binding protein 3 isoform 3 |
| NM_019297 | Chrnb2 | 0.0004 | 1.77 | neuronal nicotinic acetylcholine receptor beta 2 |
| XM_235007 | Chst11 | 0.0306 | 1.46 | similar to Carbohydrate sulfotransferase 11 |
| NM_001029911 | Cit | 0.0038 | 1.59 | citron |
| XM_243040 | Clstn1 | 0.0001 | 1.44 | similar to calsyntenin 1 |
| XM_345143 | Col4a3bp | 0.0146 | 1.38 | similar to procollagen, type IV, alpha 3 binding protein |
| XM_341227 | Cpeb2 | 0.0113 | 1.39 | similar to cytoplasmic polyadenylation element binding protein 2 isoform A |
| NM_022864 | Cplx1 | 0.0045 | 1.63 | complexin 1 |
| NM_019302 | Crk | 0.0041 | 1.51 | v-crk sarcoma virus CT10 oncogene homolog |
| NM_022288 | Csnk1g1 | 0.0498 | 1.34 | casein kinase 1, gamma 1 |
| NM_053824 | Csnk2a1 | 0.0059 | 1.40 | casein kinase II, alpha 1 polypeptide |
| XM_232077 | Ctnna2 | 0.0113 | 1.42 | similar to Alpha-2 catenin |
| XM_228172 | Dcbld1 | 0.0005 | 2.25 | similar to Discoidin, CUB and LCCL domain-containing protein 1 precursor |
| XM_235480 | Ddx17 | 0.0023 | 1.47 | similar to DEAD box polypeptide 17 isoform p82 |
| XM_216452 | Dhcr24 | 0.0065 | 1.33 | 24-dehydrocholesterol reductase |
| XM_345790 | Diras1 | 0.0049 | 1.56 | similar to DIRAS family, GTP-binding RAS-like 1 |
| XM_345790 | Diras1 | 0.0082 | 1.71 | similar to DIRAS family, GTP-binding RAS-like 1 |
| XM_225214 | Diras2 | 0.0214 | 1.66 | similar to GTP-binding protein Di-Ras2 |
| NM_019621 | Dlgh4 | 0.0001 | 1.82 | postsynaptic density protein 95 |
| XM_343462 | Dnajc13 | 0.0090 | 1.37 | similar to DnaJ (Hsp40) homolog, subfamily C, member 13 |
| NM_022850 | Dpp6 | 0.0003 | 1.45 | dipeptidylpeptidase 6 |
| XM_218514 | Dpy19l3 | 0.0162 | 1.57 | similar to dpy-19-like 3 |
| NM_019226 | Dync1h1 | 0.0125 | 1.44 | dynein, cytoplasmic, heavy chain 1 |
| XM_233986 | E2f6 | 0.0103 | 1.35 | similar to E2F transcription factor 6 |
| NM_001002815 | Ece2 | 0.0482 | 1.41 | endothelin-converting enzyme 2 |

| | | | | |
|---|---|---|---|---|
| XM_341803 | Egfl4 | 0.0175 | 1.47 | similar to EGF-like domain-containing protein 4 |
| XM_233544 | Eif2c1 | 0.0478 | 1.36 | similar to eukaryotic translation initiation factor 2C, 1 |
| XM_001078496 | Elk1 | 0.0018 | 1.46 | similar to ETS domain-containing protein Elk-1 |
| XM_001078496 | Elk1 | 0.0031 | 1.64 | similar to ETS domain-containing protein Elk-1 |
| XM_001078496 | Elk1 | 0.0048 | 1.68 | similar to ETS domain-containing protein Elk-1 |
| XM_001078496 | Elk1 | 0.0013 | 1.82 | similar to ETS domain-containing protein Elk-1 |
| XM_001078496 | Elk1 | 0.0040 | 1.85 | similar to ETS domain-containing protein Elk-1 |
| XM_001078496 | Elk1 | 0.0122 | 1.85 | similar to ETS domain-containing protein Elk-1 |
| XM_001078496 | Elk1 | 0.0054 | 1.88 | similar to ETS domain-containing protein Elk-1 |
| XM_001078496 | Elk1 | 0.0022 | 2.08 | similar to ETS domain-containing protein Elk-1 |
| XM_231650 | Ephb6 | 0.0338 | 1.52 | similar to Ephrin type-B receptor 6 precursor |
| NM_017003 | Erbb2 | 0.0442 | 1.74 | v-erb-b2 erythroblastic leukemia viral oncogene homolog 2 |
| XM_232515 | Etnk1 | 0.0326 | 1.46 | similar to Ethanolamine kinase 1 (EKI 1) |
| XM_239510 | Ets2 | 0.0050 | 1.67 | similar to C-ets-2 protein |
| XM_217715 | Eya4 | 0.0342 | 1.34 | similar to Eyes absent homolog 4 |
| NM_031344 | Fads2 | 0.0336 | 1.78 | fatty acid desaturase 2 |
| NM_001012739 | Fau | 0.0052 | 1.33 | Finkel-Biskis-Reilly murine sarcoma virus ubiquitously expressed |
| XM_341091 | Fbxo21 | 0.0048 | 1.62 | similar to F-box only protein 21 |
| XM_001058601 | Fmn2 | 0.0028 | 1.69 | similar to Formin-2 |
| NM_012561 | Fst | 0.0343 | 1.34 | follistatin |
| XM_225228 | Gfod1 | 0.0069 | 1.47 | similar to glucose-fructose oxidoreductase domain containing 1 |
| NM_031034 | Gna12 | 0.0195 | 1.52 | guanine nucleotide binding protein (G protein) alpha 12 |
| NM_031034 | Gna12 | 0.0171 | 1.57 | guanine nucleotide binding protein (G protein) alpha 12 |
| AF189020 | Gnai2 | 0.0142 | 1.48 | guanine nucleotide binding protein (G protein), alpha inhibiting polypeptide 2 |
| NM_031036 | Gnaq | 0.0386 | 1.36 | guanine nucleotide binding protein, alpha q polypeptide |
| NM_017274 | Gpam | 0.0302 | 1.40 | glycerol-3-phosphate acyltransferase, mitochondrial |
| NM_001012185 | Gpiap1 | 0.0119 | 1.32 | GPI-anchored membrane protein 1 |
| XM_225625 | Gpr158 | 0.0181 | 1.38 | similar to G protein-coupled receptor 158 isoform a |
| XM_215095 | Gprc5b | 0.0154 | 1.55 | similar to G protein-coupled receptor, family C, group 5, member B |
| NM_017344 | Gsk3a | 0.0001 | 2.98 | glycogen synthase kinase 3 alpha |
| NM_022285 | Hapln2 | 0.0286 | 1.34 | hyaluronan and proteoglycan link protein 2 |
| XM_238362 | Hdac11 | 0.0011 | 1.48 | similar to histone deacetylase 11 |
| NM_139327 | Hmga1 | 0.0158 | 1.35 | high mobility group AT-hook 1 |

| | | | | |
|---|---|---|---|---|
| NM_053310 | Homer3 | 0.0298 | 1.32 | homer homolog 3 |
| XM_222017 | Hrbl | 0.0254 | 1.52 | similar to HIV-1 Rev-binding protein-like protein isoform 2 |
| XM_237060 | Hs6st1 | 0.0043 | 1.48 | similar to heparan sulfate 6-O-sulfotransferase 1 |
| XM_217648 | Hspa12a | 0.0033 | 1.38 | similar to heat shock protein 12A |
| NM_139260 | Il3ra | 0.0129 | 1.42 | interleukin 3 receptor, alpha chain |
| XM_233441 | Jmjd2a | 0.0008 | 1.65 | similar to Jumonji domain-containing protein 2A |
| NM_001003711 | Jph4 | 0.0001 | 1.37 | junctophilin 4 |
| NM_017303 | Kcnab1 | 0.0335 | 1.54 | potassium voltage-gated channel, shaker-related subfamily, beta 1 |
| NM_012856 | Kcnc1 | 0.0079 | 1.36 | potassium voltage gated channel, Shaw-related subfamily, 1 |
| XM_001072764 | Kcng1 | 0.0117 | 1.34 | similar to Potassium voltage-gated channel subfamily G 1 |
| NM_031602 | Kcnj10 | 0.0050 | 1.67 | potassium inwardly-rectifying channel J10 |
| XM_241981 | Kif5c | 0.0079 | 1.54 | similar to kinesin family member 5C |
| NM_001015029 | Kpna6 | 0.0196 | 1.80 | karyopherin alpha 6 |
| NM_001011910 | Lap3 | 0.0342 | 1.53 | leucine aminopeptidase 3 |
| NM_021851 | Lin7c | 0.0094 | 1.34 | lin-7 homolog C |
| XM_217385 | Lman2l | 0.0087 | 1.44 | similar to VIP36-like protein precursor |
| XM_237217 | LOC316457 | 0.0341 | 1.35 | similar to phosphatidylinositol 5-kinase, type III isoform 2 |
| XM_340893 | LOC360618 | 0.0397 | 1.49 | hypothetical protein LOC360618 |
| XM_575106 | LOC499768 | 0.0081 | 1.50 | similar to TBC1 domain family, member 13 |
| XM_001057149 | LOC680430 | 0.0181 | 1.32 | similar to germinal histone H4 gene |
| XM_001058014 | LOC680616 | 0.0003 | 1.91 | similar to Protein phosphatase 1 regulatory inhibitor subunit 16B |
| XM_001056512 | LOC681927 | 0.0435 | 1.35 | similar to SEC24 related gene family, member C isoform 4 |
| XM_001063618 | LOC685393 | 0.0031 | 2.46 | similar to lin-9 homolog |
| XM_001076463 | LOC691002 | 0.0167 | 1.66 | similar to dihydrodiol dehydrogenase (dimeric) |
| XM_237521 | Lpin2 | 0.0406 | 1.42 | similar to lipin 2 |
| XM_001055209 | Lrp1b | 0.0132 | 1.85 | similar to low density lipoprotein-related protein 1B |
| NM_001012062 | Map3k7ip2 | 0.0169 | 1.79 | mitogen-activated protein kinase kinase kinase 7 interacting protein 2 |
| XM_225726 | Mapk4 | 0.0005 | 2.10 | similar to Mitogen-activated protein kinase 4 |
| XM_341399 | Mapk8 | 0.0017 | 1.51 | similar to Mitogen-activated protein kinase 8 |
| XM_235169 | Mdm2 | 0.0165 | 1.34 | similar to transformed mouse 3T3 cell double minute 2 |
| NM_001009962 | Metrn | 0.0013 | 2.24 | meteorin, glial cell differentiation regulator |

| | | | | |
|---|---|---|---|---|
| NM_0010099 62 | Metrn | 0.0002 | 2.32 | meteorin, glial cell differentiation regulator |
| XM_224916 | Mfhas1 | 0.0078 | 1.33 | similar to malignant fibrous histiocytoma amplified sequence 1 |
| NM_019239 | Mgat3 | 0.0499 | 1.33 | mannoside acetyl glucosaminyltransferase 3 |
| NM_019239 | Mgat3 | 0.0341 | 1.59 | mannoside acetyl glucosaminyltransferase 3 |
| XM_221136 | Mgat5b | 0.0097 | 1.45 | similar to beta1,6-N-acetylglucosaminyltransferase IX |
| NM_0010083 70 | MGC10583 0 | 0.0422 | 1.64 | similar to Ras-related protein Rab-1B |
| NM_0010095 37 | MGC72997 | 0.0005 | 1.41 | hypothetical protein LOC494339 |
| NM_0010076 42 | MGC93920 | 0.0081 | 1.54 | hypothetical protein LOC295663 |
| XM_222997 | Mixl1 | 0.0252 | 1.70 | similar to Mix1 homeobox-like 1 |
| XM_239639 | Mmp17 | 0.0051 | 1.49 | similar to Matrix metalloproteinase-17 precursor |
| XM_239639 | Mmp17 | 0.0020 | 1.51 | similar to Matrix metalloproteinase-17 precursor |
| XM_340912 | Mpp2 | 0.0035 | 1.59 | similar to MAGUK p55 subfamily member 2 |
| XM_343301 | Mpped1 | 0.0000 | 2.38 | similar to CG16717-PA |
| NM_148890 | Msi1h | 0.0293 | 1.53 | Musashi homolog 1 |
| XM_213426 | Mtmr4 | 0.0014 | 1.56 | similar to myotubularin related protein 4 |
| XM_224271 | Mtmr6 | 0.0069 | 1.36 | similar to myotubularin related protein 6 |
| XM_233490 | Mycl1 | 0.0304 | 1.75 | similar to L-myc proto-oncogene protein |
| NM_013194 | Myh9 | 0.0165 | 1.43 | myosin, heavy polypeptide 9 |
| NM_182844 | Myrip | 0.0203 | 1.46 | myosin VIIA and Rab interacting protein |
| NM_022856 | Nab1 | 0.0279 | 1.55 | Ngfi-A binding protein 1 |
| XM_215931 | Ncoa5 | 0.0247 | 1.45 | similar to nuclear receptor coactivator 5 |
| XM_341248 | Nf2 | 0.0027 | 1.44 | similar to neurofibromatosis 2 |
| XM_343954 | Nog | 0.0004 | 1.41 | similar to Noggin precursor |
| NM_080577 | Nploc4 | 0.0053 | 1.94 | nuclear protein localization 4 |
| NM_017323 | Nr2c2 | 0.0386 | 1.40 | nuclear receptor subfamily 2, group C, member 2 |
| NM_031130 | Nr2f1 | 0.0058 | 1.50 | nuclear receptor subfamily 2, group F, member 1 |
| NM_145098 | Nrp1 | 0.0314 | 1.34 | neuropilin 1 |
| NM_021748 | Nsf | 0.0064 | 1.57 | N-ethylmaleimide sensitive fusion protein |
| NM_053463 | Nucb1 | 0.0357 | 1.33 | nucleobindin 1 |
| NM_0010242 43 | Nudt3 | 0.0000 | 1.75 | nudix (nucleotide diphosphate linked moiety X)-type motif 3 |
| NM_139091 | Nupl1 | 0.0310 | 1.38 | nucleoporin like 1 |
| NM_0010002 63 | Olr384 | 0.0432 | 1.67 | olfactory receptor Olr384 |
| XM_223556 | Osbp2 | 0.0058 | 1.34 | similar to oxysterol binding protein 2 |
| NM_017294 | Pacsin1 | 0.0092 | 1.42 | protein kinase C and casein kinase substrate in neurons 1 |
| NM_053306 | Pak2 | 0.0468 | 1.77 | p21-activated kinase 2 |
| XM_226334 | Papd5 | 0.0077 | 1.69 | similar to PAP associated domain-containing 5 |
| NM_053933 | Pcdha4 | 0.0104 | 1.41 | protocadherin alpha 4 |
| NM_022382 | Pde4dip | 0.0047 | 1.45 | myomegalin |
| NM_031591 | Pecam | 0.0466 | 1.42 | platelet/endothelial cell adhesion molecule |

| | | | | |
|---|---|---|---|---|
| NM_080477 | Pfkfb2 | 0.0002 | 1.84 | 6-phosphofructo-2-kinase/fructose-2, 6-bisphosphatase 2 isoform c |
| NM_057135 | Pfkfb3 | 0.0453 | 1.67 | 6-phosphofructo-2-kinase/fructose-2, 6-biphosphatase 3 |
| NM_019333 | Pfkfb4 | 0.0276 | 1.98 | 6-phosphofructo-2-kinase/fructose-2, 6-biphosphatase 4 |
| NM_001008374 | Pgrmc2 | 0.0154 | 1.39 | progesterone receptor membrane component 2 |
| XM_233700 | Phf13 | 0.0067 | 1.68 | similar to PHD finger protein 13 |
| NM_001017376 | Phyhip | 0.0000 | 1.47 | phytanoyl-CoA hydroxylase interacting protein |
| NM_053926 | Pip5k2a | 0.0279 | 1.67 | phosphatidylinositol-4-phosphate 5-kinase, type II, alpha |
| NM_053550 | Pip5k2b | 0.0057 | 1.47 | phosphatidylinositol-4-phosphate 5-kinase, type II, beta |
| XM_220629 | Pitpnm3 | 0.0278 | 1.32 | similar to Pitpnm family member 3 |
| XM_217372 | Plekhb2 | 0.0011 | 2.17 | similar to pleckstrin homology domain containing, family B member 2 |
| XM_236640 | Plxnb1 | 0.0191 | 1.35 | similar to plexin B1 |
| XM_235521 | Poldip3 | 0.0002 | 1.71 | similar to DNA polymerase delta interacting protein 3 |
| NM_017252 | Pou3f4 | 0.0326 | 1.81 | POU domain, class 3, transcription factor 4 |
| XM_341856 | Ppfia3 | 0.0347 | 1.32 | similar to Liprin-alpha-3 |
| NM_175755 | Ppm1f | 0.0400 | 1.40 | protein phosphatase 1F |
| NM_001013173 | Ppm1h | 0.0010 | 1.88 | protein phosphatase 1H |
| XM_222656 | Ppp1r12b | 0.0095 | 1.45 | similar to protein phosphatase 1, regulatory subunit 12B isoform b |
| NM_057116 | Ppp2r2c | 0.0001 | 2.44 | protein phosphatase 2, regulatory subunit B (PR 52), gamma isoform |
| XM_216739 | Ppp2r5e | 0.0263 | 1.45 | similar to epsilon isoform of regulatory subunit B56, protein phosphatase 2A |
| NM_017309 | Ppp3r1 | 0.0079 | 1.55 | protein phospatase 3, regulatory subunit B, alpha isoform,type 1 |
| XM_341661 | Prkaca | 0.0019 | 1.56 | similar to cAMP-dependent protein kinase, alpha-catalytic subunit |
| NM_019264 | Prkar2a | 0.0482 | 1.71 | protein kinase, cAMP-dependent, regulatory, type 2, alpha |
| NM_172022 | Prosapip1 | 0.0007 | 1.58 | ProSAPiP1 protein |
| XM_213385 | Prpf8 | 0.0003 | 2.37 | similar to Pre-mRNA-processing-splicing factor 8 |
| XM_213385 | Prpf8 | 0.0000 | 2.44 | similar to Pre-mRNA-processing-splicing factor 8 |
| NM_198738 | Psat1 | 0.0271 | 1.36 | phosphoserine aminotransferase 1 |
| XM_001066749 | Psd | 0.0097 | 1.33 | similar to pleckstrin and Sec7 domain containing homolog isoform 2 |
| XM_220754 | Psmd11 | 0.0111 | 1.34 | similar to 26S proteasome non-ATPase regulatory subunit 11 |
| NM_013081 | Ptk2 | 0.0154 | 1.46 | PTK2 protein tyrosine kinase 2 |
| NM_001008521 | Ptk9 | 0.0122 | 1.47 | protein tyrosine kinase 9 |
| NM_021740 | Ptma | 0.0094 | 1.36 | prothymosin alpha |

| | | | | |
|---|---|---|---|---|
| XM_341109 | Ptpn4 | 0.0449 | 1.33 | similar to protein tyrosine phosphatase, non-receptor type 4 |
| NM_017336 | Ptpro | 0.0099 | 1.41 | receptor-type protein tyrosine phosphatase D30 |
| NM_145094 | Rab31 | 0.0012 | 1.78 | RAB31, member RAS oncogene family |
| XM_213824 | Rab5b | 0.0003 | 1.51 | RAB5B, member RAS oncogene family |
| NM_019211 | Rasgrp1 | 0.0064 | 1.42 | RAS guanyl releasing protein 1 |
| XM_342003 | Rasgrp2 | 0.0014 | 1.35 | RAS guanyl releasing protein 2 (calcium and DAG-regulated) |
| XM_220782 | Rasl10b | 0.0090 | 1.41 | similar to RAS-like, family 10, member B |
| NM_031094 | Rbl2 | 0.0243 | 1.54 | retinoblastoma-like 2 (p130) |
| XM_231176 | Rbm18 | 0.0009 | 1.40 | similar to RNA binding motif protein 18 |
| XM_231176 | Rbm18 | 0.0047 | 1.48 | similar to RNA binding motif protein 18 |
| XM_214035 | Recc1 | 0.0106 | 1.55 | similar to Activator 1 140 kDa subunit |
| NM_144741 | Retn | 0.0365 | 1.84 | resistin |
| XM_235656 | RGD1305671 | 0.0058 | 1.70 | similar to DISCO Interacting Protein 2 CG7020-PA |
| XM_001078354 | RGD1306259 | 0.0101 | 1.52 | similar to neuron navigator 3 |
| XM_343289 | RGD1306591 | 0.0143 | 1.60 | similar to Putative MAP kinase-activating protein C22orf5 |
| XM_235527 | RGD1306698 | 0.0296 | 1.91 | similar to Malonyl CoA-acyl carrier protein transacylase |
| XM_226983 | RGD1307225 | 0.0051 | 1.42 | similar to EGF-like-domain, multiple 3 |
| XM_345231 | RGD1307244 | 0.0070 | 1.40 | similar to CG5805-PA |
| NM_001025012 | RGD1307449 | 0.0338 | 1.40 | hypothetical protein LOC316687 |
| NM_001014226 | RGD1308915 | 0.0120 | 1.36 | hypothetical protein LOC363545 |
| XM_220262 | RGD1308952 | 0.0005 | 1.61 | similar to Rab11 family-interacting protein 3 |
| XM_226732 | RGD1310139 | 0.0092 | 1.69 | similar to microtubule associated serine/threonine kinase 2 |
| XM_230641 | RGD1311344 | 0.0371 | 1.54 | double zinc ribbon and ankyrin repeat domains 1 |
| XM_575355 | RGD1561053 | 0.0372 | 1.66 | similar to claudin 12 |
| XM_576077 | RGD1561926 | 0.0296 | 1.40 | similar to Nuclear protein SkiP |
| XM_577254 | RGD1564490 | 0.0062 | 2.22 | similar to coronin, actin binding protein 1C |
| XM_576400 | RGD1564560 | 0.0002 | 2.40 | similar to Probable ATP-dependent RNA helicase DDX6 |
| XM_217645 | RGD1565768 | 0.0002 | 1.98 | actin-binding LIM protein 1 |
| NM_170666 | Rims4 | 0.0144 | 1.50 | regulating synaptic membrane exocytosis 4 |
| XM_342522 | Rnf24 | 0.0444 | 1.57 | similar to ring finger protein 24 |
| NM_133518 | Rph3a | 0.0002 | 1.70 | rabphilin 3A homolog |
| XM_216515 | Rragc | 0.0126 | 1.39 | Ras-related GTP binding C |
| NM_080402 | Ryk | 0.0120 | 1.37 | receptor-like tyrosine kinase |

| | | | | |
|---|---|---|---|---|
| XM_223945 | Samd4 | 0.0036 | 1.65 | similar to sterile alpha motif domain containing 4 isoform 1 |
| NM_054001 | Scarb2 | 0.0013 | 1.35 | CD36 antigen (collagen type I receptor, thrombospondin receptor)-like 2 |
| NM_012877 | Scn2b | 0.0007 | 2.09 | sodium channel, voltage-gated, type II, beta polypeptide |
| XM_345848 | Scrt1 | 0.0477 | 1.56 | similar to scratch homolog 1, zinc finger protein |
| NM_017310 | Sema3a | 0.0137 | 1.40 | sema domain, immunoglobulin domain (Ig), |
| XM_221566 | Senp7 | 0.0037 | 1.34 | similar to SUMO1/sentrin specific protease 7 isoform 2 |
| XM_220423 | Sep-08 | 0.0239 | 1.55 | similar to Septin-8 |
| NM_201350 | Shank2 | 0.0062 | 1.35 | SH3/ankyrin domain gene 2 isoform a |
| XM_001057072 | Skil | 0.0281 | 1.37 | similar to Ski-like protein isoform 1 |
| NM_134363 | Slc12a5 | 0.0042 | 1.40 | solute carrier family 12 member 5 |
| XM_001060536 | Slc12a7 | 0.0057 | 1.40 | similar to Solute carrier family 12 member 7 |
| NM_022270 | Slc22a4 | 0.0408 | 1.65 | solute carrier family 22 (organic cation transporter), member 4 |
| NM_145677 | Slc25a25 | 0.0007 | 1.81 | mitochondrial Ca2+-dependent solute carrier |
| XM_342286 | Slc39a1 | 0.0379 | 1.36 | similar to Zinc transporter ZIP1 |
| NM_053811 | Slc9a3r2 | 0.0020 | 1.58 | solute carrier family 9 isoform 3 regulator 2 |
| NM_053811 | Slc9a3r2 | 0.0022 | 1.62 | solute carrier family 9 isoform 3 regulator 2 |
| XM_230716 | Snph | 0.0001 | 1.88 | hypothetical protein |
| NM_001013085 | Snx10 | 0.0049 | 1.76 | sorting nexin 10 |
| XM_232945 | Snx30 | 0.0082 | 1.40 | similar to sorting nexin family member 30 |
| XM_221947 | Snx8 | 0.0053 | 1.81 | similar to sorting nexin 8 |
| XM_340879 | Spag9 | 0.0003 | 1.56 | similar to sperm associated antigen 9 isoform 1 |
| XM_213437 | Spop | 0.0028 | 1.53 | similar to Speckle-type POZ protein |
| NM_138918 | Ss18l1 | 0.0044 | 1.75 | synovial sarcoma translocation gene on chromosome 18-like 1 |
| NM_013036 | Sstr4 | 0.0193 | 1.38 | somatostatin receptor 4 |
| NM_001013219 | St3gal1 | 0.0011 | 2.42 | ST3 beta-galactoside alpha-2,3-sialyltransferase 1 |
| NM_203337 | St3gal4 | 0.0068 | 1.45 | alpha-2,3-sialyltransferase ST3Gal IV |
| XM_344970 | Stk32c | 0.0000 | 2.59 | similar to serine/threonine kinase 32C |
| NM_019148 | Strn | 0.0371 | 1.45 | striatin, calmodulin binding protein |
| NM_021659 | Syt7 | 0.0002 | 1.65 | synaptotagmin 7 |
| NM_021659 | Syt7 | 0.0001 | 1.95 | synaptotagmin 7 |
| NM_138840 | Tgoln2 | 0.0021 | 1.40 | trans-golgi network protein 1 |
| NM_012887 | Tmpo | 0.0160 | 1.42 | thymopoietin |
| XM_222478 | Tnpo2 | 0.0016 | 1.37 | similar to transportin 2 (importin 3, karyopherin beta 2b) |
| XM_341961 | Tollip | 0.0002 | 2.22 | similar to toll interacting protein |
| XM_340810 | Tom1l2 | 0.0005 | 1.47 | similar to target of myb1-like 2 isoform 2 |
| NM_212519 | Tomm70a | 0.0054 | 1.40 | translocase of outer mitochondrial membrane 70 homolog A |
| XM_340806 | Trim11 | 0.0331 | 1.34 | similar to tripartite motif protein 11 |

| | | | | |
|---|---|---|---|---|
| XM_342183 | Trim23 | 0.0000 | 1.49 | similar to GTP-binding protein ARD-1 |
| NM_0010040 90 | Tspan5 | 0.0003 | 1.68 | transmembrane 4 superfamily member 9 |
| XM_221962 | Ttyh3 | 0.0033 | 1.44 | similar to tweety 3 |
| NM_013077 | Tub | 0.0020 | 1.41 | tubby |
| NM_031138 | Ube2b | 0.0487 | 2.18 | ubiquitin conjugating enzyme |
| XM_221132 | Ube2o | 0.0002 | 1.46 | similar to ubiquitin-conjugating enzyme E2O |
| XM_215948 | Ube2v1 | 0.0025 | 1.39 | similar to ubiquitin-conjugating enzyme E2 variant 1 |
| XM_341867 | Ube3a | 0.0161 | 1.33 | similar to ubiquitin protein ligase E3A isoform 2 |
| NM_031795 | Ugcg | 0.0343 | 1.35 | UDP-glucose ceramide glucosyltransferase |
| NM_022206 | Unc5a | 0.0048 | 1.35 | unc-5 homolog A |
| XM_220534 | Usp22 | 0.0053 | 1.51 | hypothetical protein |
| XM_220798 | Usp32 | 0.0030 | 1.46 | similar to ubiquitin specific protease 32 |
| XM_0010686 66 | Usp33 | 0.0359 | 1.42 | similar to ubiquitin specific protease 33 |
| NM_0010247 90 | Usp7 | 0.0004 | 1.36 | ubiquitin specific protease 7 (herpes virus-associated) |
| NM_013090 | Vamp1 | 0.0058 | 1.46 | vesicle-associated membrane protein 1 |
| NM_013090 | Vamp1 | 0.0020 | 1.68 | vesicle-associated membrane protein 1 |
| NM_031353 | Vdac1 | 0.0000 | 2.27 | voltage-dependent anion channel 1 |
| XM_225314 | Vmp | 0.0204 | 1.50 | similar to vesicular membrane protein p24 |
| NM_0010131 67 | Wasf2 | 0.0168 | 1.36 | WAS protein family, member 2 |
| NM_0010141 35 | Wdr1 | 0.0115 | 1.45 | WD repeat domain 1 |
| XM_232730 | Wdtc1 | 0.0063 | 1.60 | similar to WD and tetratricopeptide repeats 1 |
| XM_220714 | XM_22071 4 | 0.0129 | 1.37 | similar to spinster-like protein |
| XM_223739 | XM_22373 9 | 0.0020 | 1.33 | predicted diacylglycerol kinase, theta |
| XM_226100 | XM_22610 0 | 0.0022 | 1.84 | zinc finger protein 239 |
| XM_237097 | XM_23709 7 | 0.0113 | 1.59 | similar to RIKEN cDNA 2310066K23 |
| XM_243623 | XM_24362 3 | 0.0349 | 1.43 | predicted mitogen-activated protein kinase kinase kinase 7 interacting protein 1 |
| XM_341137 | XM_34113 7 | 0.0003 | 1.48 | similar to constitutive photomorphogenic protein 1 |
| XM_342562 | XM_34256 2 | 0.0398 | 1.37 | similar to RIKEN cDNA B230339M05 |
| XM_343028 | XM_34302 8 | 0.0449 | 1.67 | transmembrane protein 214 |
| XM_343348 | XM_34334 8 | 0.0351 | 1.43 | LOC363015 |
| XM_343871 | XM_34387 1 | 0.0002 | 1.89 | similar to p19 |
| XM_344884 | XM_34488 4 | 0.0233 | 1.59 | similar to hypothetical protein MGC25497 |
| XM_345618 | XM_34561 8 | 0.0282 | 1.37 | similar to agrin precursor |

| | | | | |
|---|---|---|---|---|
| XM_232194 | Xpc | 0.0006 | 1.59 | similar to DNA-repair protein complementing XP-C cells homolog |
| XM_223117 | Xpr1 | 0.0228 | 1.54 | similar to xenotropic and polytropic retrovirus receptor 1 |
| XM_341912 | Xylt1 | 0.0343 | 2.01 | similar to Xylosyltransferase 1 |
| NM_0010131 81 | Zbtb16 | 0.0000 | 1.61 | zinc finger and BTB domain containing 16 |
| NM_138613 | Zfp179 | 0.0479 | 1.52 | zinc finger protein 179 |
| NM_022678 | Zfp238 | 0.0007 | 1.34 | zinc finger protein 238 |
| XM_226153 | Zfp521 | 0.0420 | 2.42 | similar to zinc finger protein 521 isoform 1 |
| XM_230026 | Zfp533 | 0.0043 | 1.56 | similar to zinc finger protein 533 |
| XM_219325 | Zfpn1a5 | 0.0155 | 2.34 | similar to zinc finger protein, subfamily 1A, 5 |
| XM_233341 | Zfyve9 | 0.0015 | 1.84 | similar to Zinc finger FYVE domain-containing protein 9 |
| NM_203367 | Zmynd11 | 0.0265 | 1.33 | BS69 protein beta isoform |
| XM_344076 | Znf498 | 0.0162 | 1.44 | similar to zinc finger protein 498 |

**Supplementary Table 7: List of genes differentially downregulated in the mPFC transcriptome of adult JFlx animals.**

Six month old JFlx animals and their age-matched controls were subjected to microarray analysis. The list of all the significantly downregulated genes is shown in the table ($p < 0.05$ ; Fold change < 0.74).

| Systematic Name | GeneName | *P* Value | Fold Change | Product |
|---|---|---|---|---|
| XM_225368 | Abt1 | 0.0048 | 0.48 | similar to activator of basal transcription |
| XM_235977 | Adamts8 | 0.0087 | 0.47 | similar to ADAMTS-8 precursor |
| NM_022190 | Agc1 | 0.0034 | 0.74 | aggrecan 1 |
| NM_001008342 | Akr1e1 | 0.0060 | 0.75 | aldo-keto reductase family 1, member E1 |
| NM_012496 | Aldob | 0.0422 | 0.71 | aldolase B |
| NM_012822 | Alox5 | 0.0166 | 0.70 | arachidonate 5-lipoxygenase |
| NM_012779 | Aqp5 | 0.0168 | 0.73 | aquaporin 5 |
| XM_221775 | Arhgef18 | 0.0400 | 0.71 | similar to rho/rac guanine nucleotide exchange factor 18 |
| NM_012912 | Atf3 | 0.0437 | 0.58 | activating transcription factor 3 |
| NM_001011972 | Atp6v0d2 | 0.0107 | 0.51 | ATPase, H+ transporting, V0 subunit D, isoform 2 |
| NM_212490 | Atp6v1g2 | 0.0068 | 0.71 | ATPase, H+ transporting, V1 subunit G isoform 2 |
| NM_053019 | Avpr1a | 0.0363 | 0.64 | arginine vasopressin receptor 1A |
| NM_012826 | Azgp1 | 0.0462 | 0.39 | alpha-2-glycoprotein 1, zinc |
| XM_345362 | B3galt1 | 0.0205 | 0.65 | similar to Beta-1,3-galactosyltransferase 1 |
| NM_016993 | Bcl2 | 0.0356 | 0.76 | B-cell leukemia/lymphoma 2 |
| NM_001007666 | Bcs1l | 0.0373 | 0.64 | BCS1-like |
| XM_342529 | Bfsp1 | 0.0178 | 0.67 | similar to beaded filament structural protein in lens-CP94 |
| NM_012863 | Bhlhb8 | 0.0007 | 0.45 | muscle, intestine and stomach expression 1 |
| XM_218837 | Blm | 0.0030 | 0.70 | similar to Bloom syndrome protein homolog |
| BM986492 | BM986492 | 0.0284 | 0.48 | EST594086 Rat gene index |
| XM_235552 | Brd1 | 0.0348 | 0.63 | similar to bromodomain containing protein 1 |
| NM_001013176 | Btg4 | 0.0044 | 0.45 | B-cell translocation gene 4 |
| NM_001013176 | Btg4 | 0.0342 | 0.51 | B-cell translocation gene 4 |
| XM_574426 | Ccne1 | 0.0312 | 0.74 | similar to G1/S-specific cyclin-E1 |
| NM_021866 | Ccr2 | 0.0255 | 0.60 | chemokine (C-C motif) receptor 2 |
| NM_134327 | Cd69 | 0.0022 | 0.56 | CD69 antigen |
| NM_133571 | Cdc25a | 0.0386 | 0.58 | cell division cycle 25 homolog A |
| XM_221438 | Cdgap | 0.0167 | 0.48 | similar to Cdc42 GTPase-activating protein |
| NM_053644 | Cdh23 | 0.0247 | 0.72 | cadherin 23 |
| NM_012702 | Ceacam3 | 0.0175 | 0.70 | carcinoembryonic antigen-related cell adhesion molecule 3 |
| XM_342521 | Cenpb | 0.0051 | 0.74 | similar to Major centromere autoantigen B |
| NM_001004098 | Cenpc1 | 0.0020 | 0.68 | centromere protein C 1 |
| NM_139087 | Cgref1 | 0.0053 | 0.72 | cell growth regulator with EF hand domain 1 |
| NM_001009640 | Cirh1a | 0.0263 | 0.66 | cirrhosis, autosomal recessive 1A (cirhin) |
| NM_001009640 | Cirh1a | 0.0279 | 0.73 | cirrhosis, autosomal recessive 1A (cirhin) |

| | | | | |
|---|---|---|---|---|
| NM_031699 | Cldn1 | 0.0131 | 0.71 | claudin 1 |
| NM_001003707 | Clec4d | 0.0319 | 0.51 | C-type lectin, superfamily member 8 |
| NM_134377 | Clstn2 | 0.0090 | 0.71 | calsyntenin 2 |
| NM_013225 | Cntn6 | 0.0128 | 0.73 | contactin 6 |
| XM_222002 | Cops6 | 0.0056 | 0.69 | similar to COP9 signalosome subunit 6 |
| NM_001029909 | CPG2 | 0.0145 | 0.39 | CPG2 protein |
| XM_235061 | Cradd | 0.0142 | 0.70 | similar to Death domain-containing protein CRADD |
| XM_001065365 | Ctxn | 0.0444 | 0.69 | cortexin |
| XM_215274 | Cuedc2 | 0.0309 | 0.76 | CUE domain containing 2 |
| XM_341542 | Cul2 | 0.0025 | 0.70 | similar to cullin 2 |
| NM_153721 | Cxcl7 | 0.0037 | 0.64 | pro-platelet basic protein |
| NM_012538 | Cyp11b2 | 0.0202 | 0.74 | cytochrome P450, family 11, subfamily B, polypeptide 2 |
| NM_024130 | Dctn1 | 0.0154 | 0.74 | dynactin 1 |
| NM_001004239 | Dctn2 | 0.0075 | 0.74 | dynactin 2 |
| NM_032063 | Dll1 | 0.0450 | 0.47 | delta-like 1 |
| NM_001024879 | Dnpep | 0.0465 | 0.66 | aspartyl aminopeptidase |
| XM_345909 | Dock6 | 0.0143 | 0.74 | similar to Dedicator of cytokinesis protein 6 |
| XM_224344 | Dok2 | 0.0388 | 0.52 | similar to Docking protein 2 |
| NM_012549 | Edn2 | 0.0204 | 0.71 | endothelin 2 |
| NM_053950 | Eif2b4 | 0.0179 | 0.59 | eukaryotic translation initiation factor 2B, subunit 4 delta |
| NM_053950 | Eif2b4 | 0.0099 | 0.66 | eukaryotic translation initiation factor 2B, subunit 4 delta |
| NM_001008324 | Eif4b | 0.0018 | 0.56 | eukaryotic translation initiation factor 4B |
| NM_172326 | Elac2 | 0.0012 | 0.72 | elaC homolog 2 |
| NM_178106 | Entpd3 | 0.0113 | 0.71 | ectonucleoside triphosphate diphosphohydrolase 3 |
| XM_231658 | Epha1 | 0.0097 | 0.55 | similar to Eph receptor A1 |
| XM_226987 | Evi1 | 0.0106 | 0.58 | similar to Ecotropic virus integration 1 site protein |
| NM_144756 | Faim2 | 0.0228 | 0.70 | Fas apoptotic inhibitory molecule 2 |
| NM_001024237 | Farsla | 0.0025 | 0.71 | phenylalanine-tRNA synthetase-like, alpha subunit |
| NM_022211 | Fgf5 | 0.0200 | 0.45 | fibroblast growth factor 5 |
| XM_342763 | Fkbp4 | 0.0000 | 0.75 | similar to FK506-binding protein 4 |
| XM_222082 | Fkbp6 | 0.0431 | 0.66 | similar to FK506-binding protein 6 |
| NM_001008279 | Fliih | 0.0218 | 0.68 | flightless I homolog |
| NM_019172 | Galr2 | 0.0003 | 0.45 | galanin receptor 2 |
| NM_019172 | Galr2 | 0.0058 | 0.51 | galanin receptor 2 |
| NM_019172 | Galr2 | 0.0032 | 0.52 | galanin receptor 2 |
| NM_019172 | Galr2 | 0.0009 | 0.58 | galanin receptor 2 |
| NM_019172 | Galr2 | 0.0016 | 0.61 | galanin receptor 2 |
| XM_227762 | Gbp4 | 0.0050 | 0.54 | similar to guanylate nucleotide binding protein 4 |
| NM_012565 | Gck | 0.0341 | 0.74 | glucokinase |
| NM_173312 | Gcnt3 | 0.0149 | 0.49 | glucosaminyl (N-acetyl) transferase 3, mucin type |
| NM_019151 | Gdf8 | 0.0499 | 0.75 | growth differentiation factor 8 |

| | | | | |
|---|---|---|---|---|
| NM_017094 | Ghr | 0.0206 | 0.69 | growth hormone receptor |
| NM_145680 | Gimap5 | 0.0308 | 0.73 | GTPase, IMAP family member 5 isoform 2 |
| XM_219785 | Gldc | 0.0076 | 0.54 | similar to glycine decarboxylase |
| BC072694 | Glul | 0.0014 | 0.70 | Glul protein |
| NM_022255 | Gpr173 | 0.0064 | 0.64 | G-protein coupled receptor 173 |
| NM_080579 | Gpr19 | 0.0101 | 0.74 | G protein-coupled receptor 19 |
| XM_231251 | Gpr21 | 0.0033 | 0.57 | similar to RAB GTPase activating protein 1 isoform a |
| NM_138549 | Gpsn2 | 0.0233 | 0.67 | glycoprotein, synaptic 2 |
| NM_017113 | Grn | 0.0001 | 0.73 | granulin |
| NM_001012067 | Grwd1 | 0.0357 | 0.75 | glutamate-rich WD repeat containing 1 |
| NM_053345 | Gtf2a2 | 0.0306 | 0.67 | general transcription factor Iia 2 |
| XM_343535 | Gtpbp2 | 0.0154 | 0.71 | similar to GTP binding protein 2 |
| NM_017015 | Gusb | 0.0227 | 0.47 | glucuronidase, beta |
| NM_130826 | Hadha | 0.0021 | 0.67 | mitochondrial trifunctional protein, alpha subunit |
| NM_033349 | Hagh | 0.0004 | 0.68 | hydroxyacyl glutathione hydrolase |
| NM_033349 | Hagh | 0.0035 | 0.74 | hydroxyacyl glutathione hydrolase |
| NM_001011934 | Hbs1l | 0.0111 | 0.75 | Hbs1-like |
| NM_013064 | Hcrtr1 | 0.0435 | 0.72 | hypocretin receptor 1 |
| NM_022605 | Hpse | 0.0351 | 0.69 | heparanase |
| NM_053318 | Hpx | 0.0480 | 0.72 | hemopexin |
| XM_232247 | Il17r | 0.0232 | 0.64 | similar to Interleukin-17 receptor precursor |
| NM_053374 | Il18bp | 0.0270 | 0.70 | interferon gamma inducing factor binding protein |
| NM_017183 | Il8rb | 0.0483 | 0.69 | interleukin 8 receptor, beta |
| XM_215184 | Incenp | 0.0004 | 0.70 | similar to inner centromere protein |
| NM_001012007 | Irgm | 0.0248 | 0.69 | immunity-related GTPase family, M |
| NM_031046 | Itpr2 | 0.0022 | 0.50 | inositol 1,4,5-triphosphate receptor 2 |
| NM_053486 | Kif3c | 0.0167 | 0.72 | kinesin family member 3C |
| NM_053486 | Kif3c | 0.0127 | 0.73 | kinesin family member 3C |
| NM_181479 | Kir3dl1 | 0.0252 | 0.72 | killer cell immunoglobulin-like receptor, three domains, long cytoplasmic tail, 1 |
| XM_223423 | Klf3_mapped | 0.0327 | 0.55 | similar to Kruppel-like factor 3 |
| XM_218649 | Klk7 | 0.0399 | 0.59 | similar to kallikrein 7 |
| XM_234264 | L2hgdh | 0.0020 | 0.49 | similar to Y45G12B.3 |
| XM_237536 | Lama1 | 0.0217 | 0.65 | similar to Laminin alpha-1 chain precursor |
| XM_219866 | Lama2 | 0.0266 | 0.64 | similar to Laminin alpha-2 chain precursor |
| NM_199384 | Laptm4a | 0.0111 | 0.72 | lysosomal-associated protein transmembrane 4 alpha |
| NM_001009637 | Lars | 0.0468 | 0.74 | leucyl-tRNA synthetase |
| NM_001014094 | LOC315883 | 0.0144 | 0.69 | hypothetical protein LOC315883 |
| NM_001024351 | LOC500761 | 0.0305 | 0.60 | hypothetical protein LOC500761 |
| XM_001057185 | LOC680266 | 0.0035 | 0.53 | similar to Guanine nucleotide-binding protein beta subunit-like protein 1 |

| | | | | |
|---|---|---|---|---|
| XM_001068107 | LOC68393 2 | 0.0495 | 0.60 | similar to PWWP domain containing 2 |
| XM_232107 | Loxl3 | 0.0010 | 0.49 | similar to lysyl oxidase-like 3 |
| XM_341047 | Lrch4 | 0.0215 | 0.66 | similar to leucine rich repeat protein 4, neuronal |
| NM_031050 | Lum | 0.0193 | 0.59 | lumican |
| NM_001009620 | MGC10560 1 | 0.0071 | 0.60 | claudin-like protein 24 |
| NM_001009620 | MGC10560 1 | 0.0024 | 0.63 | claudin-like protein 24 |
| XM_220785 | Mmp28 | 0.0044 | 0.73 | matrix metallopeptidase 28 |
| XM_216323 | Mrpl15 | 0.0392 | 0.67 | similar to mitochondrial ribosomal protein L15 |
| NM_133539 | Mrpl17 | 0.0068 | 0.73 | mitochondrial ribosomal protein L17 |
| XM_213513 | Mrps7 | 0.0287 | 0.75 | similar to mitchondrial ribosomal protein S7 |
| NM_030863 | Msn | 0.0443 | 0.54 | moesin |
| XM_227485 | Msr2 | 0.0470 | 0.76 | similar to macrophage scavenger receptor 2 |
| NM_001012038 | Mtmr3 | 0.0051 | 0.74 | myotubularin related protein 3 |
| NM_133546 | Myd116 | 0.0052 | 0.73 | myeloid differentiation primary response gene 116 |
| NM_019325 | Myh4 | 0.0161 | 0.63 | myosin, heavy polypeptide 4, skeletal muscle |
| NM_012987 | Nes | 0.0376 | 0.71 | nestin |
| NM_019326 | Neurod2 | 0.0202 | 0.72 | neurogenic differentiation 2 |
| NM_012799 | Nmbr | 0.0368 | 0.57 | neuromedin B receptor |
| NM_022239 | Nmu | 0.0121 | 0.56 | neuromedin |
| NM_030841 | Nptxr | 0.0408 | 0.73 | neuronal pentraxin receptor |
| NM_031981 | Nsfl1c | 0.0017 | 0.76 | NSFL1 (p97) cofactor (p47) |
| NM_053830 | Nup107 | 0.0130 | 0.56 | nucleoporin 107 |
| NM_001001076 | Olr1064 | 0.0277 | 0.73 | olfactory receptor Olr1064 |
| NM_001000867 | Olr1166 | 0.0063 | 0.58 | olfactory receptor Olr1166 |
| NM_001000167 | Olr153 | 0.0437 | 0.42 | olfactory receptor Olr153 |
| NM_001000532 | Olr1568 | 0.0138 | 0.48 | olfactory receptor Olr1568 |
| XM_344047 | Olr1571 | 0.0215 | 0.62 | similar to olfactory receptor 19 |
| NM_001000689 | Olr34 | 0.0030 | 0.52 | olfactory receptor Olr34 |
| NM_001000390 | Olr422 | 0.0030 | 0.54 | olfactory receptor Olr422 |
| NM_001000311 | Olr502 | 0.0134 | 0.70 | olfactory receptor Olr502 |
| NM_001000922 | Olr735 | 0.0164 | 0.47 | olfactory receptor Olr735 |
| XM_222641 | Optc | 0.0305 | 0.75 | similar to opticin |
| NM_053656 | P2rx2 | 0.0080 | 0.67 | purinergic receptor P2X2 |
| NM_053289 | Pap | 0.0409 | 0.48 | pancreatitis-associated protein |
| XM_213447 | Pcgf2 | 0.0266 | 0.56 | similar to Polycomb group RING finger protein 2 |
| XM_225895 | Pde6a | 0.0474 | 0.60 | similar to phosphodiesterase 6A, cGMP-specific, rod, alpha |
| NM_023978 | Per3 | 0.0143 | 0.71 | period homolog 3 |
| NM_145676 | Pgea1 | 0.0176 | 0.76 | PKD2 interactor, golgi and endoplasmic reticulum associated 1 |
| NM_019334 | Pitx2 | 0.0043 | 0.49 | paired-like homeodomain transcription factor 2 isoform 2 |
| XM_342340 | Pla2g12a | 0.0002 | 0.63 | similar to group XII-1 phospholipase A2 isoform 1 |

| | | | | |
|---|---|---|---|---|
| NM_080688 | Plcd4 | 0.0472 | 0.69 | phospholipase C, delta 4 |
| NM_001025750 | Plek | 0.0159 | 0.73 | pleckstrin |
| NM_013094 | Plin | 0.0316 | 0.68 | perilipin |
| NM_053827 | Plod1 | 0.0032 | 0.30 | procollagen-lysine, 2-oxoglutarate 5-dioxygenase 1 |
| XM_223080 | Plxna2 | 0.0017 | 0.73 | similar to plexin A2 |
| NM_134393 | Pmfbp1 | 0.0150 | 0.51 | polyamine modulated factor 1 binding protein 1 |
| NM_032077 | Pon1 | 0.0254 | 0.65 | paraoxonase 1 |
| NM_013124 | Pparg | 0.0250 | 0.55 | peroxisome proliferator activated receptor, gamma |
| NM_134359 | Ppp4c | 0.0006 | 0.73 | protein phosphatase 4, catalytic subunit |
| XM_230542 | Prnd | 0.0080 | 0.65 | similar to prion protein dublet |
| NM_012803 | Proc | 0.0286 | 0.68 | protein C |
| NM_023976 | Prx | 0.0106 | 0.46 | periaxin |
| XM_226439 | Psmd7 | 0.0132 | 0.72 | similar to 26S proteasome non-ATPase regulatory subunit 7 |
| NM_001012143 | Ptdsr | 0.0175 | 0.75 | jumonji domain containing 6 |
| NM_013100 | Ptger1 | 0.0354 | 0.57 | prostaglandin E receptor 1 |
| NM_031088 | Ptger2 | 0.0258 | 0.71 | prostaglandin E receptor 2, subtype EP2 |
| XM_220982 | Ptrf | 0.0105 | 0.24 | similar to Polymerase I and transcript release factor |
| NM_017234 | Pxmp3 | 0.0000 | 0.56 | peroxin 2 |
| NM_145774 | Rab38 | 0.0004 | 0.52 | Rab38, member of RAS oncogene family |
| XM_236242 | Rab39 | 0.0494 | 0.72 | similar to RAB39, member RAS oncogene family |
| NM_017313 | Rab3ip | 0.0007 | 0.74 | RAB3A interacting protein |
| NM_031774 | Rabac1 | 0.0434 | 0.73 | rab acceptor 1 (prenylated) |
| NM_019124 | Rabep1 | 0.0094 | 0.73 | rabaptin 5 |
| NM_001013152 | Rcl1 | 0.0235 | 0.64 | RNA terminal phosphate cyclase-like 1 |
| NM_001011916 | Rec8L1 | 0.0032 | 0.59 | REC8 homolog |
| XM_344661 | Reep5 | 0.0337 | 0.75 | similar to receptor accessory protein 5 |
| XM_223688 | Rel | 0.0094 | 0.43 | similar to C-Rel proto-oncogene protein |
| XM_576183 | Rexo1 | 0.0490 | 0.72 | similar to transcription elongation factor B polypeptide 3 binding protein 1 isoform 1 |
| NM_001004266 | RGD1303144 | 0.0260 | 0.67 | HypB, AroK, ADK and rho factor domain containing protein RGD1303144 |
| XM_228076 | RGD1305633 | 0.0163 | 0.53 | similar to Protein KIAA0179 |
| NM_001029923 | RGD1306596 | 0.0282 | 0.67 | hypothetical protein LOC362595 |
| XM_220112 | RGD1306962 | 0.0468 | 0.74 | similar to F55A4.8a |
| XM_235513 | RGD1307309 | 0.0358 | 0.61 | similar to proliferation associated nuclear element 1 isoform 1 |
| NM_001037192 | RGD1307890 | 0.0472 | 0.67 | hypothetical protein LOC304851 |
| NM_001013974 | RGD1308813 | 0.0391 | 0.69 | hypothetical protein LOC303606 |
| XM_227686 | RGD1311822 | 0.0360 | 0.54 | similar to CG10428-PA |

| | | | | |
|---|---|---|---|---|
| XM_576402 | RGD1562173 | 0.0057 | 0.73 | similar to Zinc-finger protein ZPR1 |
| XM_216520 | Rhbdl2 | 0.0367 | 0.75 | similar to rhomboid-related protein 2 |
| NM_001013430 | Rhoh | 0.0209 | 0.69 | ras homolog gene family, member H |
| XM_221619 | Ripk4 | 0.0064 | 0.45 | similar to receptor-interacting serine-threonine kinase 4 |
| NM_022697 | Rpl28 | 0.0097 | 0.48 | ribosomal protein L28 |
| NM_012521 | S100g | 0.0099 | 0.55 | calbindin 3 |
| NM_023955 | Scamp2 | 0.0270 | 0.68 | secretory carrier membrane protein 2 |
| XM_228030 | Scube3 | 0.0205 | 0.61 | similar to signal peptide, CUB and EGF-like domain containing protein 3 |
| NM_001006978 | Sec13l1 | 0.0425 | 0.75 | SEC13-like 1 |
| NM_001031642 | Serpinb1a | 0.0475 | 0.64 | serine (or cysteine) proteinase inhibitor, clade B, member 1a |
| NM_001031642 | Serpinb1a | 0.0383 | 0.67 | serine (or cysteine) proteinase inhibitor, clade B, member 1a |
| NM_001011892 | Serpinf2 | 0.0253 | 0.63 | serine (or cysteine) peptidase inhibitor, clade F, member 2 |
| NM_139341 | Slc15a3 | 0.0158 | 0.72 | peptide/histidine transporter PHT2 |
| NM_053580 | Slc27a1 | 0.0397 | 0.74 | solute carrier family 27, member 1 |
| NM_052983 | Slc5a5 | 0.0216 | 0.61 | solute carrier family 5, member 5 |
| NM_012654 | Slc9a3 | 0.0105 | 0.74 | solute carrier family 9, member 3 |
| NM_053372 | Slpi | 0.0285 | 0.55 | secretory leukocyte peptidase inhibitor |
| XM_213972 | Smyd2 | 0.0077 | 0.68 | similar to SET and MYND domain-containing protein 2 |
| NM_080777 | Sncb | 0.0009 | 0.70 | synuclein, beta |
| NM_053411 | Snx1 | 0.0092 | 0.66 | sorting nexin 1 |
| NM_053411 | Snx1 | 0.0033 | 0.70 | sorting nexin 1 |
| NM_001024781 | Sox18 | 0.0105 | 0.76 | SRY-box containing gene 18 |
| XM_342826 | Spag8 | 0.0478 | 0.63 | similar to sperm associated antigen 8 |
| XM_342235 | Spata5 | 0.0215 | 0.58 | similar to spermatogenesis associated 5 |
| XM_342134 | Spock2 | 0.0002 | 0.57 | hypothetical protein |
| NM_175843 | Sqstm1 | 0.0163 | 0.70 | sequestosome 1 isoform 1 |
| NM_001011965 | Stom | 0.0135 | 0.39 | stomatin |
| NM_012664 | Syp | 0.0121 | 0.75 | synaptophysin |
| NM_012664 | Syp | 0.0018 | 0.75 | synaptophysin |
| NM_001009540 | Tacstd2 | 0.0373 | 0.57 | tumor-associated calcium signal transducer 2 |
| NM_001009344 | Tbrg1 | 0.0237 | 0.74 | transforming growth factor beta regulated gene 1 |
| NM_021673 | Tcam1 | 0.0249 | 0.56 | testicular cell adhesion molecule 1 |
| BC059146 | Tgfb1i4 | 0.0003 | 0.75 | Tgfb1i4 protein |
| XM_573983 | Tgfbi | 0.0488 | 0.47 | similar to transforming growth factor, beta induced |
| XM_214778 | Thbs2 | 0.0411 | 0.62 | similar to Thrombospondin-2 precursor |
| NM_001012155 | Tm9sf1 | 0.0198 | 0.69 | transmembrane 9 superfamily member 1 |
| NM_001001799 | Tmem35 | 0.0055 | 0.67 | spinal cord expression protein 4 |
| XM_213211 | Tnfrsf17 | 0.0122 | 0.60 | similar to Tumor necrosis factor receptor superfamily member 17 |

| | | | | |
|---|---|---|---|---|
| XM_213211 | Tnfrsf17 | 0.0295 | 0.63 | similar to Tumor necrosis factor receptor superfamily member 17 |
| XM_222022 | Trfr2 | 0.0016 | 0.53 | similar to Transferrin receptor protein 2 |
| NM_001012210 | Trim13 | 0.0141 | 0.56 | tripartite motif protein 13 |
| NM_080903 | Trim63 | 0.0422 | 0.52 | tripartite motif-containing 63 |
| NM_012680 | Tsc2 | 0.0036 | 0.74 | tuberous sclerosis 2 |
| NM_001012176 | Tsga2 | 0.0155 | 0.61 | testis specific gene A2 |
| NM_001013886 | Tubb2b | 0.0315 | 0.65 | tubulin, beta-like |
| NM_019179 | Tyms | 0.0200 | 0.74 | thymidylate synthase |
| XM_215924 | Ube2c | 0.0040 | 0.32 | similar to ubiquitin-conjugating enzyme E2C |
| NM_001008882 | Uhrf1_mapped | 0.0461 | 0.62 | ubiquitin-like, containing PHD and RING finger domains, 1 |
| AY588414 | Ush1c | 0.0418 | 0.56 | harmonin a1 |
| XM_236240 | Usp28 | 0.0039 | 0.40 | similar to ubiquitin specific protease 28 |
| NM_173112 | V1rb7 | 0.0358 | 0.67 | vomeronasal 1 receptor, B7 |
| XM_227428 | Vps72 | 0.0090 | 0.72 | similar to Vacuolar protein sorting protein 72 homolog |
| XM_227428 | Vps72 | 0.0082 | 0.73 | similar to Vacuolar protein sorting protein 72 homolog |
| NM_017154 | Xdh | 0.0031 | 0.74 | xanthine dehydrogenase |
| XM_218589 | XM_218589 | 0.0031 | 0.54 | similar to HPS5 protein major form |
| XM_218621 | XM_218621 | 0.0250 | 0.63 | predicted interleukin 4 induced 1 |
| XM_219475 | XM_219475 | 0.0153 | 0.74 | similar to cDNA sequence BC023151 |
| XM_219775 | XM_219775 | 0.0282 | 0.66 | similar to PD-1-ligand precursor |
| XM_219913 | XM_219913 | 0.0215 | 0.54 | phosphatidylinositol-4-phosphate 5-kinase, type 1, beta |
| XM_222900 | XM_222900 | 0.0009 | 0.66 | immunoglobulin superfamily, member 8 |
| XM_342715 | XM_342715 | 0.0204 | 0.73 | adducin 2 (beta) |
| XM_220618 | Ybx2 | 0.0356 | 0.44 | similar to Y-box-binding protein 2 |
| NM_001012169 | Zfp143 | 0.0007 | 0.47 | zinc finger protein 143 |
| XM_221622 | Zfp295 | 0.0412 | 0.48 | similar to zinc finger protein 295 |
| XM_230857 | Zfp334 | 0.0314 | 0.71 | hypothetical protein |
| XM_343509 | Znf167 | 0.0095 | 0.46 | similar to zinc finger protein ZFP isoform 1 |
| XM_344862 | Znf324 | 0.0237 | 0.68 | similar to zinc finger protein 324 |

**Supplementary Table 8: Functional categories found significantly enriched in the mPFC transcriptome of the PNFlx animals.**

The mPFC transcriptome of adult PNFlx animals was analyzed and regulated genes were subjected to the functional analysis using DAVID. Given in the table is a list of functional categories that were significantly upregulated in the PNFlx group.

| Gene Category | *P* Value | No. of genes regulated |
|---|---|---|
| GO:0005737~cytoplasm | 0.0000 | 88 |
| GO:0044424~intracellular part | 0.0000 | 103 |
| GO:0005622~intracellular | 0.0001 | 104 |
| GO:0006163~purine nucleotide metabolic process | 0.0001 | 10 |
| GO:0043231~intracellular membrane-bounded organelle | 0.0002 | 79 |
| GO:0043227~membrane-bounded organelle | 0.0002 | 79 |
| GO:0055086~nucleobase, nucleoside and nucleotide metabolic process | 0.0002 | 12 |
| GO:0065007~biological regulation | 0.0003 | 70 |
| GO:0051179~localization | 0.0004 | 40 |
| GO:0009117~nucleotide metabolic process | 0.0004 | 11 |
| GO:0006753~nucleoside phosphate metabolic process | 0.0004 | 11 |
| GO:0022892~substrate-specific transporter activity | 0.0006 | 19 |
| GO:0051239~regulation of multicellular organismal process | 0.0007 | 21 |
| GO:0006139~nucleobase, nucleoside, nucleotide and nucleic acid metabolic process | 0.0009 | 30 |
| GO:0043229~intracellular organelle | 0.0009 | 85 |
| GO:0005515~protein binding | 0.0009 | 82 |
| GO:0043226~organelle | 0.0010 | 85 |
| GO:0006811~ion transport | 0.0012 | 16 |
| GO:0046483~heterocycle metabolic process | 0.0013 | 11 |
| GO:0050789~regulation of biological process | 0.0013 | 64 |
| GO:0007611~learning or memory | 0.0014 | 7 |
| GO:0045471~response to ethanol | 0.0015 | 6 |
| GO:0015672~monovalent inorganic cation transport | 0.0015 | 10 |
| GO:0031420~alkali metal ion binding | 0.0016 | 8 |
| GO:0008324~cation transmembrane transporter activity | 0.0016 | 13 |
| GO:0040008~regulation of growth | 0.0018 | 10 |
| GO:0015075~ion transmembrane transporter activity | 0.0019 | 15 |
| GO:0006812~cation transport | 0.0019 | 13 |
| GO:0022857~transmembrane transporter activity | 0.0021 | 17 |
| GO:0009150~purine ribonucleotide metabolic process | 0.0022 | 7 |
| GO:0030828~positive regulation of cGMP biosynthetic process | 0.0023 | 3 |
| GO:0030825~positive regulation of cGMP metabolic process | 0.0023 | 3 |
| GO:0005829~cytosol | 0.0024 | 22 |
| GO:0031967~organelle envelope | 0.0025 | 14 |
| GO:0022891~substrate-specific transmembrane transporter activity | 0.0025 | 16 |
| GO:0006810~transport | 0.0026 | 33 |
| GO:0031975~envelope | 0.0026 | 14 |
| GO:0005215~transporter activity | 0.0026 | 20 |

| | | |
|---|---|---|
| GO:0009259~ribonucleotide metabolic process | 0.0027 | 7 |
| GO:0044057~regulation of system process | 0.0028 | 10 |
| GO:0034641~cellular nitrogen compound metabolic process | 0.0030 | 31 |
| GO:0044422~organelle part | 0.0030 | 46 |
| GO:0007165~signal transduction | 0.0030 | 27 |
| GO:0051234~establishment of localization | 0.0031 | 33 |
| GO:0009987~cellular process | 0.0031 | 86 |
| GO:0005102~receptor binding | 0.0031 | 16 |
| GO:0044444~cytoplasmic part | 0.0032 | 59 |
| GO:0006807~nitrogen compound metabolic process | 0.0034 | 32 |
| GO:0006164~purine nucleotide biosynthetic process | 0.0034 | 7 |
| GO:0030826~regulation of cGMP biosynthetic process | 0.0036 | 3 |
| GO:0030001~metal ion transport | 0.0041 | 11 |
| GO:0005739~mitochondrion | 0.0042 | 23 |
| GO:0030823~regulation of cGMP metabolic process | 0.0044 | 3 |
| GO:0044446~intracellular organelle part | 0.0045 | 45 |
| GO:0048519~negative regulation of biological process | 0.0052 | 26 |
| GO:0065008~regulation of biological quality | 0.0055 | 23 |
| GO:0032502~developmental process | 0.0058 | 37 |
| GO:0043502~regulation of muscle adaptation | 0.0062 | 3 |
| GO:0009144~purine nucleoside triphosphate metabolic process | 0.0071 | 6 |
| GO:0031966~mitochondrial membrane | 0.0086 | 10 |
| GO:0009141~nucleoside triphosphate metabolic process | 0.0088 | 6 |
| GO:0009165~nucleotide biosynthetic process | 0.0090 | 7 |
| GO:0031090~organelle membrane | 0.0092 | 18 |
| GO:0009743~response to carbohydrate stimulus | 0.0096 | 5 |
| GO:0031402~sodium ion binding | 0.0097 | 5 |
| GO:0030234~enzyme regulator activity | 0.0101 | 13 |
| GO:0034654~nucleobase, nucleoside, nucleotide and nucleic acid biosynthetic process | 0.0106 | 7 |
| GO:0034404~nucleobase, nucleoside and nucleotide biosynthetic process | 0.0106 | 7 |
| GO:0005802~trans-Golgi network | 0.0112 | 4 |
| GO:0010033~response to organic substance | 0.0112 | 17 |
| GO:0050794~regulation of cellular process | 0.0114 | 57 |
| GO:0007568~aging | 0.0126 | 6 |
| GO:0048523~negative regulation of cellular process | 0.0127 | 23 |
| GO:0005740~mitochondrial envelope | 0.0129 | 10 |
| GO:0048639~positive regulation of developmental growth | 0.0144 | 3 |
| GO:0005626~insoluble fraction | 0.0145 | 15 |
| GO:0051241~negative regulation of multicellular organismal process | 0.0155 | 6 |
| GO:0007613~memory | 0.0156 | 4 |
| GO:0030801~positive regulation of cyclic nucleotide metabolic process | 0.0158 | 3 |
| GO:0030804~positive regulation of cyclic nucleotide biosynthetic process | 0.0158 | 3 |
| GO:0045981~positive regulation of nucleotide metabolic process | 0.0158 | 3 |
| GO:0030810~positive regulation of nucleotide biosynthetic process | 0.0158 | 3 |
| GO:0005488~binding | 0.0163 | 108 |
| GO:0030955~potassium ion binding | 0.0166 | 5 |
| GO:0046873~metal ion transmembrane transporter activity | 0.0170 | 8 |

| | | |
|---|---|---|
| GO:0004437~inositol or phosphatidylinositol phosphatase activity | 0.0178 | 3 |
| GO:0048545~response to steroid hormone stimulus | 0.0183 | 8 |
| GO:0006814~sodium ion transport | 0.0183 | 5 |
| GO:0005244~voltage-gated ion channel activity | 0.0189 | 6 |
| GO:0022832~voltage-gated channel activity | 0.0189 | 6 |
| GO:0009725~response to hormone stimulus | 0.0195 | 11 |
| GO:0022804~active transmembrane transporter activity | 0.0209 | 8 |
| GO:0005624~membrane fraction | 0.0213 | 14 |
| GO:0044087~regulation of cellular component biogenesis | 0.0218 | 5 |
| GO:0009154~purine ribonucleotide catabolic process | 0.0221 | 3 |
| GO:0007242~intracellular signaling cascade | 0.0230 | 16 |
| GO:0009261~ribonucleotide catabolic process | 0.0238 | 3 |
| GO:0000267~cell fraction | 0.0244 | 17 |
| GO:0005261~cation channel activity | 0.0247 | 7 |
| GO:0000904~cell morphogenesis involved in differentiation | 0.0247 | 7 |
| GO:0050793~regulation of developmental process | 0.0252 | 13 |
| GO:0043167~ion binding | 0.0257 | 37 |
| GO:0030315~T-tubule | 0.0260 | 3 |
| GO:0046872~metal ion binding | 0.0271 | 36 |
| GO:0032990~cell part morphogenesis | 0.0274 | 7 |
| GO:0009152~purine ribonucleotide biosynthetic process | 0.0278 | 5 |
| GO:0044429~mitochondrial part | 0.0279 | 11 |
| GO:0009205~purine ribonucleoside triphosphate metabolic process | 0.0285 | 5 |
| GO:0022843~voltage-gated cation channel activity | 0.0288 | 5 |
| GO:0009749~response to glucose stimulus | 0.0289 | 4 |
| GO:0007612~learning | 0.0289 | 4 |
| GO:0051270~regulation of cell motion | 0.0290 | 6 |
| GO:0009199~ribonucleoside triphosphate metabolic process | 0.0293 | 5 |
| GO:0032879~regulation of localization | 0.0299 | 12 |
| GO:0009260~ribonucleotide biosynthetic process | 0.0308 | 5 |
| GO:0044238~primary metabolic process | 0.0316 | 57 |
| GO:0005216~ion channel activity | 0.0316 | 8 |
| GO:0042493~response to drug | 0.0319 | 8 |
| GO:0001666~response to hypoxia | 0.0320 | 6 |
| GO:0043169~cation binding | 0.0325 | 36 |
| GO:0044271~nitrogen compound biosynthetic process | 0.0328 | 8 |
| GO:0009746~response to hexose stimulus | 0.0332 | 4 |
| GO:0034284~response to monosaccharide stimulus | 0.0332 | 4 |
| GO:0044237~cellular metabolic process | 0.0340 | 54 |
| GO:0022836~gated channel activity | 0.0342 | 7 |
| GO:0031981~nuclear lumen | 0.0343 | 16 |
| GO:0070013~intracellular organelle lumen | 0.0348 | 19 |
| GO:0043200~response to amino acid stimulus | 0.0350 | 3 |
| GO:0022838~substrate specific channel activity | 0.0362 | 8 |
| GO:0014874~response to stimulus involved in regulation of muscle adaptation | 0.0364 | 2 |
| GO:0008219~cell death | 0.0374 | 9 |
| GO:0022890~inorganic cation transmembrane transporter activity | 0.0380 | 5 |

| | | |
|---|---|---|
| GO:0048878~chemical homeostasis | 0.0384 | 10 |
| GO:0006928~cell motion | 0.0388 | 9 |
| GO:0005509~calcium ion binding | 0.0389 | 12 |
| GO:0009719~response to endogenous stimulus | 0.0391 | 11 |
| GO:0042383~sarcolemma | 0.0399 | 4 |
| GO:0070482~response to oxygen levels | 0.0399 | 6 |
| GO:0016265~death | 0.0407 | 9 |
| GO:0031175~neuron projection development | 0.0409 | 7 |
| GO:0050680~negative regulation of epithelial cell proliferation | 0.0413 | 3 |
| GO:0032355~response to estradiol stimulus | 0.0414 | 4 |
| GO:0006813~potassium ion transport | 0.0425 | 5 |
| GO:0007610~behavior | 0.0426 | 9 |
| GO:0015267~channel activity | 0.0428 | 8 |
| GO:0022803~passive transmembrane transporter activity | 0.0428 | 8 |
| GO:0048667~cell morphogenesis involved in neuron differentiation | 0.0428 | 6 |
| GO:0048468~cell development | 0.0437 | 12 |
| GO:0016528~sarcoplasm | 0.0450 | 3 |
| GO:0033279~ribosomal subunit | 0.0466 | 4 |
| GO:0043233~organelle lumen | 0.0469 | 19 |
| GO:0043627~response to estrogen stimulus | 0.0482 | 5 |
| GO:0048812~neuron projection morphogenesis | 0.0490 | 6 |
| GO:0044455~mitochondrial membrane part | 0.0494 | 4 |
| GO:0015078~hydrogen ion transmembrane transporter activity | 0.0497 | 4 |
| GO:0000287~magnesium ion binding | 0.0497 | 7 |
| GO:0022627~cytosolic small ribosomal subunit | 0.0497 | 3 |

**Supplementary Table 9: Functional categories found significantly downregulated in the mPFC transcriptome of the PNFlx animals.**

The mPFC transcriptome of adult PNFlx animals was analyzed and regulated genes were subjected to the functional analysis using DAVID. Given in the table is a list of functional categories that were significantly downregulated in the PNFlx group.

| Gene Category | *P* Value | No. of genes regulated |
|---|---|---|
| GO:0007612~learning | 0.0007 | 6 |
| GO:0030246~carbohydrate binding | 0.0014 | 11 |
| GO:0007611~learning or memory | 0.0019 | 7 |
| GO:0045211~postsynaptic membrane | 0.0022 | 7 |
| GO:0022803~passive transmembrane transporter activity | 0.0023 | 11 |
| GO:0015267~channel activity | 0.0023 | 11 |
| GO:0010646~regulation of cell communication | 0.0025 | 20 |
| GO:0000267~cell fraction | 0.0026 | 20 |
| GO:0005737~cytoplasm | 0.0030 | 75 |
| GO:0044444~cytoplasmic part | 0.0040 | 59 |
| GO:0010035~response to inorganic substance | 0.0051 | 9 |
| GO:0019904~protein domain specific binding | 0.0053 | 10 |
| GO:0005515~protein binding | 0.0058 | 83 |
| GO:0022838~substrate specific channel activity | 0.0059 | 10 |
| GO:0032496~response to lipopolysaccharide | 0.0060 | 6 |
| GO:0005488~binding | 0.0061 | 116 |
| GO:0015837~amine transport | 0.0069 | 6 |
| GO:0010648~negative regulation of cell communication | 0.0078 | 8 |
| GO:0002237~response to molecule of bacterial origin | 0.0087 | 6 |
| GO:0022857~transmembrane transporter activity | 0.0092 | 16 |
| GO:0048878~chemical homeostasis | 0.0094 | 12 |
| GO:0010033~response to organic substance | 0.0096 | 18 |
| GO:0043005~neuron projection | 0.0103 | 11 |
| GO:0042995~cell projection | 0.0114 | 15 |
| GO:0043170~macromolecule metabolic process | 0.0123 | 51 |
| GO:0044456~synapse part | 0.0135 | 8 |
| GO:0016070~RNA metabolic process | 0.0140 | 13 |
| GO:0065007~biological regulation | 0.0145 | 67 |
| GO:0006811~ion transport | 0.0146 | 14 |
| GO:0045744~negative regulation of G-protein coupled receptor protein signaling pathway | 0.0148 | 3 |
| GO:0007568~aging | 0.0164 | 6 |
| GO:0032147~activation of protein kinase activity | 0.0166 | 5 |
| GO:0016917~GABA receptor activity | 0.0168 | 3 |
| GO:0050796~regulation of insulin secretion | 0.0170 | 4 |
| GO:0016791~phosphatase activity | 0.0173 | 7 |
| GO:0044422~organelle part | 0.0174 | 43 |
| GO:0046676~negative regulation of insulin secretion | 0.0179 | 3 |
| GO:0002792~negative regulation of peptide secretion | 0.0179 | 3 |

| | | |
|---|---|---|
| GO:0051241~negative regulation of multicellular organismal process | 0.0201 | 6 |
| GO:0031090~organelle membrane | 0.0208 | 17 |
| GO:0005886~plasma membrane | 0.0213 | 34 |
| GO:0002791~regulation of peptide secretion | 0.0223 | 4 |
| GO:0006812~cation transport | 0.0237 | 11 |
| GO:0042592~homeostatic process | 0.0238 | 14 |
| GO:0022891~substrate-specific transmembrane transporter activity | 0.0239 | 14 |
| GO:0044260~cellular macromolecule metabolic process | 0.0240 | 44 |
| GO:0044446~intracellular organelle part | 0.0242 | 42 |
| GO:0007610~behavior | 0.0244 | 10 |
| GO:0015672~monovalent inorganic cation transport | 0.0248 | 8 |
| GO:0043197~dendritic spine | 0.0256 | 4 |
| GO:0048523~negative regulation of cellular process | 0.0259 | 23 |
| GO:0003723~RNA binding | 0.0268 | 10 |
| GO:0005625~soluble fraction | 0.0268 | 8 |
| GO:0015850~organic alcohol transport | 0.0269 | 3 |
| GO:0001932~regulation of protein amino acid phosphorylation | 0.0271 | 6 |
| GO:0032502~developmental process | 0.0285 | 36 |
| GO:0009725~response to hormone stimulus | 0.0293 | 11 |
| GO:0045202~synapse | 0.0322 | 9 |
| GO:0050794~regulation of cellular process | 0.0324 | 58 |
| GO:0005739~mitochondrion | 0.0342 | 20 |
| GO:0031420~alkali metal ion binding | 0.0345 | 6 |
| GO:0016208~AMP binding | 0.0349 | 3 |
| GO:0009966~regulation of signal transduction | 0.0349 | 14 |
| GO:0050789~regulation of biological process | 0.0354 | 61 |
| GO:0009607~response to biotic stimulus | 0.0370 | 8 |
| GO:0044424~intracellular part | 0.0372 | 90 |
| GO:0006350~transcription | 0.0375 | 13 |
| GO:0008282~ATP-sensitive potassium channel complex | 0.0385 | 2 |
| GO:0006810~transport | 0.0388 | 30 |
| GO:0045860~positive regulation of protein kinase activity | 0.0407 | 6 |
| GO:0065008~regulation of biological quality | 0.0408 | 21 |
| GO:0048518~positive regulation of biological process | 0.0411 | 27 |
| GO:0005216~ion channel activity | 0.0417 | 8 |
| GO:0051179~localization | 0.0419 | 34 |
| GO:0051707~response to other organism | 0.0426 | 7 |
| GO:0005215~transporter activity | 0.0426 | 17 |
| GO:0051234~establishment of localization | 0.0434 | 30 |
| GO:0008152~metabolic process | 0.0443 | 67 |
| GO:0030001~metal ion transport | 0.0461 | 9 |
| GO:0033674~positive regulation of kinase activity | 0.0472 | 6 |
| GO:0009968~negative regulation of signal transduction | 0.0480 | 6 |
| GO:0007260~tyrosine phosphorylation of STAT protein | 0.0484 | 2 |
| GO:0042277~peptide binding | 0.0487 | 6 |
| GO:0043227~membrane-bounded organelle | 0.0487 | 69 |
| GO:0043566~structure-specific DNA binding | 0.0487 | 5 |
| GO:0046888~negative regulation of hormone secretion | 0.0489 | 3 |

| | | |
|---|---|---|
| GO:0042578~phosphoric ester hydrolase activity | 0.0499 | 7 |

**Supplementary Table 10: Functional categories found significantly enriched in the mPFC transcriptome of the JFlx animals.**

The mPFC transcriptome of adult JFlx animals was analyzed and regulated genes were subjected to the functional analysis using DAVID. Given in the table is a list of functional categories that were significantly upregulated in the JFlx group.

| **Gene Category** | ***P Value*** | **No. of genes regulated** |
|---|---|---|
| GO:0009987~cellular process | 0.0000 | 114 |
| GO:0005515~protein binding | 0.0000 | 105 |
| GO:0022008~neurogenesis | 0.0000 | 25 |
| GO:0030154~cell differentiation | 0.0000 | 39 |
| GO:0048869~cellular developmental process | 0.0000 | 39 |
| GO:0045202~synapse | 0.0000 | 18 |
| GO:0007399~nervous system development | 0.0000 | 29 |
| GO:0048699~generation of neurons | 0.0000 | 22 |
| GO:0022604~regulation of cell morphogenesis | 0.0000 | 11 |
| GO:0032502~developmental process | 0.0000 | 52 |
| GO:0048468~cell development | 0.0000 | 23 |
| GO:0005886~plasma membrane | 0.0000 | 50 |
| GO:0044459~plasma membrane part | 0.0000 | 34 |
| GO:0048856~anatomical structure development | 0.0000 | 45 |
| GO:0010975~regulation of neuron projection development | 0.0000 | 9 |
| GO:0010769~regulation of cell morphogenesis involved in differentiation | 0.0000 | 9 |
| GO:0000904~cell morphogenesis involved in differentiation | 0.0000 | 13 |
| GO:0005488~binding | 0.0000 | 132 |
| GO:0007275~multicellular organismal development | 0.0000 | 46 |
| GO:0005829~cytosol | 0.0000 | 29 |
| GO:0051128~regulation of cellular component organization | 0.0000 | 16 |
| GO:0048731~system development | 0.0000 | 42 |
| GO:0050768~negative regulation of neurogenesis | 0.0000 | 7 |
| GO:0031344~regulation of cell projection organization | 0.0000 | 9 |
| GO:0030030~cell projection organization | 0.0000 | 15 |
| GO:0010721~negative regulation of cell development | 0.0000 | 7 |
| GO:0042995~cell projection | 0.0000 | 22 |
| GO:0031345~negative regulation of cell projection organization | 0.0000 | 6 |
| GO:0050767~regulation of neurogenesis | 0.0001 | 11 |
| GO:0045664~regulation of neuron differentiation | 0.0001 | 10 |
| GO:0048523~negative regulation of cellular process | 0.0001 | 32 |
| GO:0030054~cell junction | 0.0001 | 16 |
| GO:0050770~regulation of axonogenesis | 0.0001 | 7 |
| GO:0017076~purine nucleotide binding | 0.0001 | 34 |
| GO:0044456~synapse part | 0.0001 | 12 |
| GO:0001883~purine nucleoside binding | 0.0001 | 30 |
| GO:0001882~nucleoside binding | 0.0001 | 30 |
| GO:0031175~neuron projection development | 0.0001 | 12 |

| | | |
|---|---|---|
| GO:0051960~regulation of nervous system development | 0.0001 | 11 |
| GO:0050794~regulation of cellular process | 0.0001 | 71 |
| GO:0032989~cellular component morphogenesis | 0.0001 | 14 |
| GO:0048519~negative regulation of biological process | 0.0002 | 33 |
| GO:0060284~regulation of cell development | 0.0002 | 11 |
| GO:0046777~protein amino acid autophosphorylation | 0.0002 | 7 |
| GO:0022603~regulation of anatomical structure morphogenesis | 0.0002 | 11 |
| GO:0005737~cytoplasm | 0.0002 | 86 |
| GO:0019226~transmission of nerve impulse | 0.0002 | 12 |
| GO:0032555~purine ribonucleotide binding | 0.0002 | 32 |
| GO:0032553~ribonucleotide binding | 0.0002 | 32 |
| GO:0030554~adenyl nucleotide binding | 0.0002 | 29 |
| GO:0065007~biological regulation | 0.0002 | 78 |
| GO:0000902~cell morphogenesis | 0.0002 | 13 |
| GO:0043412~biopolymer modification | 0.0002 | 28 |
| GO:0050771~negative regulation of axonogenesis | 0.0003 | 5 |
| GO:0006464~protein modification process | 0.0003 | 27 |
| GO:0032879~regulation of localization | 0.0003 | 18 |
| GO:0048666~neuron development | 0.0003 | 13 |
| GO:0030182~neuron differentiation | 0.0003 | 15 |
| GO:0051129~negative regulation of cellular component organization | 0.0003 | 8 |
| GO:0045596~negative regulation of cell differentiation | 0.0004 | 10 |
| GO:0043687~post-translational protein modification | 0.0004 | 24 |
| GO:0032559~adenyl ribonucleotide binding | 0.0005 | 27 |
| GO:0007268~synaptic transmission | 0.0005 | 10 |
| GO:0003873~6-phosphofructo-2-kinase activity | 0.0006 | 3 |
| GO:0044444~cytoplasmic part | 0.0006 | 67 |
| GO:0051179~localization | 0.0006 | 43 |
| GO:0051641~cellular localization | 0.0007 | 19 |
| GO:0050789~regulation of biological process | 0.0007 | 72 |
| GO:0005794~Golgi apparatus | 0.0007 | 18 |
| GO:0019904~protein domain specific binding | 0.0008 | 12 |
| GO:0005524~ATP binding | 0.0008 | 26 |
| GO:0050793~regulation of developmental process | 0.0008 | 18 |
| GO:0045595~regulation of cell differentiation | 0.0008 | 15 |
| GO:0010646~regulation of cell communication | 0.0008 | 22 |
| GO:0009653~anatomical structure morphogenesis | 0.0008 | 24 |
| GO:0042802~identical protein binding | 0.0010 | 16 |
| GO:0004331~fructose-2,6-bisphosphate 2-phosphatase activity | 0.0010 | 3 |
| GO:0030001~metal ion transport | 0.0010 | 13 |
| GO:0006003~fructose 2,6-bisphosphate metabolic process | 0.0011 | 3 |
| GO:0000166~nucleotide binding | 0.0011 | 35 |
| GO:0016773~phosphotransferase activity, alcohol group as acceptor | 0.0012 | 17 |
| GO:0044446~intracellular organelle part | 0.0012 | 51 |
| GO:0051093~negative regulation of developmental process | 0.0014 | 10 |
| GO:0042733~embryonic digit morphogenesis | 0.0015 | 4 |
| GO:0044422~organelle part | 0.0015 | 51 |
| GO:0019899~enzyme binding | 0.0016 | 14 |

| | | |
|---|---|---|
| GO:0016020~membrane | 0.0017 | 86 |
| GO:0016301~kinase activity | 0.0017 | 18 |
| GO:0009898~internal side of plasma membrane | 0.0018 | 8 |
| GO:0044424~intracellular part | 0.0018 | 104 |
| GO:0007269~neurotransmitter secretion | 0.0021 | 5 |
| GO:0008443~phosphofructokinase activity | 0.0021 | 3 |
| GO:0016043~cellular component organization | 0.0023 | 34 |
| GO:0043005~neuron projection | 0.0025 | 13 |
| GO:0007167~enzyme linked receptor protein signaling pathway | 0.0026 | 10 |
| GO:0007154~cell communication | 0.0028 | 14 |
| GO:0016772~transferase activity, transferring phosphorus-containing groups | 0.0028 | 19 |
| GO:0065008~regulation of biological quality | 0.0029 | 26 |
| GO:0006796~phosphate metabolic process | 0.0030 | 19 |
| GO:0051239~regulation of multicellular organismal process | 0.0031 | 21 |
| GO:0051049~regulation of transport | 0.0031 | 13 |
| GO:0006793~phosphorus metabolic process | 0.0031 | 19 |
| GO:0031420~alkali metal ion binding | 0.0033 | 8 |
| GO:0007409~axonogenesis | 0.0033 | 8 |
| GO:0016044~membrane organization | 0.0033 | 10 |
| GO:0045211~postsynaptic membrane | 0.0035 | 7 |
| GO:0051234~establishment of localization | 0.0035 | 36 |
| GO:0019203~carbohydrate phosphatase activity | 0.0036 | 3 |
| GO:0006813~potassium ion transport | 0.0037 | 7 |
| GO:0046872~metal ion binding | 0.0039 | 44 |
| GO:0051649~establishment of localization in cell | 0.0046 | 16 |
| GO:0043169~cation binding | 0.0050 | 44 |
| GO:0006468~protein amino acid phosphorylation | 0.0050 | 15 |
| GO:0004672~protein kinase activity | 0.0051 | 14 |
| GO:0030955~potassium ion binding | 0.0052 | 6 |
| GO:0006812~cation transport | 0.0053 | 13 |
| GO:0031090~organelle membrane | 0.0053 | 20 |
| GO:0006810~transport | 0.0056 | 35 |
| GO:0019898~extrinsic to membrane | 0.0059 | 10 |
| GO:0048667~cell morphogenesis involved in neuron differentiation | 0.0059 | 8 |
| GO:0001501~skeletal system development | 0.0064 | 9 |
| GO:0016791~phosphatase activity | 0.0067 | 8 |
| GO:0043167~ion binding | 0.0068 | 44 |
| GO:0048812~neuron projection morphogenesis | 0.0073 | 8 |
| GO:0042063~gliogenesis | 0.0074 | 5 |
| GO:0007242~intracellular signaling cascade | 0.0074 | 19 |
| GO:0050804~regulation of synaptic transmission | 0.0076 | 7 |
| GO:0007267~cell-cell signaling | 0.0076 | 10 |
| GO:0042578~phosphoric ester hydrolase activity | 0.0078 | 9 |
| GO:0044425~membrane part | 0.0087 | 73 |
| GO:0050769~positive regulation of neurogenesis | 0.0093 | 5 |
| GO:0065009~regulation of molecular function | 0.0095 | 17 |
| GO:0007264~small GTPase mediated signal transduction | 0.0099 | 8 |

| | | |
|---|---|---|
| GO:0051969~regulation of transmission of nerve impulse | 0.0102 | 7 |
| GO:0001837~epithelial to mesenchymal transition | 0.0104 | 3 |
| GO:0005083~small GTPase regulator activity | 0.0104 | 7 |
| GO:0016358~dendrite development | 0.0107 | 4 |
| GO:0015672~monovalent inorganic cation transport | 0.0109 | 9 |
| GO:0001505~regulation of neurotransmitter levels | 0.0114 | 5 |
| GO:0016055~Wnt receptor signaling pathway | 0.0114 | 5 |
| GO:0000785~chromatin | 0.0115 | 7 |
| GO:0005622~intracellular | 0.0116 | 106 |
| GO:0006928~cell motion | 0.0117 | 11 |
| GO:0006000~fructose metabolic process | 0.0118 | 3 |
| GO:0005741~mitochondrial outer membrane | 0.0119 | 5 |
| GO:0014069~postsynaptic density | 0.0119 | 5 |
| GO:0031252~cell leading edge | 0.0120 | 6 |
| GO:0048858~cell projection morphogenesis | 0.0123 | 8 |
| GO:0040008~regulation of growth | 0.0127 | 9 |
| GO:0019200~carbohydrate kinase activity | 0.0127 | 3 |
| GO:0030425~dendrite | 0.0128 | 8 |
| GO:0051338~regulation of transferase activity | 0.0130 | 9 |
| GO:0010720~positive regulation of cell development | 0.0134 | 5 |
| GO:0016740~transferase activity | 0.0134 | 26 |
| GO:0043231~intracellular membrane-bounded organelle | 0.0137 | 78 |
| GO:0031644~regulation of neurological system process | 0.0138 | 7 |
| GO:0008022~protein C-terminus binding | 0.0140 | 6 |
| GO:0043227~membrane-bounded organelle | 0.0143 | 78 |
| GO:0007283~spermatogenesis | 0.0151 | 8 |
| GO:0048232~male gamete generation | 0.0151 | 8 |
| GO:0032990~cell part morphogenesis | 0.0151 | 8 |
| GO:0051018~protein kinase A binding | 0.0158 | 3 |
| GO:0048513~organ development | 0.0165 | 27 |
| GO:0016310~phosphorylation | 0.0169 | 15 |
| GO:0010629~negative regulation of gene expression | 0.0169 | 11 |
| GO:0048609~reproductive process in a multicellular organism | 0.0174 | 11 |
| GO:0032504~multicellular organism reproduction | 0.0174 | 11 |
| GO:0001726~ruffle | 0.0180 | 4 |
| GO:0044057~regulation of system process | 0.0185 | 9 |
| GO:0042476~odontogenesis | 0.0186 | 4 |
| GO:0012505~endomembrane system | 0.0191 | 14 |
| GO:0045934~negative regulation of nucleobase, nucleoside, nucleotide and nucleic acid metabolic process | 0.0195 | 11 |
| GO:0008104~protein localization | 0.0197 | 14 |
| GO:0007276~gamete generation | 0.0201 | 9 |
| GO:0031968~organelle outer membrane | 0.0201 | 5 |
| GO:0015379~potassium:chloride symporter activity | 0.0203 | 2 |
| GO:0006836~neurotransmitter transport | 0.0205 | 5 |
| GO:0008285~negative regulation of cell proliferation | 0.0205 | 8 |
| GO:0042127~regulation of cell proliferation | 0.0213 | 14 |

| | | |
|---|---|---|
| GO:0051172~negative regulation of nitrogen compound metabolic process | 0.0214 | 11 |
| GO:0048646~anatomical structure formation involved in morphogenesis | 0.0214 | 9 |
| GO:0005215~transporter activity | 0.0216 | 19 |
| GO:0045926~negative regulation of growth | 0.0219 | 5 |
| GO:0005516~calmodulin binding | 0.0219 | 5 |
| GO:0022414~reproductive process | 0.0224 | 14 |
| GO:0000139~Golgi membrane | 0.0229 | 6 |
| GO:0019867~outer membrane | 0.0229 | 5 |
| GO:0043066~negative regulation of apoptosis | 0.0232 | 9 |
| GO:0007215~glutamate signaling pathway | 0.0236 | 3 |
| GO:0042325~regulation of phosphorylation | 0.0237 | 10 |
| GO:0007411~axon guidance | 0.0240 | 5 |
| GO:0000003~reproduction | 0.0245 | 14 |
| GO:0043069~negative regulation of programmed cell death | 0.0251 | 9 |
| GO:0060548~negative regulation of cell death | 0.0255 | 9 |
| GO:0003001~generation of a signal involved in cell-cell signaling | 0.0255 | 5 |
| GO:0016481~negative regulation of transcription | 0.0265 | 10 |
| GO:0050790~regulation of catalytic activity | 0.0266 | 14 |
| GO:0006814~sodium ion transport | 0.0271 | 5 |
| GO:0019538~protein metabolic process | 0.0273 | 36 |
| GO:0014013~regulation of gliogenesis | 0.0276 | 3 |
| GO:0010558~negative regulation of macromolecule biosynthetic process | 0.0284 | 11 |
| GO:0051174~regulation of phosphorus metabolic process | 0.0295 | 10 |
| GO:0019220~regulation of phosphate metabolic process | 0.0295 | 10 |
| GO:0001503~ossification | 0.0295 | 5 |
| GO:0043549~regulation of kinase activity | 0.0296 | 8 |
| GO:0005626~insoluble fraction | 0.0308 | 15 |
| GO:0044267~cellular protein metabolic process | 0.0313 | 30 |
| GO:0048729~tissue morphogenesis | 0.0315 | 7 |
| GO:0001558~regulation of cell growth | 0.0318 | 6 |
| GO:0031327~negative regulation of cellular biosynthetic process | 0.0325 | 11 |
| GO:0005911~cell-cell junction | 0.0333 | 6 |
| GO:0004713~protein tyrosine kinase activity | 0.0336 | 5 |
| GO:0005694~chromosome | 0.0340 | 9 |
| GO:0009890~negative regulation of biosynthetic process | 0.0365 | 11 |
| GO:0031982~vesicle | 0.0375 | 13 |
| GO:0044260~cellular macromolecule metabolic process | 0.0383 | 45 |
| GO:0043226~organelle | 0.0383 | 85 |
| GO:0005634~nucleus | 0.0383 | 46 |
| GO:0019905~syntaxin binding | 0.0397 | 3 |
| GO:0060348~bone development | 0.0406 | 5 |
| GO:0060666~dichotomous subdivision of terminal units involved in salivary gland branching | 0.0410 | 2 |
| GO:0048843~negative regulation of axon extension involved in axon guidance | 0.0410 | 2 |
| GO:0004721~phosphoprotein phosphatase activity | 0.0412 | 5 |
| GO:0019953~sexual reproduction | 0.0413 | 9 |

| | | |
|---|---|---|
| GO:0031625~ubiquitin protein ligase binding | 0.0421 | 3 |
| GO:0048471~perinuclear region of cytoplasm | 0.0423 | 7 |
| GO:0005509~calcium ion binding | 0.0425 | 13 |
| GO:0031323~regulation of cellular metabolic process | 0.0427 | 34 |
| GO:0008287~protein serine/threonine phosphatase complex | 0.0430 | 3 |
| GO:0050772~positive regulation of axonogenesis | 0.0436 | 3 |
| GO:0007165~signal transduction | 0.0455 | 25 |
| GO:0006811~ion transport | 0.0459 | 13 |
| GO:0033036~macromolecule localization | 0.0460 | 15 |
| GO:0021954~central nervous system neuron development | 0.0461 | 3 |
| GO:0043232~intracellular non-membrane-bounded organelle | 0.0464 | 28 |
| GO:0043228~non-membrane-bounded organelle | 0.0464 | 28 |
| GO:0051130~positive regulation of cellular component organization | 0.0468 | 6 |
| GO:0032501~multicellular organismal process | 0.0471 | 56 |
| GO:0044427~chromosomal part | 0.0478 | 8 |
| GO:0007265~Ras protein signal transduction | 0.0478 | 4 |
| GO:0004674~protein serine/threonine kinase activity | 0.0486 | 9 |
| GO:0030509~BMP signaling pathway | 0.0487 | 3 |
| GO:0008283~cell proliferation | 0.0490 | 7 |
| GO:0009100~glycoprotein metabolic process | 0.0491 | 5 |
| GO:0030308~negative regulation of cell growth | 0.0493 | 4 |
| GO:0042981~regulation of apoptosis | 0.0499 | 13 |

**Supplementary Table 11: Functional categories found significantly downregulated in the mPFC transcriptome of the JFlx animals.**

The mPFC transcriptome of adult JFlx animals was analyzed and regulated genes were subjected to the functional analysis using DAVID. Given in the table is a list of functional categories that were significantly downregulated in the JFlx group.

| Gene Category | *P* Value | No. of genes regulated |
|---|---|---|
| GO:0005886~plasma membrane | 0.0000 | 44 |
| GO:0048878~chemical homeostasis | 0.0001 | 16 |
| GO:0050801~ion homeostasis | 0.0001 | 14 |
| GO:0044444~cytoplasmic part | 0.0001 | 65 |
| GO:0005773~vacuole | 0.0001 | 10 |
| GO:0005737~cytoplasm | 0.0001 | 80 |
| GO:0065008~regulation of biological quality | 0.0002 | 27 |
| GO:0005488~binding | 0.0002 | 109 |
| GO:0044422~organelle part | 0.0003 | 50 |
| GO:0043226~organelle | 0.0004 | 87 |
| GO:0051240~positive regulation of multicellular organismal process | 0.0005 | 10 |
| GO:0006873~cellular ion homeostasis | 0.0007 | 12 |
| GO:0003073~regulation of systemic arterial blood pressure | 0.0007 | 5 |
| GO:0043229~intracellular organelle | 0.0007 | 86 |
| GO:0042995~cell projection | 0.0007 | 18 |
| GO:0055082~cellular chemical homeostasis | 0.0007 | 12 |
| GO:0044424~intracellular part | 0.0009 | 97 |
| GO:0042592~homeostatic process | 0.0009 | 17 |
| GO:0007154~cell communication | 0.0010 | 14 |
| GO:0008289~lipid binding | 0.0011 | 11 |
| GO:0001990~regulation of systemic arterial blood pressure by hormone | 0.0012 | 4 |
| GO:0044446~intracellular organelle part | 0.0016 | 47 |
| GO:0050886~endocrine process | 0.0018 | 4 |
| GO:0007610~behavior | 0.0020 | 12 |
| GO:0009991~response to extracellular stimulus | 0.0020 | 10 |
| GO:0001653~peptide receptor activity | 0.0021 | 6 |
| GO:0008528~peptide receptor activity, G-protein coupled | 0.0021 | 6 |
| GO:0042277~peptide binding | 0.0021 | 8 |
| GO:0003044~regulation of systemic arterial blood pressure mediated by a chemical signal | 0.0027 | 4 |
| GO:0003014~renal system process | 0.0029 | 4 |
| GO:0008217~regulation of blood pressure | 0.0031 | 6 |
| GO:0055080~cation homeostasis | 0.0031 | 9 |
| GO:0019725~cellular homeostasis | 0.0033 | 12 |
| GO:0009605~response to external stimulus | 0.0036 | 17 |
| GO:0017046~peptide hormone binding | 0.0038 | 4 |
| GO:0043167~ion binding | 0.0039 | 39 |
| GO:0048731~system development | 0.0040 | 32 |

| | | |
|---|---|---|
| GO:0042802~identical protein binding | 0.0045 | 13 |
| GO:0009987~cellular process | 0.0046 | 86 |
| GO:0031667~response to nutrient levels | 0.0048 | 9 |
| GO:0048856~anatomical structure development | 0.0049 | 33 |
| GO:0042598~vesicular fraction | 0.0050 | 9 |
| GO:0005622~intracellular | 0.0053 | 99 |
| GO:0043169~cation binding | 0.0053 | 38 |
| GO:0070271~protein complex biogenesis | 0.0054 | 11 |
| GO:0006461~protein complex assembly | 0.0054 | 11 |
| GO:0005829~cytosol | 0.0058 | 21 |
| GO:0051384~response to glucocorticoid stimulus | 0.0060 | 6 |
| GO:0005543~phospholipid binding | 0.0068 | 6 |
| GO:0005902~microvillus | 0.0069 | 4 |
| GO:0032787~monocarboxylic acid metabolic process | 0.0074 | 9 |
| GO:0031960~response to corticosteroid stimulus | 0.0076 | 6 |
| GO:0055065~metal ion homeostasis | 0.0076 | 7 |
| GO:0046887~positive regulation of hormone secretion | 0.0079 | 4 |
| GO:0032868~response to insulin stimulus | 0.0080 | 6 |
| GO:0046883~regulation of hormone secretion | 0.0081 | 5 |
| GO:0014074~response to purine | 0.0083 | 3 |
| GO:0043227~membrane-bounded organelle | 0.0089 | 73 |
| GO:0007399~nervous system development | 0.0095 | 18 |
| GO:0009743~response to carbohydrate stimulus | 0.0099 | 5 |
| GO:0031090~organelle membrane | 0.0100 | 18 |
| GO:0005624~membrane fraction | 0.0100 | 15 |
| GO:0043005~neuron projection | 0.0103 | 11 |
| GO:0043436~oxoacid metabolic process | 0.0109 | 12 |
| GO:0019752~carboxylic acid metabolic process | 0.0109 | 12 |
| GO:0006082~organic acid metabolic process | 0.0112 | 12 |
| GO:0016817~hydrolase activity, acting on acid anhydrides | 0.0115 | 12 |
| GO:0005774~vacuolar membrane | 0.0121 | 4 |
| GO:0042562~hormone binding | 0.0121 | 4 |
| GO:0044463~cell projection part | 0.0123 | 8 |
| GO:0005509~calcium ion binding | 0.0124 | 13 |
| GO:0042180~cellular ketone metabolic process | 0.0124 | 12 |
| GO:0065007~biological regulation | 0.0124 | 64 |
| GO:0000267~cell fraction | 0.0128 | 18 |
| GO:0030005~cellular di-, tri-valent inorganic cation homeostasis | 0.0129 | 7 |
| GO:0046872~metal ion binding | 0.0133 | 36 |
| GO:0043231~intracellular membrane-bounded organelle | 0.0136 | 72 |
| GO:0032879~regulation of localization | 0.0137 | 13 |
| GO:0007275~multicellular organismal development | 0.0141 | 33 |
| GO:0042391~regulation of membrane potential | 0.0141 | 6 |
| GO:0005792~microsome | 0.0143 | 8 |
| GO:0042491~auditory receptor cell differentiation | 0.0147 | 3 |
| GO:0051412~response to corticosterone stimulus | 0.0147 | 3 |
| GO:0003013~circulatory system process | 0.0149 | 6 |
| GO:0008015~blood circulation | 0.0149 | 6 |

| | | |
|---|---|---|
| GO:0007204~elevation of cytosolic calcium ion concentration | 0.0151 | 5 |
| GO:0005626~insoluble fraction | 0.0154 | 15 |
| GO:0002376~immune system process | 0.0156 | 14 |
| GO:0006950~response to stress | 0.0158 | 22 |
| GO:0048511~rhythmic process | 0.0161 | 6 |
| GO:0055066~di-, tri-valent inorganic cation homeostasis | 0.0162 | 7 |
| GO:0043232~intracellular non-membrane-bounded organelle | 0.0163 | 28 |
| GO:0043228~non-membrane-bounded organelle | 0.0163 | 28 |
| GO:0016337~cell-cell adhesion | 0.0166 | 7 |
| GO:0019226~transmission of nerve impulse | 0.0166 | 8 |
| GO:0042327~positive regulation of phosphorylation | 0.0166 | 5 |
| GO:0044437~vacuolar part | 0.0173 | 4 |
| GO:0043434~response to peptide hormone stimulus | 0.0176 | 7 |
| GO:0005525~GTP binding | 0.0176 | 8 |
| GO:0051260~protein homooligomerization | 0.0177 | 5 |
| GO:0044430~cytoskeletal part | 0.0181 | 14 |
| GO:0010562~positive regulation of phosphorus metabolic process | 0.0183 | 5 |
| GO:0045937~positive regulation of phosphate metabolic process | 0.0183 | 5 |
| GO:0016043~cellular component organization | 0.0183 | 28 |
| GO:0005515~protein binding | 0.0185 | 73 |
| GO:0006874~cellular calcium ion homeostasis | 0.0186 | 6 |
| GO:0017111~nucleoside-triphosphatase activity | 0.0186 | 11 |
| GO:0006810~transport | 0.0189 | 30 |
| GO:0043679~nerve terminal | 0.0190 | 4 |
| GO:0048545~response to steroid hormone stimulus | 0.0190 | 8 |
| GO:0051179~localization | 0.0191 | 34 |
| GO:0042803~protein homodimerization activity | 0.0194 | 8 |
| GO:0030285~integral to synaptic vesicle membrane | 0.0194 | 2 |
| GO:0032502~developmental process | 0.0203 | 35 |
| GO:0055074~calcium ion homeostasis | 0.0204 | 6 |
| GO:0009725~response to hormone stimulus | 0.0206 | 11 |
| GO:0051385~response to mineralocorticoid stimulus | 0.0208 | 3 |
| GO:0051239~regulation of multicellular organismal process | 0.0209 | 17 |
| GO:0005764~lysosome | 0.0212 | 6 |
| GO:0000323~lytic vacuole | 0.0212 | 6 |
| GO:0048518~positive regulation of biological process | 0.0213 | 27 |
| GO:0051050~positive regulation of transport | 0.0213 | 7 |
| GO:0051234~establishment of localization | 0.0214 | 30 |
| GO:0006629~lipid metabolic process | 0.0217 | 13 |
| GO:0030003~cellular cation homeostasis | 0.0217 | 7 |
| GO:0051480~cytosolic calcium ion homeostasis | 0.0218 | 5 |
| GO:0019001~guanyl nucleotide binding | 0.0218 | 8 |
| GO:0032561~guanyl ribonucleotide binding | 0.0218 | 8 |
| GO:0031668~cellular response to extracellular stimulus | 0.0227 | 4 |
| GO:0016462~pyrophosphatase activity | 0.0237 | 11 |
| GO:0060113~inner ear receptor cell differentiation | 0.0242 | 3 |
| GO:0019228~regulation of action potential in neuron | 0.0245 | 4 |

| | | |
|---|---|---|
| GO:0016818~hydrolase activity, acting on acid anhydrides, in phosphorus-containing anhydrides | 0.0247 | 11 |
| GO:0051047~positive regulation of secretion | 0.0251 | 5 |
| GO:0010033~response to organic substance | 0.0253 | 16 |
| GO:0006875~cellular metal ion homeostasis | 0.0254 | 6 |
| GO:0030193~regulation of blood coagulation | 0.0259 | 3 |
| GO:0035091~phosphoinositide binding | 0.0263 | 4 |
| GO:0010627~regulation of protein kinase cascade | 0.0265 | 7 |
| GO:0051049~regulation of transport | 0.0278 | 10 |
| GO:0008361~regulation of cell size | 0.0293 | 6 |
| GO:0008585~female gonad development | 0.0295 | 4 |
| GO:0065003~macromolecular complex assembly | 0.0304 | 11 |
| GO:0006631~fatty acid metabolic process | 0.0311 | 6 |
| GO:0009628~response to abiotic stimulus | 0.0329 | 9 |
| GO:0042493~response to drug | 0.0331 | 8 |
| GO:0034284~response to monosaccharide stimulus | 0.0339 | 4 |
| GO:0009746~response to hexose stimulus | 0.0339 | 4 |
| GO:0022607~cellular component assembly | 0.0339 | 13 |
| GO:0045177~apical part of cell | 0.0349 | 6 |
| GO:0046545~development of primary female sexual characteristics | 0.0350 | 4 |
| GO:0042490~mechanoreceptor differentiation | 0.0356 | 3 |
| GO:0050818~regulation of coagulation | 0.0356 | 3 |
| GO:0016787~hydrolase activity | 0.0359 | 25 |
| GO:0010044~response to aluminum ion | 0.0367 | 2 |
| GO:0022602~ovulation cycle process | 0.0374 | 4 |
| GO:0019955~cytokine binding | 0.0384 | 4 |
| GO:0022008~neurogenesis | 0.0387 | 12 |
| GO:0005856~cytoskeleton | 0.0404 | 16 |
| GO:0001508~regulation of action potential | 0.0410 | 4 |
| GO:0009719~response to endogenous stimulus | 0.0411 | 11 |
| GO:0004957~prostaglandin E receptor activity | 0.0420 | 2 |
| GO:0033267~axon part | 0.0420 | 4 |
| GO:0016324~apical plasma membrane | 0.0422 | 5 |
| GO:0046660~female sex differentiation | 0.0423 | 4 |
| GO:0007165~signal transduction | 0.0423 | 23 |
| GO:0043933~macromolecular complex subunit organization | 0.0423 | 11 |
| GO:0030424~axon | 0.0438 | 6 |
| GO:0019229~regulation of vasoconstriction | 0.0441 | 3 |
| GO:0031982~vesicle | 0.0444 | 12 |
| GO:0010959~regulation of metal ion transport | 0.0449 | 4 |
| GO:0042127~regulation of cell proliferation | 0.0456 | 12 |
| GO:0002925~positive regulation of humoral immune response mediated by circulating immunoglobulin | 0.0457 | 2 |
| GO:0002922~positive regulation of humoral immune response | 0.0457 | 2 |
| GO:0032847~regulation of cellular pH reduction | 0.0457 | 2 |
| GO:0042698~ovulation cycle | 0.0462 | 4 |
| GO:0032846~positive regulation of homeostatic process | 0.0464 | 3 |
| GO:0044093~positive regulation of molecular function | 0.0470 | 10 |

| | | |
|---|---|---|
| GO:0051259~protein oligomerization | 0.0473 | 6 |
| GO:0045792~negative regulation of cell size | 0.0475 | 4 |

**Supplementary Table 12: List of genes similarly regulated in the mPFC transcriptome of adult PNFlx and JFlx animals.**

Analysis of the mPFC transcriptome of adult PNFlx and JFlx animals revealed minimal overlap with very few genes regulated in common in the two models. Given in the table is a list of genes found similarly regulated in PNFlx and JFlx animals.

| Common upregulated genes in PNFlx and JFlx mPFC microarray | | |
|---|---|---|
| **Systematic Name** | **Gene Name** | **Product** |
| NM_012507 | Atp1b2 | Na+/K+ -ATPase beta 2 subunit |
| XM_341133 | Lamc1 | similar to Laminin gamma-1 chain precursor (Laminin B2 chain) |
| **Common downregulated genes in PNFlx and JFlx mPFC microarray** | | |
| **Systematic Name** | **Gene Name** | **Product** |
| XM_343146 | Silv_predicted | similar to silver |
| NM_001009275 | RGD1310224 | hypothetical protein LOC291076 |
